# Cell cycle dysregulation of globally important SAR11 bacteria resulting from environmental perturbation

**DOI:** 10.1101/2025.06.25.661606

**Authors:** Chuankai Cheng, Brittany D. Bennett, Pratixa Savalia, Hasti Asrari, Carmen Biel, Kate A. Evans, Rui Tang, J. Cameron Thrash

## Abstract

Genome streamlining is hypothesized to occur in bacteria as an adaptation to resource-limited environments, but can result in gene losses affecting fundamental aspects of cellular physiology. The most abundant marine microorganisms, SAR11 (order *Pelagibacterales*), exhibit canonical genome streamlining, but the consequences of this genotype on core cellular processes like cell division remain unexplored. Here, analysis of 470 SAR11 genomes revealed widespread absence of key cell cycle control genes. Growth experiments demonstrated that although SAR11 bacteria maintain a normal cell cycle under oligotrophic conditions, they exhibit growth inhibition and aneuploidy when exposed to nutrient enrichment, carbon source shifts, or temperature stress. Detailed growth measurements and antibiotic inhibition experiments showed that these phenotypes resulted from cell division disruption with continuing DNA replication, leading to heterogeneous subpopulations of normal and polyploid cells. This vulnerability raises questions about microbial genome evolution and the evolutionary trade-offs between adaptations to stable nutrient-limited conditions and physiological resilience.

## Introduction

Some microbial lineages have undergone significant genome reduction over evolutionary time, resulting in naturally evolved cells that reveal fundamental principles of cell biology under reduced genomic complexity^1,2^. Among them, SAR11 bacteria stand out as an exceptional model. With genome sizes typically around 1.2-1.3 Mbp—among the smallest of any free-living cells— these bacteria dominate marine ecosystems, comprising up to 40% of bacterial cells in surface seawater, and are canonical oligotrophic taxa adapted to low nutrient environments^3–5^. Their small genomes exhibit patterns collectively known as genome streamlining, such as high coding densities, few paralogs or pseudogenes, and enzyme promiscuity^6–9^, that has been proposed to help them reduce nutritional requirements to thrive in extremely oligotrophic open ocean habitats^1,5,10^. Unlike synthetic minimal cells engineered in laboratory settings^11–13^, SAR11 representatives are compelling “natural minimal cells” for investigating how genome reduction relates to increased fitness in the environment.

Despite their ecological success, SAR11 populations have numerous ostensibly disadvantageous traits and puzzling ecological patterns that suggest potential physiological vulnerabilities. Their streamlined genomes have decreased regulatory capacity^1,14,15^ and metabolic consequences like a dependence on reduced sulfur sources^16^ and amino acid and vitamin auxotrophies^17,18^. SAR11 do respond to phytoplankton blooms^19^, but die in co-culture with the dominant phytoplankton *Prochlorococcus* when it reaches stationary phase^20^ (unlike copiotrophic bacteria), and exhibit cell-cycle arrest when pyruvate concentrations are low relative to those of glycine^18^. Their small genomes also only contain single copies of ribosomal RNA genes that may result in limited translational power^21,22^, thus contributing to their obligate slow growth physiology, with cells typically dividing only around once per day under optimal conditions both *in situ* and in culture^18,23–27^.

These observations raise questions about the evolutionary trade-offs between genome streamlining and fitness. Importantly, they point to possible consequences of genome reduction in SAR11 on essential cellular processes like the cell cycle—the tightly regulated progression of cellular growth, DNA replication, and division^28^. Here we examined how SAR11 bacteria coordinate cell division and DNA replication in the context of a reduced genome, and what physiological constraints SAR11 responses impose on their ability to acclimate to environmental change.

## Results

### Most SAR11 lack canonical cell cycle regulation genes

We analyzed a previous pangenome of SAR11 with 470 representatives^27^, as well as the pangenome of 120 SAR11 representatives on UniProt, and found that SAR11 lacks many of the canonical genes that coordinate cell division and DNA replication in *Escherichia coli*^29^, *Bacillus subtilis*^30^, *Caulobacter crescentus*^31^, *Cereibacter sphaeroides*^32^, and the synthetic minimal cell JCVI-syn3^33^ **(Fig. 1, Supplementary Table Tab 1-16, Supplementary Text)**. Several regulators of replication initiation were nearly or completely absent across SAR11, including *seqA*^34^, *dam*^35^, and *chpT*^36^. Although *seqA* was largely absent from Alphaproteobacteria, *dam* and *chpT* were widespread within the Alphaproteobacteria so their absence in SAR11 is unusual. Genes essential for divisome assembly and positioning were mostly missing in SAR11, such as *zipA*^37^ (mostly gammaproteobacterial) and the widely distributed *ftsE*^38^ and *mipZ*^39^. Most or all SAR11 genomes also lacked *minC* and *minD*, key regulators involved in chromosome segregation^30^, *sepF* and *lemA*, both essential for maintaining uniform cell sizes in the minimal cell JCVI-syn3A^33^, and the Caulobacterales polarity factor *tipN*^40^. These conspicuous gene absences led us to question how SAR11 thrives globally without such important cell cycle regulators. *B. subtilis minCD* knockouts had relatively few phenotypic consequences in a nutrient-limited medium whereby 89% of mutant cells formed normal Z-rings versus only 23% in rich medium^30^. Similarly, *E. coli seqA*, *topA*, and *dam* mutants also showed exaggerated cell cycle disruption in media with greater nutrient content compared to those with less^29^. In *C. crescentus*, deletion of *tipN* produced little difference in growth or cell size under minimal medium, but significant differences in rich medium^40^. Together, these findings suggest that if losing these cell cycle regulation genes has relatively minimal phenotypic consequences under nutrient-limited growth, selective pressure to maintain them might be relaxed in the consistently nutrient-limited oceans where SAR11 dominates.

**Figure 1.**
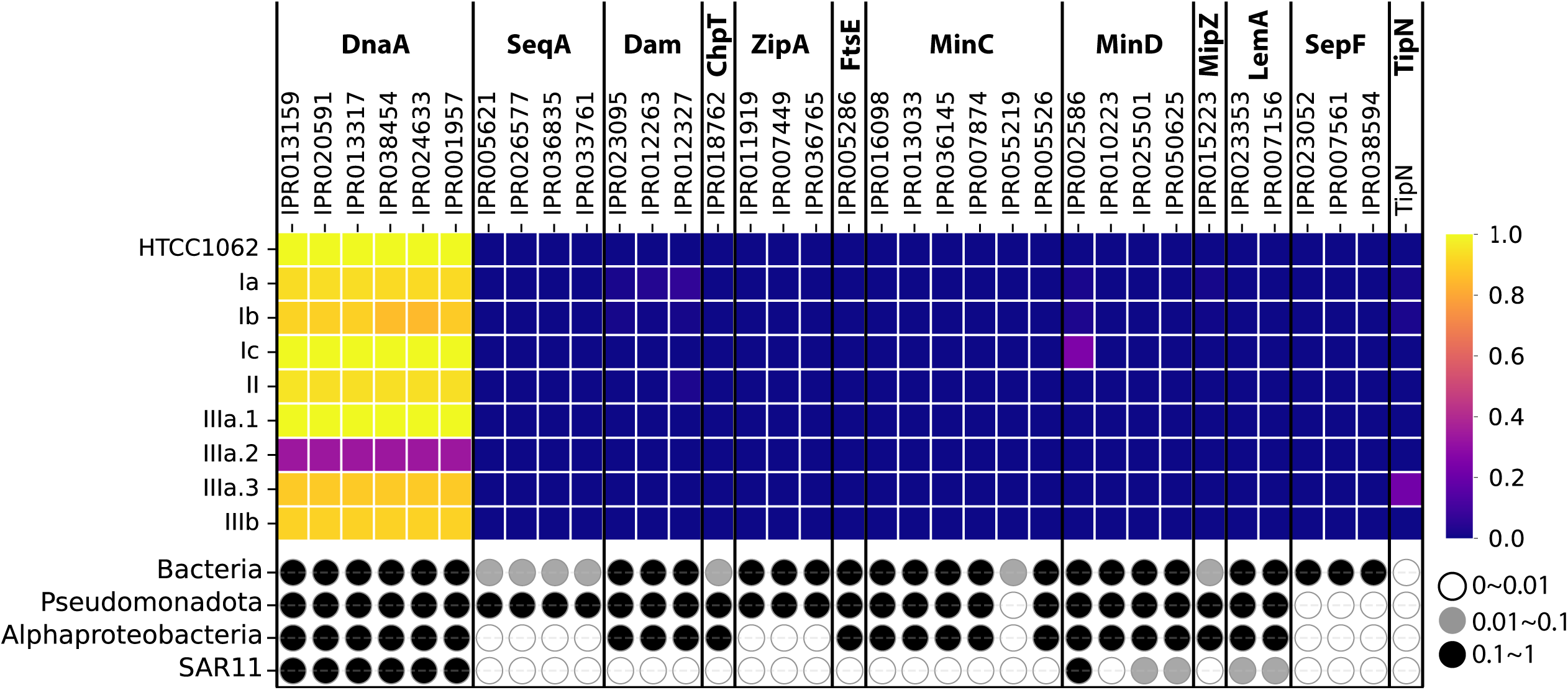
Distributions of canonical cell cycle regulatory proteins across SAR11 subclades (top) and other bacteria (bottom). The top panel is a heatmap showing the distribution of proteins within HTCC1062 and the SAR11 subclades (Ia, Ib, Ic, II, IIIa.1, IIIa.2, IIIa.3, IIIb) based on our pangenomic analysis of 470 representatives. Each column represents a different InterProScan ID for the labeled protein. Color represents the fraction of genomes with the protein present: yellow (1.0) = present in all genomes, purple (0.2-0.4) = present in a minority of genomes, dark blue (0.0) = absent from all genomes. DnaA^96,97^ serves as a positive reference expected to be present in most genomes. The bottom panel shows the taxonomic distribution of each protein across broader bacterial groups (Bacteria, Pseudomonadota, Alphaproteobacteria) in the context of the overall SAR11 clade based on UniProt searches. The the shade of the circle indicates the relative distribution of a protein in UniProt normalized to DnaA for each taxonomic group: black circles (0.1-1) indicate proteins present in 10%-100% of genomes compared to DnaA, grey circles (0.01-0.1) indicate proteins present at between 1-10% that of DnaA, and empty circles (0-0.01) indicate proteins that are absent or mostly absent, occurring in less than 1% of genomes compared to DnaA. Detailed accounting of protein distributions across individual SAR11 genomes or bacterial groups is provided in **Supplementary Table Tab 1-3**.

### Nutrient enrichment drives growth inhibition and aneuploidy

To test whether the absence of cell cycle regulation genes in SAR11 would result in deleterious growth phenotypes, we grew the SAR11 representative *Pelagibacter ubiqueversans*^41^ (previously, *Candidatus* Pelagibacter ubique^42^) strain HTCC1062 in increasingly rich media by proportionally varying the concentrations of pyruvate, glycine, ammonium, phosphate, and vitamins **(Fig. 2, Supplementary Table Tab 17)**. Rather than monotonically increasing and asymptoting, HTCC1062 growth showed a “bell-shaped” phenotype. Growth yields increased as nutrient concentrations rose from 0.5x to 3x, followed by significant decreases in both rate and yield under higher nutrient conditions (8x and 10x) **(Fig. 2a-c)**. This “bell-shaped” phenotype aligns with SAR11’s oligotrophic distribution where these organisms thrive in nutrient-limited environments but are inhibited in rich nutrient conditions.

**Figure 2.**
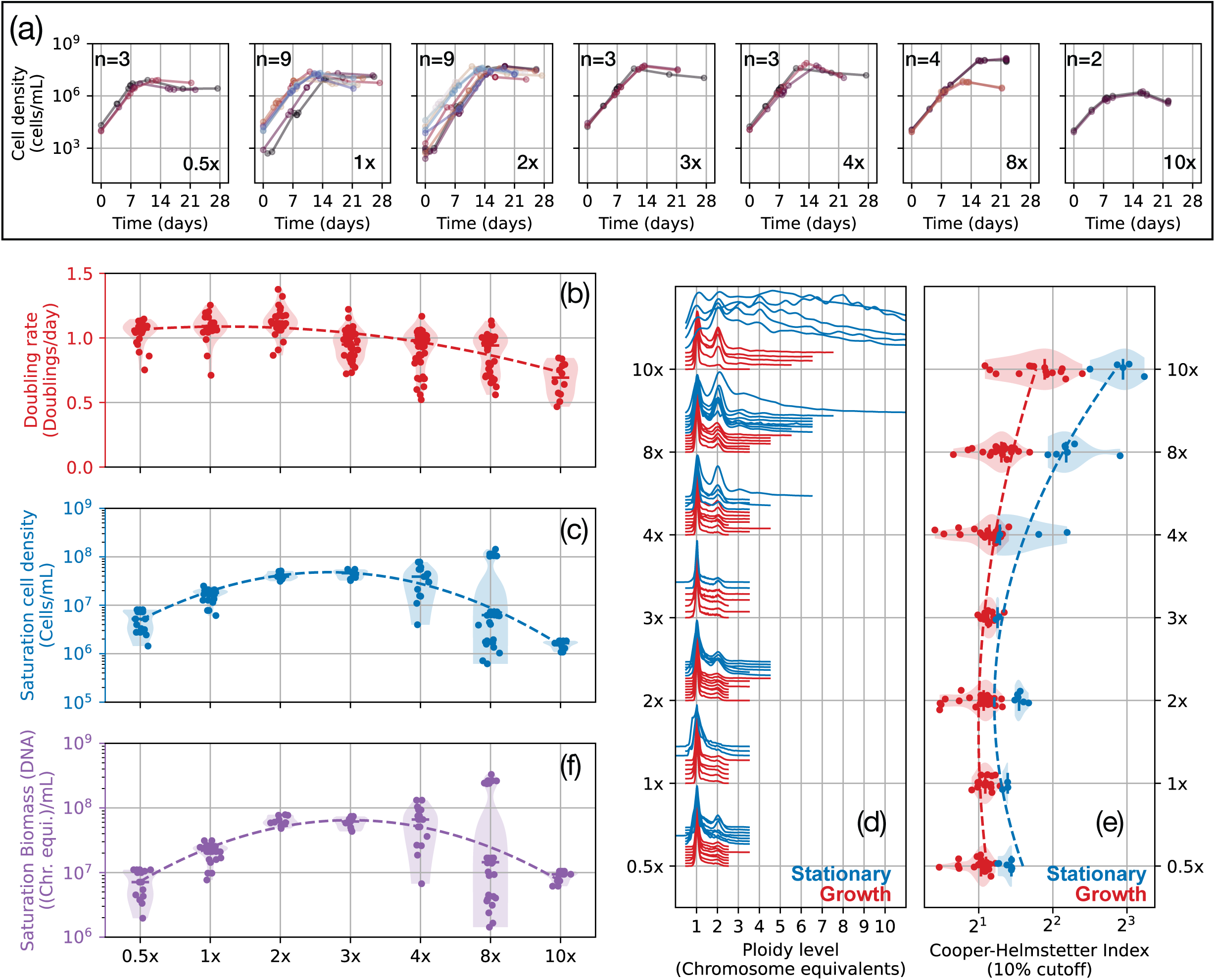
SAR11 growth response to nutrient enrichment. **(a)** Growth curves of SAR11 strain HTCC1062 under increasing nutrient concentrations (0.5x to 10x). The number of replicates for each growth curve are specified within each plot. **(b)** Instantaneous doubling rates across nutrient concentrations. **(c)** Saturation cell density across nutrient concentrations. **(d)** Flow cytometry histograms of cellular DNA content showing ploidy distributions during exponential (red) and stationary (blue) growth phases across nutrient concentrations. **(e)** Cooper-Helmstetter Index (ratio of maximum to minimum ploidy) across nutrient concentrations. Dashed lines indicate trends. **(f)** Saturation cellular DNA concentration (chromosome equivalents per mL) across nutrient concentrations. Data for this figure is available at **Supplementary Table Tabs 19, 20**.

Analysis of ploidy levels revealed distinct cell cycle patterns across nutrient conditions. Under limited nutrients (0.5x–3x), HTCC1062 cultures maintained normal ploidy (1-2 chromosomes) **(Fig. 2d)**, indicating a non-overlapping cell cycle where DNA replication completes before the next round begins **(Supplementary Text, Fig. S1)**. According to the Cooper-Helmstetter model^43^, in steady-state bacterial growth^44^ the ratio of cells at maximum ploidy to minimum ploidy—which we term the Cooper-Helmstetter Index —is approximately 2:1^45^. HTCC1062 maintained this ∼2:1 ratio under 0.5x-3x nutrient conditions, reflecting normal cell cycle progression. In contrast, under richer conditions (4x–10x nutrients), we observed HTCC1062 subpopulations with more than two chromosomes and Cooper-Helmstetter Indices >> 2, indicating aneuploidy **(Fig. 2e)**. This differs from the typical ‘overlapping cell cycle’ of copiotrophs, where despite multiple replication forks, cells maintain a Cooper-Helmstetter Index ∼2 **(Supplementary Text, Fig. S1-5)**.

Since we observed more cells with higher ploidy levels in richer nutrient conditions, we wanted to determine whether this reflected a redistribution of DNA content among a smaller population or instead represented changes in total DNA production. We quantified total DNA concentrations by calculating the product of cell density and mean cellular ploidy level. We found that the total DNA concentration also decreased significantly under 10x nutrient conditions **(Fig. 2f)**, demonstrating that, although some cells contained more chromosomes, the overall DNA quantity in the culture was substantially reduced due to lower cell numbers and dysregulated DNA replication. Thus, while individual cells accumulated excess chromosomes, the population as a whole had less DNA and suffered impaired growth.

### Specific factors disrupting the SAR11 cell cycle

Having established that elevated nutrient concentrations induced growth inhibition and aneuploidy in SAR11 strain HTCC1062, we next investigated which specific environmental factors triggered these cell cycle disruptions. Since our initial experiment proportionally varied multiple nutrients simultaneously (pyruvate, glycine, ammonium, phosphate, and vitamins, **Supplementary Table Tab 17**), we sought to determine whether particular nutrients or combinations were responsible for the observed effects, and whether other environmental parameters besides nutrients could elicit similar responses.

HTCC1062 is a glycine auxotroph^17^ and low pyruvate:glycine ratios inhibit cell cycle progression, resulting in an accumulation of cells with diploid DNA content^18^. Since this diploid phenotype was similar to our observations of aneuploidy based on proportional increases of all nutrients, we examined the pyruvate:glycine ratio effects in detail. Through re-analysis of the data from Carini *et al.* 2013^18^ and application of the Cooper-Helmstetter Index, we found indicators of polyploidy beyond two chromosome equivalents and filamentous morphologies that were not emphasized in the original study **(Supplementary Text, Fig. S6)**. In our own experiments, we confirmed that pyruvate:glycine ratios between 5:1 and 2:1 supported stable steady-state growth with normal ploidy (Cooper-Helmstetter Index ∼2), whereas ratios at or below 1:1 induced aneuploidy (Cooper-Helmstetter Index >> 2; **Fig. 3a, Extended Data Fig. 1**). The concentration of these compounds was less important than their ratio - by scaling pyruvate and glycine proportionally at 5:1, we could modulate saturation cell density without affecting growth rates or ploidy levels **(Fig. 3b, Extended Data Fig. 2a)**, thus improving growth yields 100-fold from the original AMS1 medium recipe which was most commonly used for culturing HTCC1062^18^ **(Extended Data Fig. 3, Supplementary Table Tab 18)**.

**Figure 3.**
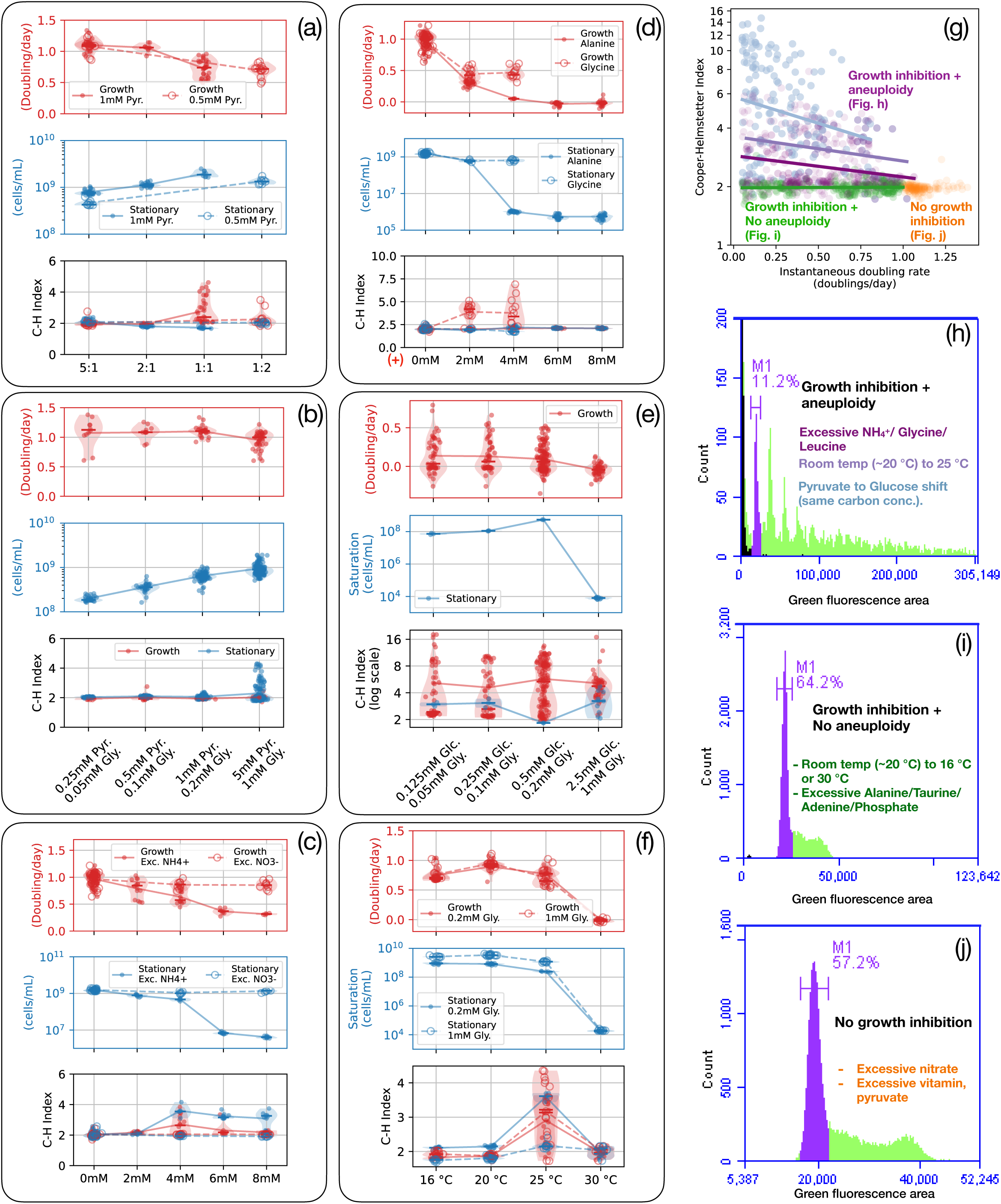
Different factors triggering cell cycle dysregulation in SAR11. **(a-f)** SAR11 growth rates and yields (top panels) with corresponding Cooper-Helmstetter Index measurements (bottom panels) under various conditions. **(a)** Effect of pyruvate-to-glycine ratios, with cultures transferred from an optimal 5:1 ratio to the experimental ratios. **(b)** Proportional scaling of pyruvate and glycine concentrations while maintaining an optimal 5:1 ratio, with cultures transferred from a 1 mM pyruvate/0.2 mM glycine baseline. **(c)** Effect of inorganic nitrogen sources: ammonium (NH_₄ ⁺_) and nitrate (NO_₃_^⁻^) at 0–8 mM, with cultures transferred from 5 mM pyruvate/1 mM glycine without additional inorganic nitrogen. **(d)** Differential effects of organic nitrogen sources: glycine versus alanine. Red “+” indicates additional compounds added to base media, not absolute concentrations. **(e)** Carbon source shifts from pyruvate to glucose while maintaining equivalent carbon concentrations. **(f)** Temperature responses (16–30°C) under different glycine concentrations, with cultures transferred from room temperature. **(g)** Relationship between instantaneous doubling rate and Cooper-Helmstetter Index revealing three distinct response categories: *i)* growth inhibition with aneuploidy, *ii)* growth inhibition without aneuploidy, and *iii)* normal growth. **(h-j)** Flow cytometry profiles of cellular DNA content under representative conditions from each response category. Additional experimental data are shown in Fig. 4 and **Extended Data Figs. 1, 2**, and **4**. Detailed nutrient conditions are provided in **Supplementary Table Tab 17**.

Although a pyruvate:glycine ratio below 1:1 led to growth defects, this did not explain the observation of polyploidy in our bulk nutrient experiments, because there we maintained the proportions of all growth compounds. We therefore tested other medium components for similar cell cycle dysregulation effects with increased concentrations. We found that excessive ammonium leads to both growth inhibition and aneuploidy (**Fig. 3c**, **Fig. 4**), whereas increased vitamin concentrations neither inhibited growth nor disrupted normal cell cycle progression **(Extended Data Fig. 2bc)**, and excessive phosphate (≥ 500 μM) did not affect the growth rate or cause aneuploidy, but significantly reduced saturation cell density **(Extended Data Fig. 4)**.

**Figure 4.**
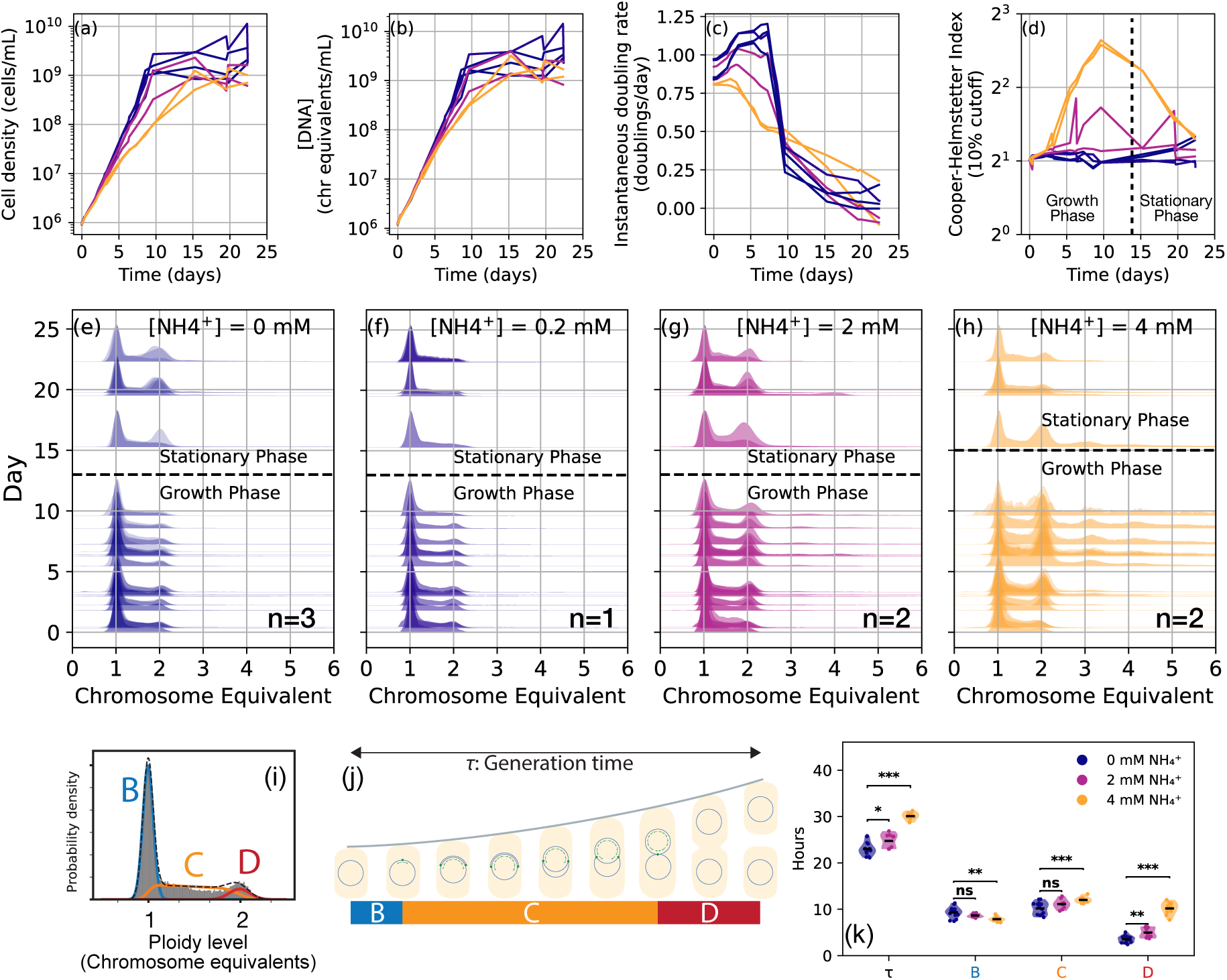
High-resolution temporal analysis of excessive ammonium effects on SAR11 cell cycle regulation. **(a-d)** Time course measurements of SAR11 growth parameters under varying ammonium concentrations (0-4 mM): (a) cell density, (b) total DNA, (c) instantaneous doubling rate, and (d) Cooper-Helmstetter Index. **(e-h)** Flow cytometry histograms showing DNA content distributions over time under different ammonium concentrations: (e) 0 mM, (f) 0.2 mM, (g) 2 mM, and (h) 4 mM NH_4_^+^. Detailed nutrient concentrations are provided in **Supplementary Table Tab 16**. Dashed lines separate exponential and stationary growth phases. **(i)** Theoretical ploidy distribution showing maximum likelihood estimation (see **Methods**) of B (between cell birth and DNA replication initiation), C (DNA replication), and D (between replication completion and cell division) periods (in colored contours) fitted to experimental measurements (gray histogram). **(j)** Schematic representation of bacterial cell cycle with B, C, and D periods across one generation (T). **(k)** Ammonium stress extends SAR11 generation time and delays cell division. Cell cycle period durations under increasing ammonium concentrations, estimated from ploidy level distributions collected during the first five days of growth. Statistical comparisons between treatments were added (* p < 0.05, ** p < 0.01, *** p < 0.001; Mann-Whitney U two-sided tests comparing 0 mM vs 2 mM and 0 mM vs 4 mM NH_4_^+^h). For the specific p-values, T: p=0.012266 (0 mM vs. 2 mM), p=0.000036 (0 mM vs. 4 mM); B: p=0.056377 (0 mM vs. 2 mM), p=0.006037 (0 mM vs. 4 mM); C: p=0.094999 (0 mM vs. 2 mM), p=0.000646 (0 mM vs. 4 mM); D: p=0.002869 (0 mM vs. 2 mM), p=0.000036 (0 mM vs. 4 mM).

Since glycine and ammonium could disrupt the cell cycle and cause growth inhibition, we tested whether other nitrogen sources would produce the same phenotype. Nitrate had no effect (as expected because HTCC1062 has no predicted nitrate assimilation genes) (**Fig. 3c**). Similarly to excess glycine and ammonium, leucine caused both growth inhibition and aneuploidy (**Extended Data Fig. 2d**), but supplemental alanine (**Fig. 3d, Extended Data Fig. 2e**), taurine **(Extended Data Fig. 2f)**, and adenine (**Extended Data Fig. 2g**) inhibited population growth without inducing polyploidy. These differential responses highlight that cell cycle disruption depends on the specific nitrogen source metabolized rather than total nitrogen availability.

We also tested the effect of changing the carbon source from pyruvate to glucose while maintaining equivalent carbon concentrations. Despite this stoichiometric consistency, we observed significant cell cycle disruption, indicating that the specific carbon source, rather than just the carbon-to-nitrogen ratio, can affect cell cycle regulation (**Fig. 3e, Extended Data Fig. 2h**).

In addition to various chemical perturbations, we tested the effect of different temperatures. At lower temperatures (16-20°C), HTCC1062 maintained normal ploidy levels even with the 1:1 pyruvate to glycine ratio (**Fig. 3f, Extended Data Fig. 2i**) that caused cell cycle disruption under our default room temperature (fluctuating around 20°C - **Extended Data Fig. 5**). As temperatures increased to 25°C, aneuploidy occurred even with a 5:1 pyruvate:glycine ratio, but was more severe with a 1:1 pyruvate to glycine ratio, while at 30°C, growth was completely inhibited regardless of nutrient concentrations (**Fig. 3f, Extended Data Fig. 2i**).

In summary, we identified three distinct patterns of SAR11 cell cycle dysregulation (**Fig. 3g-i**): *i)* growth inhibition with aneuploidy occurred in response to glucose substitution, temperature elevation to 25°C, and excessive ammonium, glycine, or leucine (**Fig. 3h**); *ii)* growth inhibition without aneuploidy was observed during temperature shifts to 30°C, and with excessive alanine, taurine, adenine, or phosphate (**Fig. 3i**); and *iii)* no growth inhibition with additional nitrate, vitamins, or pyruvate (**Fig. 3j**). Notably, when comparing the Cooper-Helmstetter Index to the instantaneous doubling rate, the data forms a triangular cluster pattern, revealing that growth inhibition is frequently associated with cell cycle dysregulation (**Fig. 3g**). Cultures maintaining optimal growth rates consistently exhibited normal ploidy (Cooper-Helmstetter Index ∼2), while growth-inhibited cultures often, but not always, displayed aberrant ploidy levels.

### Onset and fate of aneuploidy under excessive ammonium

We next investigated the progression and mechanisms of aneuploidy induction. Among all the aneuploidy inducing conditions, we chose excessive ammonium for deeper investigation because ammonium is a major and universal inorganic nitrogen source for marine microorganisms^46^ and excretion of ammonium can be induced by phytoplankton-bacteria interactions^47^. Also, the simple chemical composition of ammonium minimized the potential confounding factors compared to other aneuploidy triggering conditions, such as complex growth dynamics with extended lag times and multiple growth phases (pyruvate to glucose shifts; **Extended Data Fig. 2h**), co-involvement of carbon metabolism changes and additional nitrogen sources (amino acid supplements), or general effects on enzymes through temperature shifts^48^. We therefore conducted high-resolution sampling of HTCC1062 cultures exposed to varying ammonium concentrations (0–4 mM) **(Fig. 4)** and observed significant growth inhibition in the high ammonium treatments compared to the 0 mM ammonium samples **(Fig. 4a–c)** with simultaneous increases in the Cooper-Helmstetter Index **(Fig. 4d)** and diploid accumulation **(Fig. 4e–h)**.

During the first five days, where the growth curves under different ammonium concentrations started diverging, the population doubling time *τ* increased from 23.02 ± 1.39 hours for the 0 mM NH_4_^+^ treatment to 30.08 ± 0.81 hours in the 4 mM NH_4_^+^ treatment **(Fig. 4k)**. During this initial period of growth divergence, when the ploidy level distributions were maintained as mostly monoploids and diploids, we could also quantify cell cycle period durations using the Cooper-Helmstetter model (**Methods, Supplementary Text, Fig. S7**)^45^. In non-overlapping cell cycle progression, these periods are: B (between cell birth and DNA replication initiation), C (DNA replication), and D (between replication completion and cell division) **(Fig. 4ij)**. D period experienced the most dramatic extension under high ammonium, increasing from 3.53 ± 0.70 hours at 0 mM to 10.19 ± 1.26 hours at 4 mM NH_4_^+^ **(Fig. 4k)**, indicating delayed cell division after DNA replication completed. The C period showed modest increases (from 10.19 ± 1.28 to 12.02 ± 0.60 hours), while the B period slightly decreased with elevated ammonium (from 9.31 ± 1.11 to 7.86 ± 0.49 hours), suggesting earlier DNA replication initiation. Thus, excessive ammonium primarily impacted post-replication processes, with aneuploidy resulting from division delays coupled to continued chromosome replication.

Polyploidy appeared after day 5 in samples with high ammonium concentrations **(Fig. 4gh)**. Specifically, we observed that a third chromosome equivalent (triploidy) occurred first, indicating an imbalance in DNA replication initiation whereby only one of the chromosomes in diploids was duplicated. The appearance of odd number chromosome equivalents was consistent with the lack of regulatory genes controlling DNA replication initiation **(Supplementary Text, Fig. S4)**, and mirrored that of *E. coli* mutations in the same genes^29^. Because of aneuploidy (Cooper-Helmstetter Index >> 2), after day 5 we could no longer estimate the durations of B, C, and D periods based on the Cooper-Helmstetter model, so we used cell cycle-disrupting antibiotics (in antibiotic ‘run-off’ experiments) to validate our measurements of cell cycle periods under ammonium stress. We used cephalexin^49^, which inhibits cell division without affecting DNA replication, alongside rifampin and chloramphenicol^50^, which inhibit transcription and translation, both required for new rounds of DNA replication initiation. By monitoring the ploidy distributions of ammonium-treated samples after treatment with these antibiotics, we confirmed that excessive ammonium causes division delays while allowing continued DNA replication **(Supplementary Text, Fig. S8)**. Furthermore, samples treated with only cephalexin (cell division inhibition) confirmed that triploids developed from diploids, indicating that diploids conduct DNA replication on only one of the chromosomes. These antibiotic treatments also corroborated our B, C, and D period estimations from above and provided an orthogonal confirmation of the mechanisms underlying the cell cycle dysregulation we observed in SAR11 **(Supplementary Text, Fig. S8)**.

Using fluorescence microscopy, we observed both filamentous and normal rod-shaped HTCC1062 cells in our samples with polyploidy under excessive ammonium **(Extended Data Fig. 6a–f)**. This pleomorphic phenotype is similar to the polymorphism observed in HTCC1062 under low pyruvate:glycine ratios^18^ and previous genetic studies in *E. coli*^51^, *C. crescentus*^31^, and the minimal synthetic cell JCVI-Syn3^33^ where cell cycle regulation genes were disrupted. Dynamic light scattering measurements confirmed a trend toward increased mean cell size with higher ploidy levels, corroborating the generation of filamentous cells with larger sizes (**Extended Data Fig. 6g–j**). As the cultures reached stationary phase, we observed a decrease in the Cooper-Helmstetter Index, indicating that the frequency of polyploids decreased over time, leaving cells predominantly with 1-2 chromosome equivalents **(Fig. 4gh)**. Flow cytometric analysis revealed increased background fluorescence in high ammonium samples during the same growth stage **(Extended Data Fig. 7a–e)**, similar to patterns observed when we subjected actively growing cells to a lysis buffer, suggesting selective lysis of polyploid cells in stationary phase **(Extended Data Fig. 7f–i)**.

### Growth inhibition results from subpopulation heterogeneity

Our high-resolution temporal analysis revealed that excessive ammonium created a mixed population of euploid and aneuploid cells. This heterogeneity, with co-occurrence of both filamentous and normal rod-shaped cells, indicated that not all cells responded identically to ammonium stress. We also observed that polyploid cells were more prone to lysis as cultures entered stationary phase, suggesting that aneuploidy represents a physiological dead-end rather than an adaptive strategy for SAR11.

Inspired by previous studies on cell cycle regulation in bacteria with disrupted divisome genes^30^, we hypothesized that excessive ammonium creates two distinct subpopulations: 1) a normally dividing subset and 2) a non-dividing subset where DNA replication continues but cell division fails. We propose that the absence of the key cell cycle regulation genes discussed above **(Fig. 1, Supplementary Text)** provides the mechanistic basis for this subpopulation heterogeneity.

To test our hypothesis, we applied a quantitative subpopulation growth model that assumes population growth results from two subpopulations: one actively dividing and one non-dividing. The model predicts overall population growth rate using the active subpopulation growth rate and the fraction of actively dividing cells (inferred from the proportion of cells experiencing aneuploidy) as inputs **(Fig. 5ab)**. This model successfully predicted population doubling rates under excessive ammonium conditions **(Fig. 5c)**, providing strong quantitative validation that the population growth slowdown stems directly from the emergence of a subpopulation of chromosome-replicating but non-dividing cells **(Fig. 5d)**.

**Figure 5.**
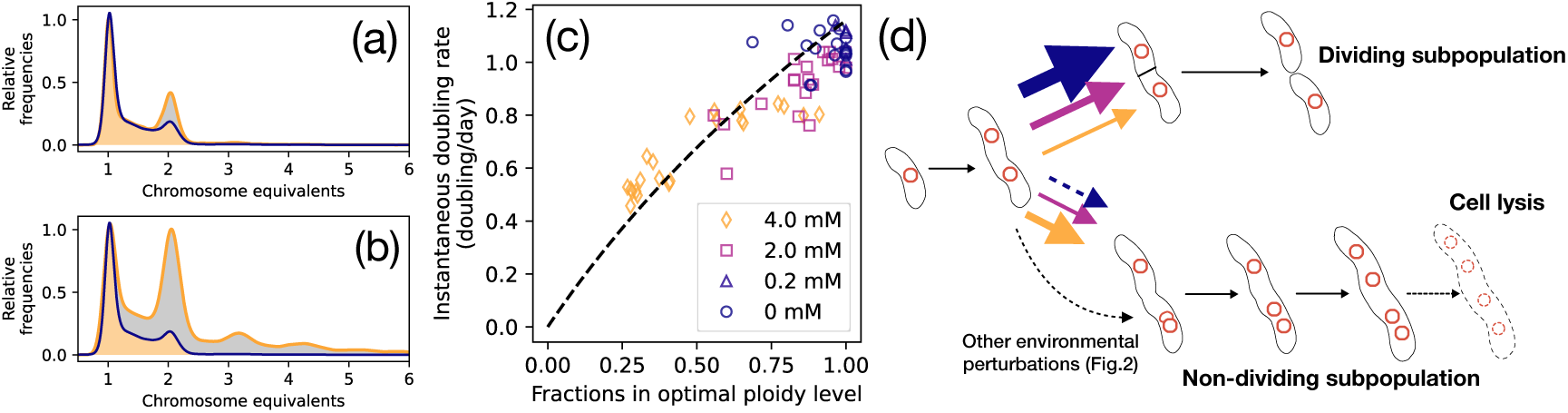
Subpopulation model quantitatively validates the link between cell cycle heterogeneity and growth inhibition. **(a-b)** Ploidy level distributions comparing optimal growth conditions (0 mM NH_4_^+^, blue contour) with high ammonium conditions (4 mM NH_4_^+^, orange contour) at **(a)** day 3 and **(b)** day 9. Area under the orange contours is shaded either gray or orange: the orange areas are exactly covered under the blue contour, indicating the subpopulation of the orange contour that is under optimal ploidy level, with the fraction of *Φ* in the subpopulation model, the gray areas represent the subpopulations that are under aneuploidy defined as *1-Φ* (see **Methods**). **(c)** Relationship between population instantaneous doubling rate *λ_p_* and fraction of cells with optimal ploidy levels *Φ* across different ammonium concentrations. Dashed curve represents predictions from the active subpopulation model, *λ_p_ = λ⋅ log2(1+Φ),* not a fit to the data (see **Methods**). **(d)** Conceptual model showing how excessive nitrogen creates heterogeneous populations: cells maintaining optimal ploidy continue dividing at normal rates while aneuploid cells fail to divide, eventually leading to cell lysis. Solid arrow colors correspond to ammonium concentrations, and the greater the ammonium, the more cells become aneuploid. The dashed black arrow reflects the fact that many other environmental perturbations also lead to aneuploidy.

### Cell cycle vulnerability is conserved in divergent SAR11

Since we found a systemic lack of cell cycle regulatory genes across the SAR11 clade (**Fig. 1**), we conducted parallel experiments on the phylogenetically distant freshwater SAR11 (LD12) strain LSUCC0530^26^ under various temperatures and ammonium chloride concentrations. The freshwater LD12/IIIb subclade is sister to the brackish-water adapted subclade IIIa, and both of these lineages likely diverged from a common ancestor with other SAR11 hundreds of millions of years ago^26,27,52^. Notably, LSUCC0530 and many other SAR11 subclade III strains also lack cell cycle regulatory genes, but possess some of the P-II nitrogen regulatory genes that are absent in HTCC1062 **(Supplementary Table Tab 3)**^26,27,53^. Comparing the growth responses in LSUCC0530 therefore afforded us an opportunity to assess whether the cell cycle dysregulation phenotype spans highly divergent SAR11 membership and if P-II nitrogen-sensing genes affect cell cycle responses in SAR11 under elevated nitrogen conditions.

We found that LSUCC0530 still exhibited significant cell cycle vulnerabilities **(Fig. 6)**. At 16°C, LSUCC0530 showed growth inhibition at higher ammonium concentrations (4 mM), with only slight increases in the Cooper-Helmstetter Index that remained close to normal levels (∼2). However, at the more optimal growth temperature of 25°C (defined by growth rates^26^), LSUCC0530 exhibited a strikingly similar phenotype to HTCC1062 - higher ammonium concentrations induced both severe growth inhibition and pronounced aneuploidy (Cooper-Helmstetter Index > 6). These results demonstrated that nitrogen-induced cell cycle dysregulation and growth inhibition occurs in diverse SAR11 lineages from different habitats, regardless of the presence of additional nitrogen regulatory genes.

**Figure 6.**
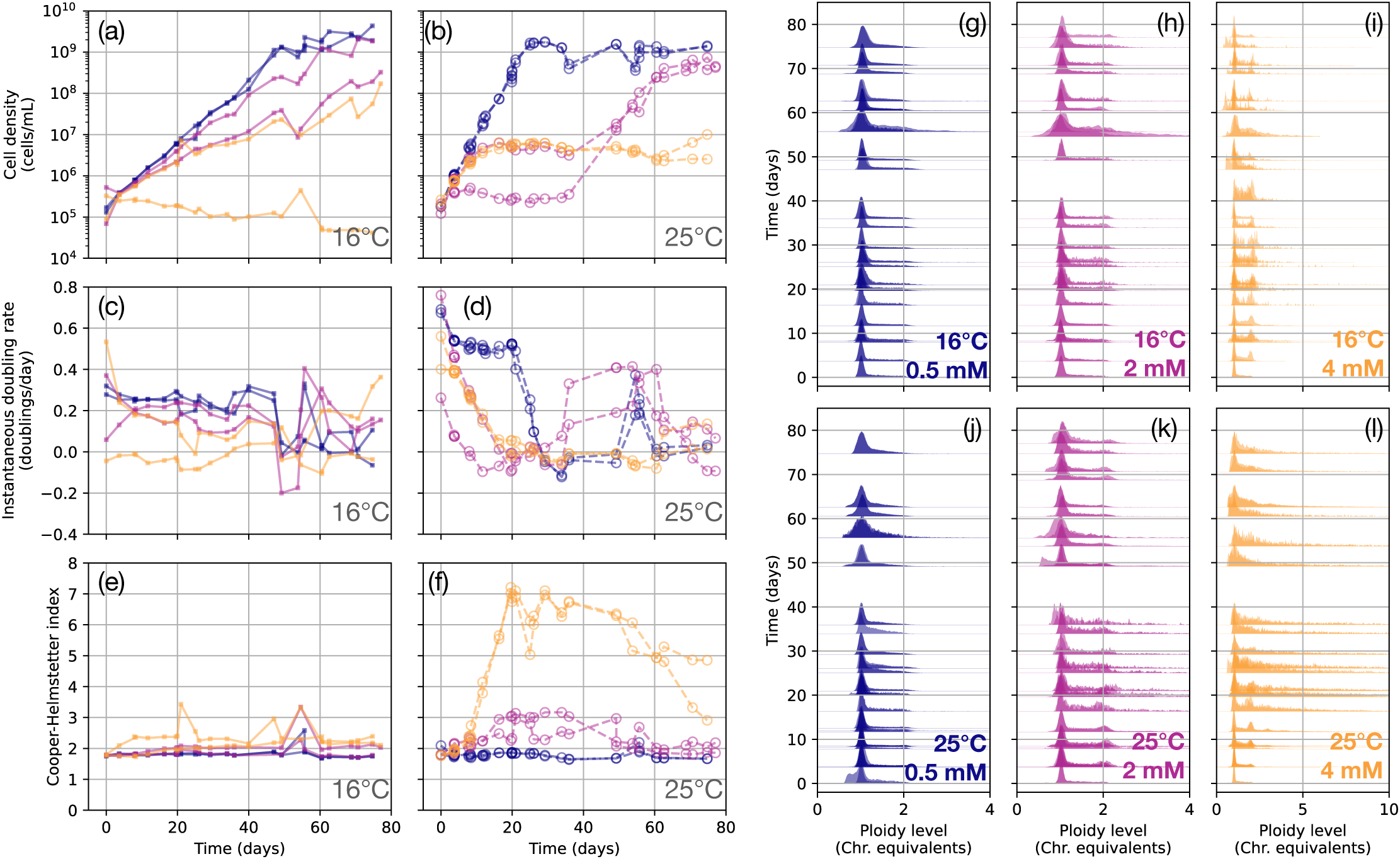
Cell cycle vulnerability extends to freshwater SAR11 strain LSUCC0530 with the P-II system. **(a-f)** Time course measurements of strain LSUCC0530 growth parameters under varying ammonium concentrations (0.5-4 mM, detailed nutrient concentrations are in **Supplementary Table Tab 16**) at two temperatures: (a,c,e) 16°C and (b,d,f) 25°C, showing (a-b) cell density, (c-d) instantaneous doubling rate, and (e-f) Cooper-Helmstetter Index. **(g-l)** Ploidy level distributions over time under different ammonium concentrations and temperatures: (g-i) 16°C and (j-l) 25°C, with ammonium concentrations of (g,j) 0.5 mM, (h,k) 2 mM, and (i,l) 4 mM NH_4_^+^. Each condition is in duplicates.

## Discussion

The physiological vulnerabilities we uncovered in SAR11 fundamentally deviate from established bacterial principles^54^. We propose that the widespread absence of multiple canonical cell cycle regulatory genes in SAR11 (**Fig. 1**) leads to the growth inhibition caused by various environmental perturbations detailed herein (**Fig. 4**). Similar negative growth effects of excessive nitrogen compared to carbon have also been identified in a SAR11 subclade 1a open ocean strain, HTCC7211^55,56^. Growth inhibition under excessive nitrogen contradicts a traditional understanding of nutrient-growth relationships that predict enhanced or unaffected growth^57,58^. Furthermore, our observations of increased mean cell sizes following growth inhibition with aneuploidy (**Extended Data Fig. 6**) show that SAR11 fails to follow the classical bacterial growth law where cell size positively correlates with growth rate^54,59^. The absence of cell division regulatory genes in SAR11 therefore appears to compromise fundamental cell size homeostasis mechanisms found in better-studied model bacteria^60^.

Our findings reveal restrictions on SAR11 growth rates beyond simple nutrient limitation. The optimal growth conditions we identified in this study were characterized by non-overlapping cell cycles, with maximum ploidy levels of 2 chromosome equivalents. The additional nutrients that would be expected to induce overlapping cell cycles and faster growth rates in *E. coli* and *B. subtilis* instead caused cell cycle disruption and growth inhibition in SAR11. This suggests that SAR11’s inability to achieve overlapping cell cycles, likely due to the absence of key regulatory genes like *minCD*, *seqA,* and *dam* (**Fig. 1**), constrains the organism to obligate slow growth. Notably, while mutants with single knockouts of these genes in model organisms (such as *E. coli*^29^) could still achieve overlapping cell cycles in rich media, these mutants retained most of their cell cycle regulatory machinery. Conversely, the lack of multiple cell cycle regulation genes in SAR11 appears to have imposed an upper limit on growth rate that is independent of nutrient availability, because it cannot safely accelerate growth through the overlapping cell cycle mechanisms that enable rapid division in well-regulated bacteria.

Although some of the concentrations used in our experiments were much higher than measured standing stocks in marine systems^61^, standing concentrations can be low due to high turnover rates^62^ and cells can still experience locally high concentrations of compounds, especially in the phycosphere^46,63^ (see a more detailed discussion in **Supplementary Text**). Regardless, our other results showing SAR11 cell cycle disruption from altered ratios of compounds, temperature shifts, and carbon source switches, where concentrations were maintained the same, indicate that the absolute concentration of compounds is not the only driver of dysregulation.

Furthermore, our observations align well with, and help explain, known ecological patterns of SAR11 in natural environments. Many regions of the open ocean are characterized by nitrogen limitation rather than carbon limitation^64^-conditions that would favor organisms like SAR11 that thrive with high C:N ratios. Indeed, SAR11 enrichment typically coincides with the growth phase of primary production but declines after phytoplankton blooms^19,20^, when C:N ratios and organic carbon compositions shift substantially^65^. The carbon landscape changes during bloom decay may parallel the shifts we tested in laboratory conditions. These patterns correlate with field observations of higher diploid frequencies (chromosome duplicated but undivided cells) and elevated cell death when SAR11 abundance declines during spring bloom^66^, suggesting potential connections between natural nutrient dynamics and the cell cycle disruptions we observed experimentally.

The loss of cell cycle regulatory genes likely reflects adaptive trade-offs driven by selection for genome streamlining in chronically oligotrophic environments. Under stable, low-nutrient conditions, maintenance of regulatory genes like *minCD* and *seqA* may provide diminishing returns, and their loss reduces the metabolic costs of expressing these genes. The SAR11 streamlined cell cycle machinery functions adequately under oligotrophic conditions (**Fig. 2e**), similarly to *B. subtilis minCD* mutants^30^, but lacks the regulatory buffering capacity to maintain homeostasis under environmental perturbations (**Fig. 3**). Given SAR11’s limited gene regulatory capacity^1,14,15^, cell cycle disruption may result from environmental perturbations shifting protein expression out of balance. Deviation in the stoichiometric ratios of proteins within the “fellowship of the Z-ring”^67^ causes cell cycle disruption in *E. coli*^51^, *C. crescentus*^31,68^, and *Mycobacterium smegmatis*^69^. Overall, the detailed mechanisms underlying SAR11 cell cycle disruption need further investigation with gene-expression studies.

Given that genome streamlining is common among dominant marine microorganisms adapted to nutrient-limited environments^1^, the cell cycle vulnerabilities we observed might be more universal and worth characterizing in further studies. In support of this hypothesis, we found that many other aquatic Alphaproteobacteria were missing some of these cell cycle regulatory genes based on our UniProt searches. Among all the widely distributed genes in the Alphaproteobacteria (*dam*, *chpT*, *ftsE*, *minC*, *minD*, *mipZ*, and *lemA*), we found that the SAR116 clade, *Candidatus* Puniceispirillales^70,71^, only possessed *minD*, and the *Parvularculales*^72^ were missing *minC*, with *dam* only present in 1/43 genomes with *dnaA* **(Supplementary Table Tabs 6, 10, 11)**. In contrast to the 1,098 *Rhodobacterales*^73^ that had *dnaA*, only 247 of them had *dam*, 10 had *ftsE*, 121 had *minC*, and 39 had *lemA* **(Supplementary Table Tabs 9, 10, 13)**. Therefore, the absence of these genes appears common among aquatic Alphaproteobacteria (SAR11 stands out as an extreme case), could be diagnostic of organisms adapted to oligotrophic environments, and might also set upper bounds on growth rates obtainable by these organisms.

The potential widespread cell cycle vulnerabilities among marine taxa might shift the way we model biogeochemical cycles. Conventional biogeochemical models typically assume that microbial growth and activity are primarily determined by bulk nutrient concentrations, following Monod kinetics or similar formulations^74,75^. Our findings challenge this paradigm by demonstrating that relative changes in nutrient conditions and compositional shifts may be equally or more important than absolute concentrations, particularly for streamlined organisms. If similar sensitivities are common among other streamlined taxa, then the ratios of nutrient standing stocks would influence not only the classical determination of nutrient limitation under Liebig’s Law of the Minimum^76,77^, but also taxon-specific physiological responses that shape ecosystem function.

Extensively missing cell cycle genes, especially in ancient lineages like SAR11 (estimated divergence from a common ancestor with other Alphaproteobacteria ∼700 million - 1 billion years ago^52,78^), raises an alternative evolutionary hypothesis that instead of these genes being lost through streamlining, their absence represents an ancestral state. In this scenario, the cell cycle regulation genes we see in *E. coli* and other fast growing organisms are adaptations for enhanced growth in nutrient-rich environments like animal guts. Indeed, it has been proposed that *minC*, *minD*, and *seqA* were missing from the last bacterial common ancestor (LBCA) core genome, but *lemA* and *sepF* were present^79^. That ancestral state reconstruction was not focused on Alphaproteobacteria and lacked SAR11 and other marine taxa. Expanded LBCA reconstructions in the future with a broader selection of genome-reduced organisms that include SAR11 could explore if SAR11 cell cycle gene absences are convergent with the LBCA through streamlining, reflective of the ancestral state, or a mixture of the two whereby the common ancestor of SAR11 and other Alphaproteobacteria lacked some of the cell cycle regulation genes and streamlining acted to remove those that remained.

The nutrient sensitivities we documented also have profound implications for culturing marine oligotrophs. Standard microbiological practices like nutrient enrichment and temperature optimization, or even simply shifting carbon sources, which is common when performing isolation experiments with artificial media or nutrient-amended natural media, may inadvertently trigger physiological disruptions that prevent successful cultivation. This may partially explain the “great plate count anomaly”^80^ and the extremely low isolation success rates for dominant organisms like SAR11 even with high throughput dilution-to-extinction approaches^81^. Many uncultured marine oligotrophs may share similar streamlining-associated vulnerabilities, requiring cultivation approaches that prioritize environmental stability and stoichiometric balance over absolute nutrient concentrations.

In conclusion, our study reveals how the absence of key regulatory genes in SAR11 has compromised its ability to coordinate cell cycle progression under environmental perturbations, indicating a likely evolutionary trade-off whereby the precise calibration of SAR11’s metabolism to oligotrophic conditions has enabled remarkable ecological dominance but at the cost of reduced physiological resilience. The direct coupling between environmental conditions and cellular physiology in these dominant marine microorganisms adds a critical dimension to ocean biogeochemistry, where nutrient dynamics and stoichiometric ratios may play more important roles than previously recognized in models based primarily on absolute concentration thresholds. If similar vulnerabilities are widespread among oligotrophic microorganisms, it will require new frameworks for understanding their ecology, evolution, cultivation, and responses to environmental change.

## Methods

### SAR11 pangenomics

We determined the presence or absence of cell cycle regulation marker genes across different SAR11 subclades with the pangenome generated previously for 470 unique genomes^27^. We took the aligned amino acid sequences of all the open reading frames from the previous pangenome summary, removed inserted gaps, and used InterProScan-5.75-106.0^82,83^ to generate updated annotations for 517,196 of 551,103 amino acid sequences (93.8%). The amino acid sequences and their InterProScan annotations are available on FigShare^84^. We targeted cell cycle proteins for analysis based on genetics studies of model organisms **(Fig. 1, Supplementary Table Tab 1)** where knock outs of the corresponding genes caused cell cycle disruption. We then searched for the gene and the organism on UniProt and identified the corresponding InterProScan ID. Generic InterProScan IDs, i.e., those that represent common protein features and therefore cover multiple genes, were excluded. We then looked for the corresponding InterProScan annotations of the target proteins in the SAR11 pangenome. The TipN protein from *C. crescentus*^40^ does not have any Interprocan IDs (as of September 5th, 2025), so we queried the amino acid sequence of TipN (Q2F5H9_CAUVC on UniProt) against all 551,103 SAR11 pangenome amino acid sequences using BLASTP v2.14.1 **(Supplementary Table Tab 15)**. The detailed presence/absence accounting of each target protein within each SAR11 genome is in **Supplementary Table Tab 3**.

### UniProt searches for targeted genes across all bacterial taxa

In addition to looking for cell cycle proteins in SAR11, we used UniProt (https://www.uniprot.org/)^85,86^ to examine the distribution of the same cell cycle proteins across other taxonomic groups at varying evolutionary distances from SAR11. All UniProt searches were conducted between July 22nd and September 5th, 2025. We took each of the InterProScan IDs in **Table 1**, e.g. IPR013317, and entered “(taxonomy_id:2) IPR013317” in the search box on UniProt. We grouped the search results by taxonomy: https://www.uniprot.org/uniprotkb?groupBy=taxonomy&query=%28taxonomy_id%3A2%29+IPR013317. The taxonomy dropdown list was expanded towards SAR11 HTCC1062, i.e., Bacteria, Pseudomonadati, Pseudomonadota, Alphaproteobacteria, Candidatus Pelagibacterales, Candidatus Pelagibacteraceae, Candidatus Pelagibacter, Pelagibacter ubique, Pelagibacter ubique (strain HTCC1062). We used the Anthropic Claude web interface (Sonnet 4) to parse the taxonomic hierarchy on the UniProt webpage and export this as csv tables. The dialogue is available through https://claude.ai/share/a932bef3-eac3-4c5b-8a12-3599b385d366. The exported csv tables were proofread manually **(Supplementary Table Tabs 4–14)**. As TipN from *C. crescentus* did not have an InterProScan ID, we obtained all the “Similar Proteins” from UniProt for the TipN designator “Q2F5H9_CAUVC” (**Supplementary Table Tab 16)**.

### Bacterial strains

The SAR11 representative *Candidatus* Pelagibacter ubique strain HTCC1062^87^ was generously provided by Dr. Stephen J. Giovannoni. Cultures of *Candidatus* Fonsibacter ubiquis strain LSUCC0530 were maintained in the Thrash Lab culture collection^26^. We prepared cryostocks in 10% DMSO or 10% glycerol and stored them in a −80°C freezer or liquid nitrogen. Differences in the reagents or storage conditions did not affect growth physiology in the subsequent cultivation experiments, e.g., **Fig.2a**, **Extended Data Fig. 2ef**, **Extended Data Fig.3**, where some replicates were revived from different cryoprotectant sources and showed reproducible phenotypes. *E. coli* strain NCM3722 was generously provided by Dr. Terence T. Hwa and was used for testing the applicability of SYBR ® Green 1 staining to infer ploidy levels (**Extended Data Fig. 9**). The cryostocks were preserved in 10% glycerol stocks and stored in −80°C freezer.

### Growth media

The media used in this study for growing SAR11 were adapted from three artificial seawater recipes: AMS1^18^, JW2^88^, and JW5^26^, with detailed chemical compositions provided in **Supplementary Table Tab 17 and 18**. For culturing strain HTCC1062, we based our formulation on AMS1 but made one key modification: we reduced the salinity from 36.1 ppt (AMS1) to 23.2 ppt, consistent with the JW2 recipe. This modification significantly improved the growth rate of HTCC1062 (**Extended Data Fig. 3a-b**), and we designated this revised medium as CCM2. For growing the freshwater SAR11 strain LSUCC0530, we further reduced the salinity of CCM2 to 1.45 ppt, matching the JW5 recipe, and referred to this modified medium as CCM5.

All media for growing SAR11 were prepared gravimetrically by assuming a density of 1.01 g/mL. This estimate was based on weighing 50 mL of prepared media in a Falcon tube with volumetric markings, yielding a mass of 50.3 g (1.006 g/mL), and was rounded to 1.01 g/mL for convenience. Accordingly, MilliQ water volumes were determined by weight rather than volume, estimated via e.g. a graduated cylinder, to improve accuracy (see **Supplementary Table Tab 18**).

To prepare 1 L of CCM2, we weighed 985.3703 g of MilliQ water into a 3.7% HCl acid-washed container (Nalgene™ Polycarbonate Bottles). We then added the following components: 15.8965 g NaCl, 7.0702 g MgCl_₂_ ·2H_₂_ O, 0.8401 g NaHCO_₃,_ 1.0098 g CaCl_₂_ ·2H_₂_ O, 0.4975 g KCl, 0.11 g sodium pyruvate (1 mM), 15 mg glycine (0.2 mM), and 2.98 mg L-methionine (0.02 mM). To this base, we added 2 mL of a 10,000x vitamin mix (the final concentration is 20x), 1 mL of a 1,000x iron mix, 10 μL of a 100,000x trace metal mix, and 0.2 mL of a 500 mM phosphate buffer. After all components were dissolved, the media were sterilized by vacuum filtration through a 0.2 μm filter (Nalgene™ Rapid-Flow™ PES filter material, 0.2μm, 75mm membrane) into Pyrex® glass bottles.

The concentrated stock solutions used for these additives were prepared and stored as follows. The 500 mM phosphate buffer (1 L, pH 8.0), based on the Cold Spring Harbor sodium phosphate buffer recipe ^89^, was prepared by dissolving 0.466 g Na_₂_ HPO_₄_ and 0.034 g NaH_₂_ PO_₄_ in 1,000 g of MilliQ water, followed by vacuum filtration through a 0.2 μm filter (Nalgene™ Rapid-Flow™ PES filter material, 0.2μm, 75mm membrane) and storage at room temperature. The 100,000x trace metal mix (1 L) was prepared by dissolving 0.18 g MnCl_₂_ ·4H_₂_ O, 20 mg ZnSO_₄_ ·H_₂_ O, 6.49 mg CoCl_₂,_ 6.57 mg Na_₂_ MoO_₄,_ 17.3 mg Na_₂_ SeO_₃,_ and 13 mg NiCl_₂_ in 1,000 g of MilliQ water. The solution was vacuum filter-sterilized as above, aliquoted, and stored at 4°C. The 1000x iron mix (1 L) consisted of 28 mg FeSO_₄_ ·7H_₂_ O and 81 mg nitrilotriacetic acid disodium salt (C_₆_ H_₇_ NO_₆_ Na_₂_), dissolved in 1,000 g of MilliQ water, sterilized by vacuum filtration as above, aliquoted, and stored at 4°C. To prepare the 10,000x vitamin mix (1 L), we dissolved 1 g NaHCO_₃,_ 1.69 g thiamin (B1), 2.6 mg riboflavin (B2), 0.985 g nicotinic acid (B3), 1.013 g calcium pantothenate (B5), 1.028 g pyridoxine (B6), 9.8 mg biotin (B7), 17.7 mg folic acid (B9), 9.5 mg vitamin B12, 0.901 g myo-inositol, and 82.3 mg 4-aminobenzoic acid in 1000 g of MilliQ water. The mixture was thoroughly dissolved, vacuum filtered through a 0.2 μm membrane as above, aliquoted into sterile polypropylene tubes (VWR^TM^), wrapped in foil to prevent light exposure, and stored at 4°C.

We used 30 mM aspartate with M63 media under 37°C for growing *E. coli*, which is a slow-growing condition that would keep *E. coli* in a non-overlapping cell cycle^54^, to help validate the applicability of using SYBR ® Green 1 for cellular DNA content quantification **(Extended Data Fig. 9)**. M63 minimal medium was prepared as follows: For 1 L of M63 media, we combined 3.0 g KH_₂_ PO_₄_, 7.0 g K_₂_ HPO_₄_ ·3H_₂_ O, 2.0 g (NH_4_)_₂_ SO_₄,_ 0.5 mg FeSO_₄_ ·7H_₂_ O, 0.246 g MgSO_₄_ ·7H_₂_ O (final concentration 1 mM), and 0.0111 g CaCl_₂_ (final concentration 0.1 mM) in approximately 900 mL of MilliQ water. After all components were dissolved, the solution was sterilized by vacuum filtration through a 0.2 μm filter (Nalgene™ Rapid-Flow™ PES filter material, 0.2μm, 75mm membrane). We then added the following filter-sterilized components: 10 mL of 1 M thiamine-HCl (final concentration 10 mg/L) and 30 mL of 1 M sodium aspartate (final concentration 30 mM). The final volume was adjusted to 1 L with sterile MilliQ water. The 1 M thiamine-HCl stock solution was prepared by dissolving 33.73 g thiamine hydrochloride in 100 mL of MilliQ water, vacuum filter-sterilized through a 0.2 μm filter as above, and stored at 4°C in the dark. The 1 M sodium aspartate stock solution was prepared by dissolving 17.71 g sodium L-aspartate monohydrate in 100 mL of MilliQ water, vacuum filter-sterilized through a 0.2 μm filter as above, and stored at 4°C.

### Bacterial Culturing

We used 3.7% HCl acid-washed and autoclaved polycarbonate flasks or bottles (Nalgene™, TriForest) for all SAR11 culture experiments. Cultures were grown at room temperature (17–23°C, **Extended Data Fig. 5**) in a regular storage cabinet without shaking, air sparging, or extra light control. We found that culturing in a volume greater than 400 mL provided better reproducibility, in contrast to the previously used 50 mL format^18^ **(Extended Data Figs. 3b-c, 8)**. Cells were revived from cyrostocks by either fully thawing the stocks (1 mL) or using a pipette tip to scratch a tiny amount (∼1–10 μL) and transferring them into fresh media. The revivals were then transferred again to new fresh media before reaching stationary phase. We restricted our number of transfers to less than 3 (at most 50 generations) to avoid potential mutations that might alter physiology. For each condition perturbation experiment, cultures were first established under optimal steady-state growth before transfer to experimental nutrient conditions. We defined optimal steady-state growth as having the optimal doubling rate among tested conditions (1 doubling/day) and steady-state ploidy level distribution (Cooper-Helmstetter Index ∼2). It is important to note that the physiological data presented in this study represent outcomes of condition shift experiments, where cultures were transferred from optimal steady-state growth conditions to test conditions, rather than cultures adapted to, or initiated directly in, the experimental media.

Note that the x-axis values in **Fig. 3d** represent additional chemical concentrations/compounds added to the standard medium. Specifically, since 1 mM glycine was present in control media (**Supplementary Table Tab 17**), “0 mM” indicates baseline 1 mM glycine, while “2 mM” represents 2 mM added to the standard 1 mM glycine (totaling 3 mM glycine). For **Fig. 4**, as there was no ammonium in the positive control medium, the concentrations are the final concentrations. The ammonium concentrations in **Fig. 6** are the final concentrations.

For growing *E. coli*, we revived the cryostock by scratching a tiny amount of unthawed samples (∼1–10 μL) using a pipette tip into the fresh liquid media in polycarbonate flasks.

### Cell counts

We measured cell densities by flow cytometry using an Accuri C6 Plus (BD) with a blue laser (488 nm) and sensors for both green (533 +/- 30 nm) and yellow (585 +/- 40 nm) fluorescence. We used a 96-well polystyrene plate (Corning® Costar®) for setting up the staining reaction. The staining protocol was strain specific to achieve clearly resolved peak distribution via green fluorescence area (reflecting ploidy level) during steady state growth **(Extended Data Fig. 9)**. For HTCC1062, each well contained 120 μL of liquid count sample, consisting of 12 μL of 100x SYBR ® Green 1 (Biotium or Lonza, final concentration 10x), 12 μL of 100x Tris-EDTA (Sigma-Aldrich ®, 10x final concentration — 100 mM Tris-HCl, 10 mM EDTA), 12 μL of 5% Glutaraldehyde (final concentration 0.5%), and 84 μL of bacterial culture. For LSUCC0530, each well contained 6 μL of 100x SYBR ® Green 1 (final concentration 5x) and 114 μL of bacterial culture. For *E. coli* NCM3722 (as a reference), each well consisted of 12 μL of 100x SYBR ® Green 1, 12 μL of 100x Tris-EDTA, 12 μL of 5% Glutaraldehyde, and 12 μL of RNAse (Invitrogen™ Ambion™ RNase Cocktail™. RNAse cocktail had little effect on the SAR11 signals). The 100x SYBR ® Green 1 was diluted from the 10,000x vendor stock (Biotium or Lonza) with 100x Tris-EDTA. The 5% Glutaraldehyde was diluted from the 25% vendor stock (Sigma-Aldrich ® Grade I) with MilliQ water. After staining, the plate was incubated in dark for at least 30 minutes before flow cytometric measurements. We maintained cell density within our count samples at below 1,000 cells/μL, and therefore the “bacterial culture” mentioned above consisted of an aliquot from the growth culture and MilliQ water as the diluent when concentrations in the growth culture exceeded 1,000 cells/µL. We used the “Medium” preset flow rate (35 μL/min, 16 μm core). The threshold was set at a minimum of 1,000 for green fluorescence intensity (labeled as “FL1-H” or “FITC-H” on the BD Accuri software). Based on the excitation/emission spectrum of SYBR ® Green 1 binding DNA, DNA fluorescence was expected to have a 3.5–6:1 ratio of green vs. yellow fluorescence (labeled as “FL2-H” or “PE-H” in the BD Accuri software). Therefore, we gated with a diagonal ellipse on the green vs. yellow fluorescence plot accordingly (**Extended Data Fig. 7b-e, g-l, Extended Data Fig. 9a,c,e,g, Supplementary Fig. 7a-e**). The cell density in the well was calculated by the number of events that were gated as cells divided by the total flow-through volume (30 μL). The dilution factor was then taken into account to calculate the cell density from the culture samples. A small portion of cell counts and ploidy level distributions, specifically those taken on day 55 in **Fig. 6**, were collected using Guava EasyCyte 5HT, with the InCyte 3.1 software. Sample preparation was completed identically as when using the Accuri. We used the preset medium flow rate (0.59 μL/sec) and gains for all the channels were default (100). The triggering channel was set as “Green-B Fluorescence (GRN-B-HLin)” (the green fluorescence intensity) with the threshold of 60.

### Estimation of instantaneous doubling time *T*

The method for estimating instantaneous doubling time is adapted from previous studies in the same lab^90,91^. We applied a sliding window for every three time points across the full growth curve, and applied a linear regression for *log2*(cell density) vs. time **(Extended Data Fig. 10)**. We used the slope +/- one standard error as the candidate instantaneous doubling rates. The candidate rates were assigned to the beginning, middle, and end of the sliding window time frame. Finally, we interpolated the spatial averages of the candidate rates as the instantaneous doubling rates vs. time. The implementation of the algorithm is included in our GitHub repository^92^.

### Ploidy level distribution

SYBR ® Green 1 selectively stains cellular DNA, and because SAR11 and *E. coli* do not exhibit autofluorescence, the resulting fluorescence areas (fluorescence intensities integrated over time for each flow cytometer event) reflect ploidy levels. When plotting histograms of green fluorescence areas from gated flow cytometry events, we observed peaks that corresponded to integral chromosome copy numbers for each strain (**Extended Data Fig. 9**). Linear regression of the green fluorescence area vs. theoretical DNA content, calculated by the expected number of chromosome equivalents multiplied by the corresponding genome size of the strain, demonstrated a strong correlation between theoretical DNA content and measured fluorescence (R² = 0.990, P < 0.001, n = 8), validating the use of fluorescence intensity as a quantitative measure of chromosome equivalents (**Extended Data Fig. 9i**). Therefore, the distributions of fluorescence signals (defined as *f(x)* where *x* is the green fluorescence area, and *f(x)* is the curve that represent the relative frequency of *x*) were mapped to ploidy level distributions (defined as *p(a)* where *a* is the ploidy level) as follows: we identified the peak regions that corresponded to one chromosome equivalent as in **Fig. 3h-j**, **Extended Data Fig. 6a-c,g-i** and noted the mean green fluorescence area as *x_G1_.* Then, *f(x): = p(a⋅ x_G1_), p(a): = f(x/x_G1_)*. To handle many samples in high throughput, we wrote a Python script to automate the gating and ploidy level extraction process, maintaining the same principles as manual analysis on the BD Accuri software described above. The Python script used the fcsparser package to read and parse raw flow cytometry files (.fcs). The detailed code is included in the GitHub repository^92^.

### Cooper-Helmstetter Index

We defined the Cooper-Helmstetter Index as the ratio between maximum and minimum ploidy level of a population. To account for variation in signal vs. noise during measurements, we tested different X% cutoffs that excluded outlier data and took the ratio between X% and (100-X)% percentiles. We found that removing the highest and lowest 10% of the data resulted in an approximately 2:1 Cooper-Helmstetter Index for the *E. coli* control data under coordinated cell cycles from previous studies, reflecting the coordination between chromosome replication and cell division **(Supplementary Text)**. Therefore, unless otherwise specified, the Cooper-Helmstetter Index values reported in this study used a 10% cutoff.

### Estimation of B, C, and D periods based on ploidy level distributions

One can simulate the expected ploidy level distribution based on the ratio between B, C, and D period duration and the generation time *τ* — B/τ, C/_T_ and D/_T_^45^. This analytical simulation was based on the Cooper-Helmstetter model whereby parent cells have 2x the DNA content of newly divided children cells^43^. However, this method can only be applied to simulate steady-state populations. Therefore, we only used this method to calculate B, C, and D periods in samples where the Cooper-Helmstetter Index was smaller than 3. To estimate B/_T_, C/_T_, and D/_T_, previous studies tested simulations that best matched the ploidy level distribution, either via manual iteration^93^ or closed-source programming^94^. We used Python for our implementation, available on GitHub^92^. Notably, the ploidy level distribution only estimates B/_T_, C/_T_, and D/_T_, i.e. the ratio between the durations of each period and the doubling time, so we calculated absolute durations of B, C, and D periods by multiplying B/_T_, C/_T_, and D/_T_ by T. To verify our estimations of B, C, and D periods, we did DNA “runoff” experiments through antibiotics treatments. The details of our cell cycle dynamics estimation and verification could be found in **Supplementary Text and Figs. S7,S8**.

### Cell lysis experiments

To investigate the fate of polyploid cells and validate our observations of increased background fluorescence in high ammonium samples **(Extended Data Fig. 7)**, we conducted controlled cell lysis experiments. We added 50 μL of DNA/RNA lysis buffer (Zymo Research) to 1 mL aliquots of culture samples in 2 mL microcentrifuge tubes. Flow cytometry measurements were taken at regular intervals (0.0, 1.2, 2.4, 3.6, 6.0, and 10.1 hours) to track changes in cellular integrity and fluorescence patterns. The gated fraction was calculated as the percentage of events falling within the elliptical gate defined by the green versus yellow fluorescence ratio characteristic of intact, DNA-stained cells, as shown in **Extended Data Fig. 7b-e, g-l.**

### Subpopulation growth model

We implemented a quantitative subpopulation growth model as previously developed^81^: *λ_p_ = λ⋅ log2(1+Φ)*, where *λ_p_*is the population doubling rate, *λ* is the doubling rate of the active subpopulation (fraction *Φ* of the total), assuming cells in the remaining fraction *(1-Φ)* do not divide. This model assumes exponential growth is driven by a fraction of cells dividing normally. We assumed across all ammonium levels that active subpopulations maintained the same doubling rate (λ = 1.1 divisions/day) as observed under no ammonium conditions, where *Φ* = 1.0 (all cells actively dividing). Under excessive nitrogen conditions, *Φ* < 1 was determined by comparing ploidy level distributions to identify cells with ≥ 2 chromosome equivalents resulting from division inhibition. We defined the ploidy level distribution of positive control (*Φ* = 1.0) as *p_0_(a)*, *a* represents the ploidy level in number of chromosome equivalents, the ploidy level distribution of samples with aneuploidy (*Φ* < 1.0) as *p(a)*. To estimate *Φ*, we normalized both ploidy distributions (p₀(a) and p(a)) to the same height at the one-chromosome peak (*a = 1*), ensuring comparability **(Fig. 5ab)**. Then, *Φ* was calculated as the ratio of their total areas: *Φ = ∫_a_ [p_0_(a)/p_0_(1)] ÷ ∫_a_ [p(a)/p(1)]*. This represents the proportion of cells under the experimental condition that match the DNA content distribution of normally dividing cells.

### Dynamic light scattering

Dynamic light scattering (DLS) measurements were performed using a Wyatt Mobius Zetasizer at the Center of Excellence in NanoBiophysics at USC. 2 mL of live cell culture was aliquoted into the cuvette (BRAND ® PS, 1.5 mL semi-micro) and loaded into the laser slot of the DLS machine. Data acquisition and instrument control were managed through the DYNAMICS V7 software. For all measurements, we applied the following parameters: DLS acquisition time of 5 seconds, read interval of 1 second, 20 DLS acquisitions per measurement, auto-attenuation time limit of 60 seconds, and normal laser mode. We utilized the pre-configured “Sodium Chloride 2.5%” solvent profile with the following specifications: refractive index of 1.3374 (measured at 589 nm and 20°C), aqueous temperature model, viscosity of 1.044 cP at 20°C, fixed dielectric constant model, and a dielectric constant value of 74.39161493. To export the DLS measurements, we selected the “regularization graph” function in the software and set the resolution bar to maximum (highest resolution) in the control panel. We configured the regularization graph with “Radius” as the x-axis and “%Number” as the y-axis, using “Rayleigh Spheres” as the analytical model. This configuration displayed a frequency histogram of Rayleigh sphere radii. Data were exported by right-clicking the cursor and selecting “Export” to obtain the histogram data. The mean Rayleigh sphere radius was calculated by taking the weighted average (dot product) of the frequency values (%Number) and their corresponding radii from the histogram.

### Epifluorescence microscopy

We stained the HTCC1062 cells in 15 mL Falcon tubes, in the same proportion as our flow cytometry staining protocol, with 7mL of cell culture, 1mL of 100x SYBR ® Green 1, 1mL of 100x Tris-EDTA, and 1mL of 5% Glutaraldehyde. The Falcon tube was incubated in the dark for at least 30 minutes, then harvested with 25 mm black polycarbonate filters (Isopore^TM^ 0.2 μm Blk PC membrane) through vacuum filtration. Filters were then placed on glass slides and imaged through fluorescence microscope Axioskop 2 with AxioCam Cm1 (Zeiss).

### Statistics & Reproducibility

No statistical method was used to predetermine sample size. Sample sizes varied across experiments based on experimental design and practical constraints, ranging from 2-10 biological replicates per condition. Discrepancies in replicate numbers resulted from multiple considerations: practical constraints on the number of conditions that could be simultaneously maintained, the use of gradient experiments (e.g., increasing ammonium concentrations) where reproducibility was confirmed by consistent trends across the gradient of conditions rather than solely by replicate numbers at individual conditions, and prioritization of resources toward optimal growth conditions that received repeated experimental attention across multiple independent batches. No data were excluded from the analyses. The experiments were not randomized, with cultures assigned to experimental conditions based on the experimental design. The investigators were not blinded to allocation during experiments and outcome assessment. Growth curve analyses used least-squares linear regression with a sliding window approach (across three time points) to estimate instantaneous doubling rates. Two-sample t-tests were performed to compare doubling rates and saturation cell densities between conditions. For ploidy level distribution analyses, linear regression validated the relationship between green fluorescence area and theoretical DNA content (R² = 0.990, P < 0.001, n = 8). The Cooper-Helmstetter Index was calculated using a 10% cutoff (ratio between 90th and 10th percentiles), which was validated to produce approximately 2:1 ratios for *E. coli* control data under coordinated cell cycles. Reproducibility was rigorously ensured through repeated independent experimental batches conducted over the course of the study. Through repeated experimentation across multiple batches, we consistently identified optimal growth conditions characterized by approximately one day doubling time and Cooper-Helmstetter Index values around 2, demonstrating the robustness and reliability of our experimental approach. Custom Python scripts for all the analyses pipelines, including growth curve analysis and automated flow cytometry data analysis, as well as those for main figure generation, are available on GitHub^92^.

## Data Availability

Epifluorescence microscopic images beyond those of **Extended Data Fig. 6**^95^ and the results of the SAR11 InterProScan analyses and prior CheckM output for all taxa^27^ are hosted externally at FigShare^84^.

Access to the source data (including growth curves and flow cytometry/ploidy level distributions data) to all the main figures (**Fig. 1-6**) are available at Supplementary Table Tabs 1-3 and our GitHub repository^92^. The Google Colab notebooks in the GitHub repository provide “push-button” pipeline to read and index the raw flow cytometry files and therefore readers are able to follow how the files got turned into the ploidy level plots in the paper.

## Code Availability

We provide python code in Google Colab Notebook format in our GitHub repository^92^. The code include the essential data analyses pipeline to recreate the figures.

## Acknowledgments

We acknowledge the Center for Advanced Research Computing at the University of Southern California for providing computing resources that have contributed to the research results reported within this publication (https://carc.usc.edu), as well as the Center of Excellence in NanoBiophysics at USC. We are grateful to Stephen J. Giovannoni from Oregon State University providing the SAR11 strain HTCC1062, and Terence T. Hwa from University of California, San Diego for providing the *E. coli* strain NCM3722. We appreciate invaluable conversations with, and feedback from, Terence T. Hwa, Chenli Liu, Suckjoon Jun, Ying-Chih Chuang, Naomi M. Levine, Noelle A. Held, and Steven E. Finkel. Funding for this work was provided by a Simons Foundation Early Career Investigator in Marine Microbial Ecology and Evolution Award and a Simons Foundation Investigator in Aquatic Microbial Ecology Award to JCT.

## Author Contributions

C.C. contributed to study development, designed and conducted growth experiments, developed flow cytometry methodology, created the code for flow cytometry data analysis, growth rate calculations, and data visualization, developed the subpopulation model, performed all bioinformatics analyses, performed fluorescence microscopy, dynamic light scattering measurements and wrote the paper. B.D.B. conducted control growth experiments. P.S. helped with fluorescence microscopy. H.A. and C.B. helped with media preparation and growth measurements. C.B. helped with media preparation and dynamic light scattering measurements. K.A.E. helped with formalization of the subpopulation model. R.T. assisted with flow cytometry method development. J.C.T. developed the study, helped with experimental design and bioinformatics strategy, obtained funding for the work, assisted with data analysis and visualization, and wrote the paper. All authors contributed to proofing and editing the final manuscript.

## Supplementary information

**Supplementary Information|** The supplementary materials include: **(1)** supplementary text covering in-depth evaluation of the cell cycle marker genes that we chose in **Fig. 1** and their presence or absence in SAR11 genomes (**Section 1**), a basic background discussion of bacterial cell cycles^28^, reanalysis of population ploidy levels from previous studies for *E. coli* ^29,45,98^, *V. cholerae*^99^ (**Section 2**), and SAR11 HTCC1062^18^ (**Section 3**), reanalysis of DNA fluorescence microscopic image of SAR11 HTCC1062^18^, the pipeline for flow cytometry data analysis (**Section 4**), our DNA “runoff” experiments with antibiotic treatments for cell cycle dynamics validation in HTCC1062 (**Section 5**), and additional points of discussion stemming from our results (**Section 6**). **(2)** 8 supplementary figures **(Fig. S1-S8)** and their captions. **(3)** Detailed descriptions of the 24 supplementary tables **(Supplementary Table Tabs 1-24)** that contain additional data and analyses.

## Extended Data Figures

**Extended Data Figure 1.**
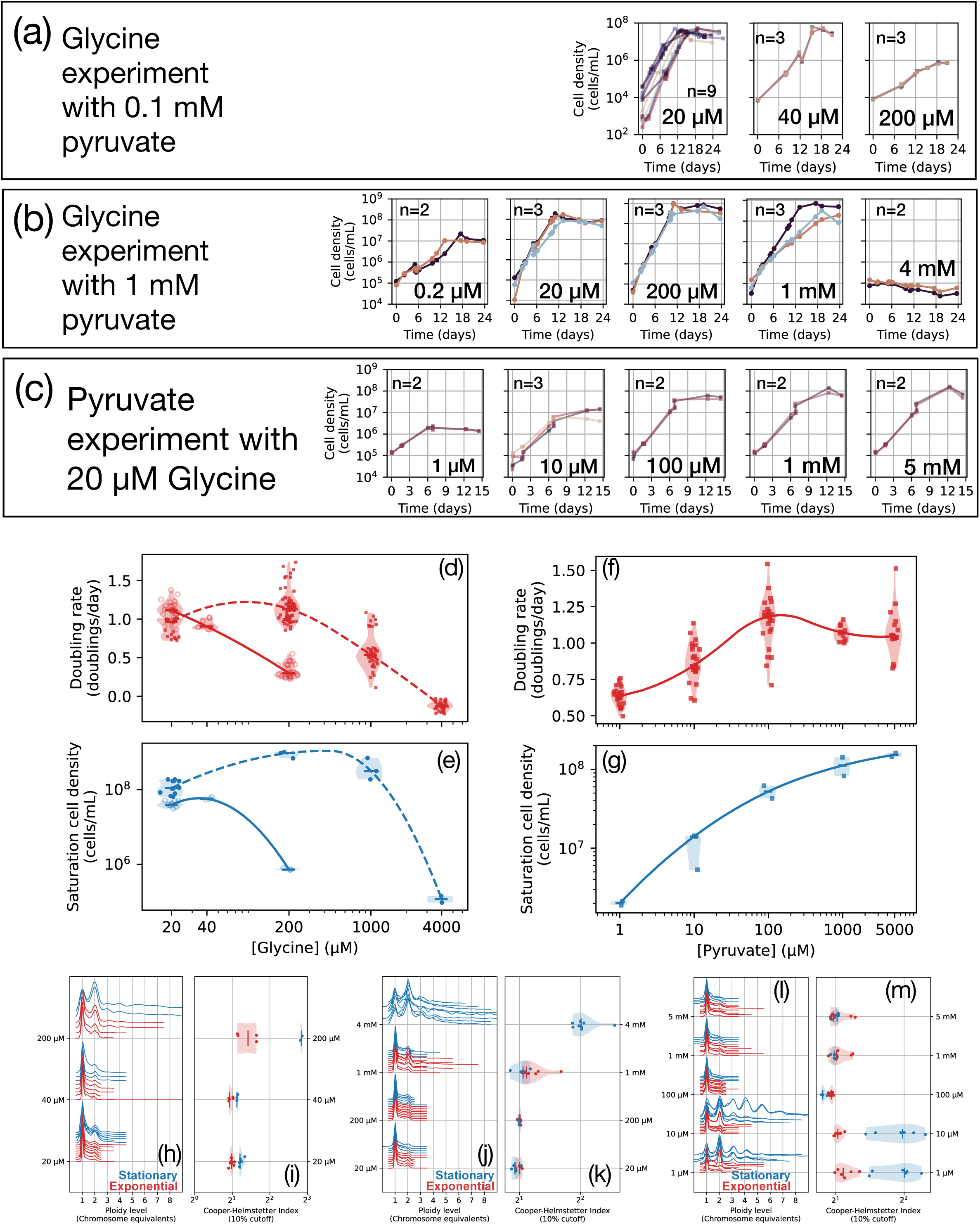
Effect of pyruvate:glycine ratios on SAR11 growth and cell cycle regulation. **(a-c)** Growth curves showing SAR11 responses to different glycine and pyruvate concentrations: (a) glycine experiment with 0.1 mM pyruvate, anchor condition 20 μM glycine; (b) glycine experiment with 1 mM pyruvate, anchor condition 200 μM glycine; (c) pyruvate experiment with 20 μM glycine, anchor condition 1 mM pyruvate. **(d-e)** Doubling rates and saturation cell density under different glycine concentrations. Dashed curves represent high pyruvate (1 mM) and vitamins (20X); solid curves represent low pyruvate (0.1 mM) and vitamins (2X). **(f-g)** Doubling rates and saturation cell density under various pyruvate concentrations with fixed glycine (20 μM) and vitamins (20X). **(h-k)** Ploidy level distributions across exponential (red) and stationary (blue) phases and Cooper-Helmstetter Index: (h,i) low pyruvate and vitamin conditions (solid curves in d,e); (j,k) high pyruvate and vitamin conditions (dashed curves in d,e). For 4 mM glycine in high pyruvate conditions (j,k), absence of red data indicates no growth. **(l,m)** Ploidy level distributions and Cooper-Helmstetter Index under various pyruvate concentrations corresponding to panels (f,g). Media compositions are provided in **Supplementary Table Tab 17**.

**Extended Data Figure 2.**
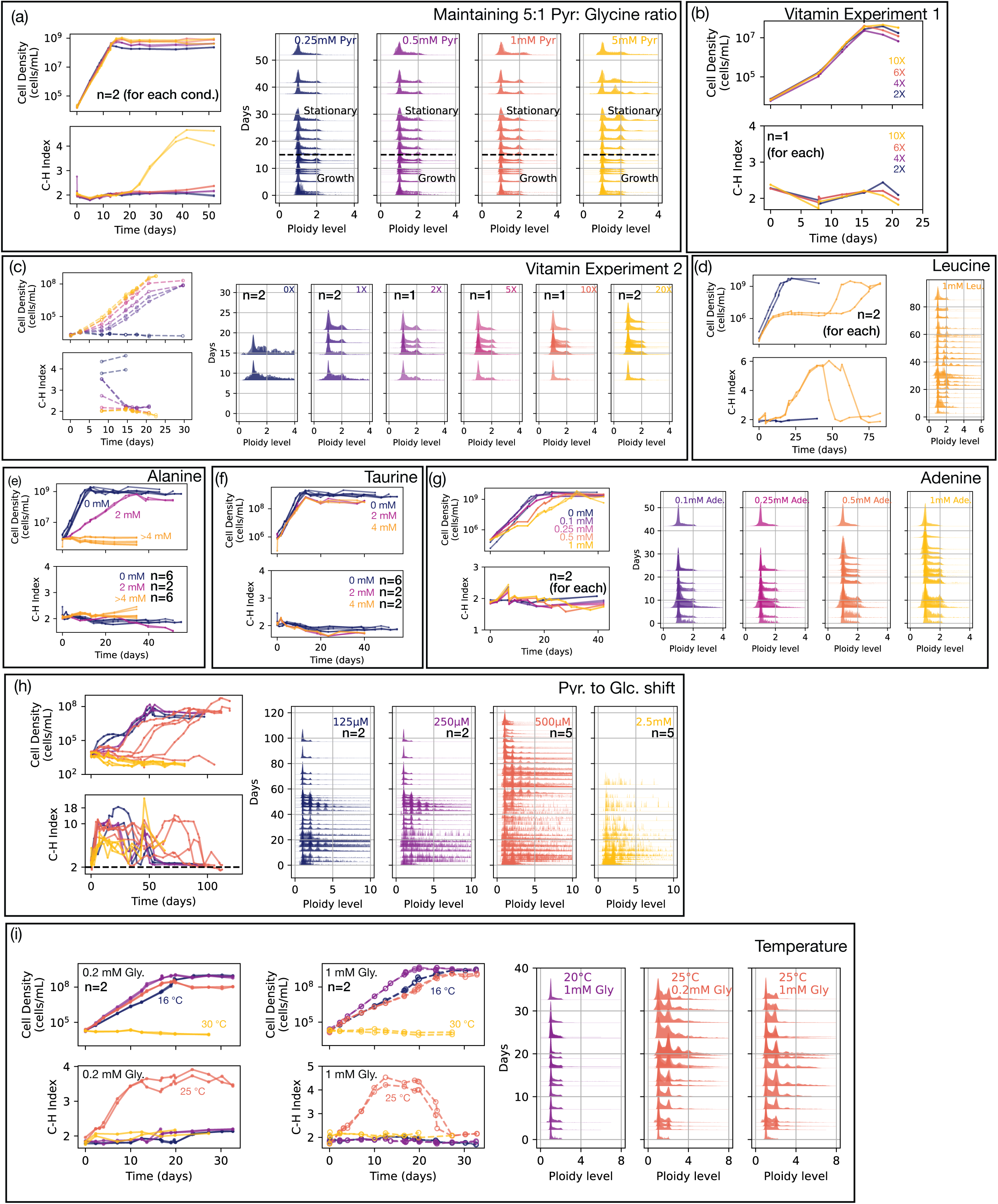
Growth curves and ploidy distributions for experiments described in Fig. 3. Detailed media compositions are provided in **Supplementary Table Tab 17.** For conditions where Cooper-Helmstetter Index remained around 2 throughout the time course, detailed ploidy distributions over time are not shown as they are essentially identical to the positive control in (a). **(a)** Raw data of Fig. 3b, showing growth curves, Cooper-Helmstetter Index over time, and ploidy distributions during growth and stationary phases. **(b,c)** Results of two independent vitamin experiments under different pyruvate (100 μM vs. 1 mM), glycine (20 μM vs. 200 μM) and ammonium chloride (800 μM vs. 0) concentrations. **(d)** Excessive leucine experiment. **(e)** Corresponds to the alanine experiment data in Fig. 3f. **(f)** Physiological effect of excessive taurine. **(g)** Effect of excessive adenine. **(h)** Pyruvate to glucose shift experiment in Fig. 3e. **(i)** Temperature experiment in Fig. 3d.

**Extended Data Figure 3.**
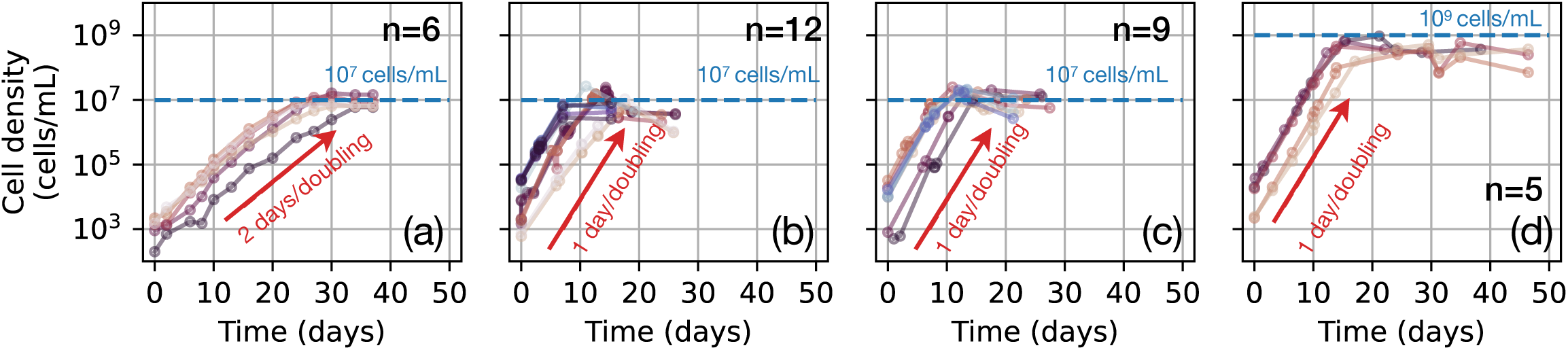
Growth curves of HTCC1062 under. **(a)** adapted AMS1 media in 50 mL, the specific concentration of each vitamin was slightly different from the original AMS1 recipe^18^. **(b)** “1x” condition as in Fig. 2 in 50 mL, **(c)** “1x” condition in large volume (> 500 mL), and **(d)** CCM2 media in large volume. The detailed media recipes can be found in **Supplementary Table Tabs 17 and 18**. The “1x” condition in (b) and (c) have the same nutrient concentrations as AMS1 media, but under lower salinity (23.3 ppt vs. 36.1 ppt). Discrepancies in the number of replicates resulted from performing different sets of experiments with different volumes. We have included data from all experiments conducted. The growth curve data are in **Supplementary Table Tab 21**.

**Extended Data Figure 4.**
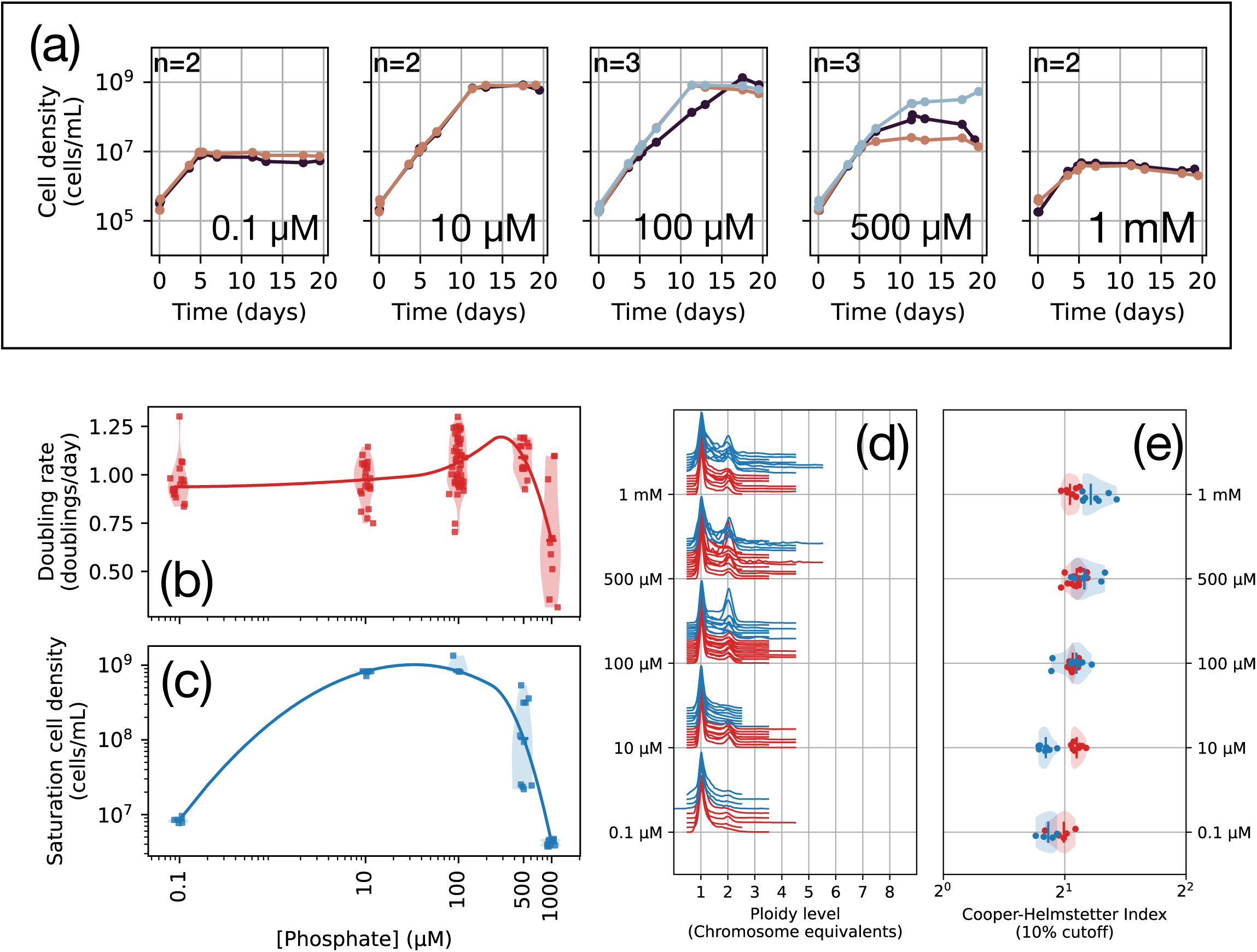
HTCC1062 cell cycle physiology under various phosphate concentrations, including (a) growth curves, (b) doubling rates, (c) saturation cell density, (d) ploidy level distribution, (e) Cooper-Helmstetter Index (10% cutoff). 100 μM was the anchor condition that was transferred to each experimental condition. For the 0.1 μM sample, we made 1L of phosphate-free media to which we transferred 1 mL of the seed culture (at 100 μM phosphate starting concentration). The 1000-fold dilution therefore set the upper limit of the carried-over phosphate at 0.1 μM. The detailed media recipes are in **Supplementary Table Tab 18**.

**Extended Data Figure 5.**
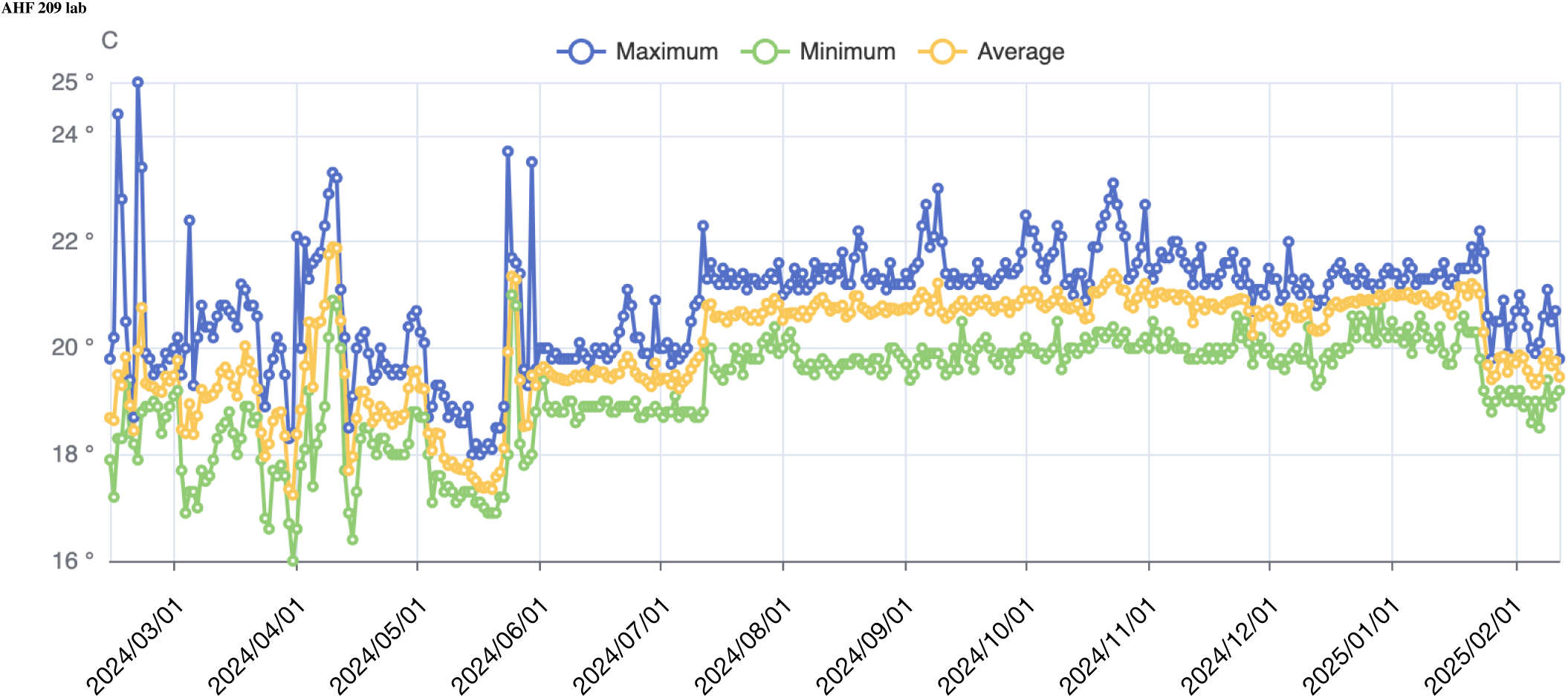
Laboratory room temperature monitoring during experiments. Temperature measurements (°C) over time showing maximum (blue), minimum (green), and average (orange) daily temperatures in the laboratory where SAR11 HTCC1062 cultures were maintained. Data collected using iMonnit temperature monitoring system. Room temperature typically fluctuated between 17-23°C, which corresponds to the “room temperature” conditions referenced throughout the experiments.

**Extended Data Figure 6.**
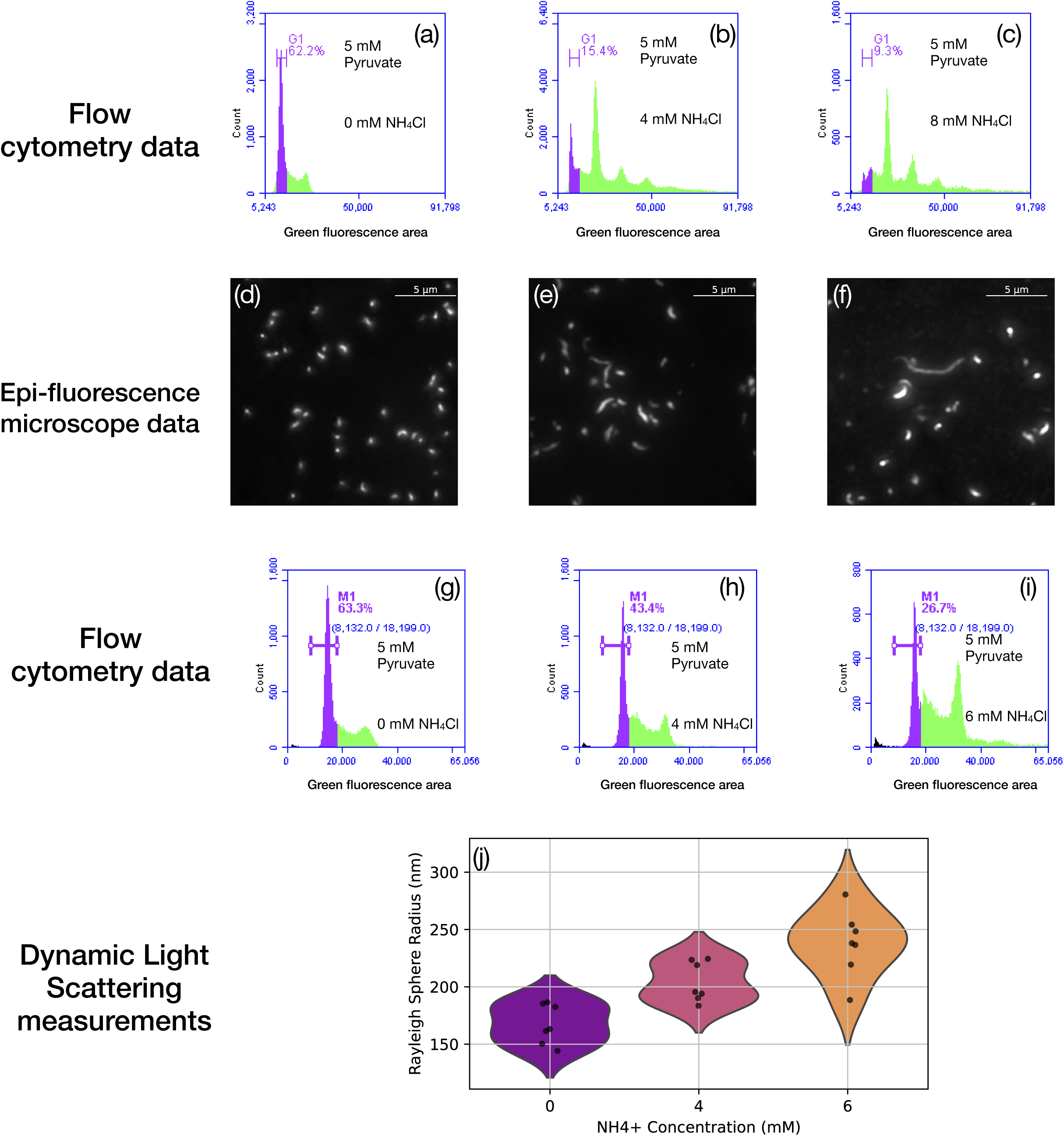
Elevated ammonium induces cell size polymorphism and morphological abnormalities in SAR11. **(a-f)** First experimental batch showing: Flow cytometry data of histograms of green fluorescence areas from exponential phase cultures with (a) 0 mM, (b) 4 mM, and (c) 8 mM NH_4_ Cl, revealing progressive increase in polyploidy with higher ammonium levels. In these green fluorescence area histogram, the population that we identified as cells with one copy number of chromosomes are gated and labeled as ‘G1’ and highlighted in purple. (d-f) Corresponding epifluorescence microscopy images of SYBR ® Green-stained cells from the same cultures, revealing normal rod-shaped morphology in control conditions versus elongated filamentous forms in high ammonium conditions. Scale bars = 5 μm. Microscopic images were taken at least 6 times at different locations of the slide for each sample: n=6 for 0 mM, n=16 for 4 mM, and n=43 for 8 mM, which are available on FigShare^95^. More images were taken for samples with higher NH_4_ Cl concentrations due to confirming the appearance of fillamentous cells were being found everywhere. **(g-j)** Independent from the data shown in (a-f), second experimental batch showing: Flow cytometry histograms of green fluorescence areas during exponential growth phase with different ammonium concentrations: (g) 0 mM, (h) 4 mM, and (i) 6 mM NH_4_ Cl. We gated and labeled the one-chromosome population as ‘M1’. The ranges of our manual gate are also displayed, as minimum green fluorescence area = 8132.0 (arbitrary unit), maximum green fluorescence area = 18199.0 (arbitrary unit). (j) Dynamic light scattering measurements of cell size (Rayleigh sphere radius) from the same cultures, demonstrating significantly increased cell size with higher ammonium concentrations. The dynamic light scattering source data is available in **Supplementary Table Tab 22**.

**Extended Data Figure 7.**
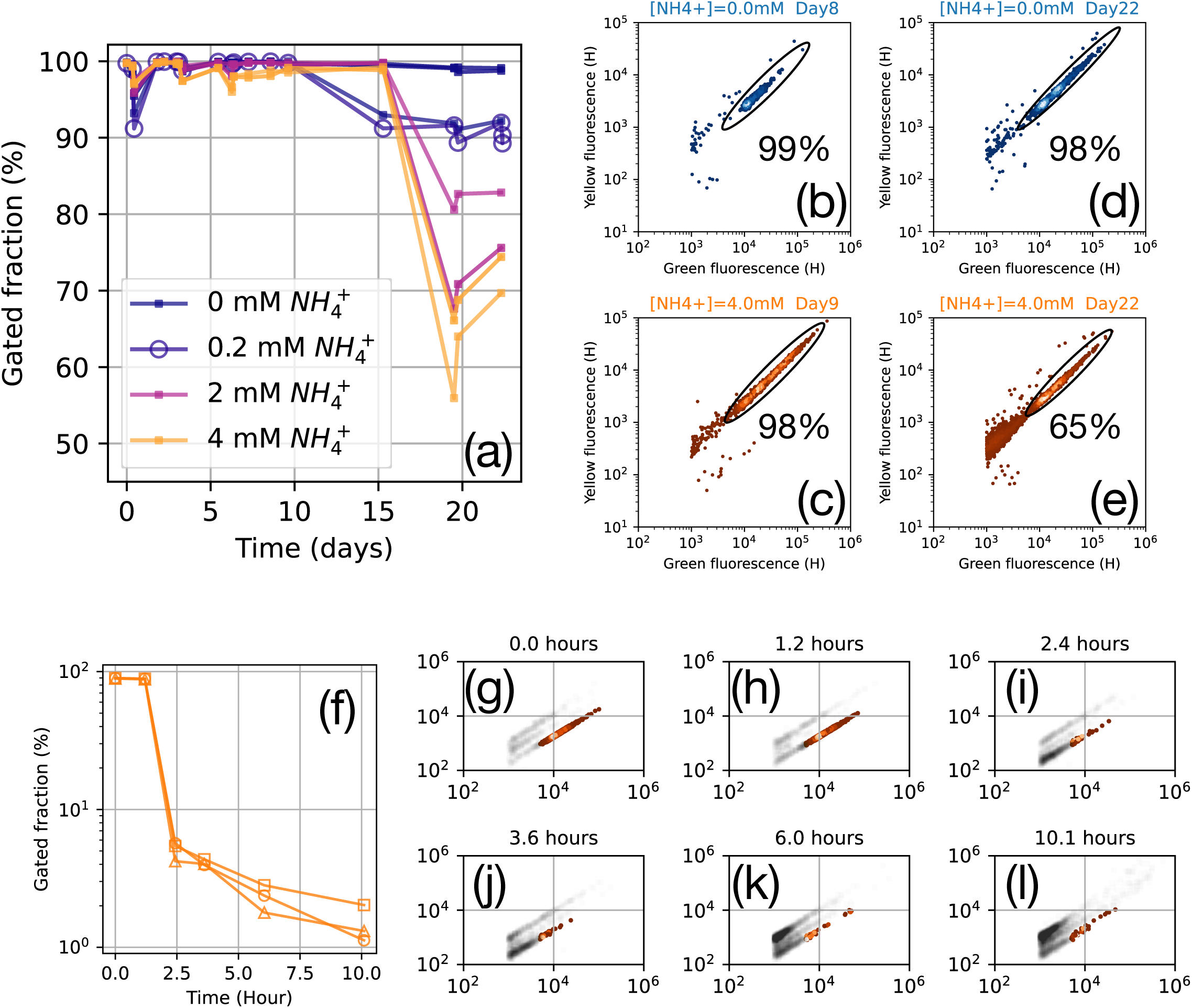
Flow cytometry gated fraction analysis and cell lysis experiments. **(a)** Variation of flow cytometry gated fraction at each time point across the growth curves under different ammonium chloride concentrations in Figure 3. Examples of the calculation of gated fractions are shown in (b-e). **(b-e)** Flow cytometry scatter plots showing green versus yellow fluorescence with gating ellipses and percentages of gated events at different time points and ammonium concentrations: (b) 0 mM NH_4+_ day 8, (c) 4 mM NH_4 +_ day 9, (d) 0 mM NH_4 +_ day 22, and (e) 4 mM NH_4 +_ day 22. (f) Time course of gated fraction decline during lysis buffer treatment. (g-l) Flow cytometry scatter plots showing the progressive breakdown of cellular integrity over time following addition of lysis buffer, with time points from 0.0 to 10.1 hours demonstrating the transition from intact cells to cellular debris.

**Extended Data Figure 8.**
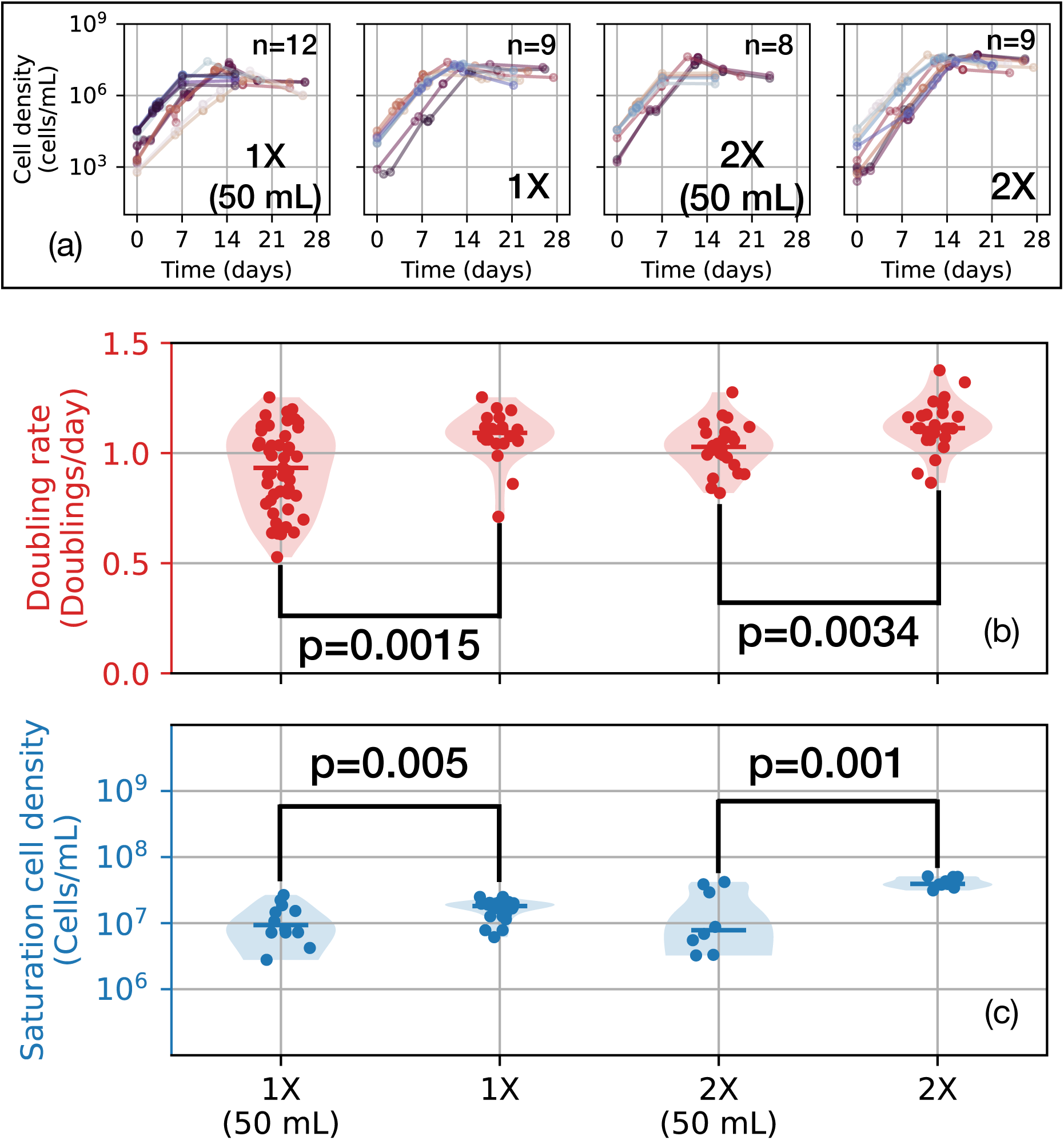
Effect of culturing volumes on the growth physiology. Samples labeled with “(50 mL)” are 50 mL liquid culturing in 125 mL flasks, otherwise are in the normal culturing conditions (> 400 mL as described in **‘Bacterial culturing’, Methods**). “1X” and “2X” media are the same as in Fig.2 and **Extended Data Fig. 1**. Here, we are showing the **(a)** growth curves, **(b, c)** violin plots with data points, and the p-value of the two-sample two-sided t-test for doubling rates and saturation cell densities. Discrepancies in the number of replicates resulted from performing different sets of experiments with different volumes. Growth curve source data are available in **Supplementary Table Tab 23**.

**Extended Data Figure 9.**
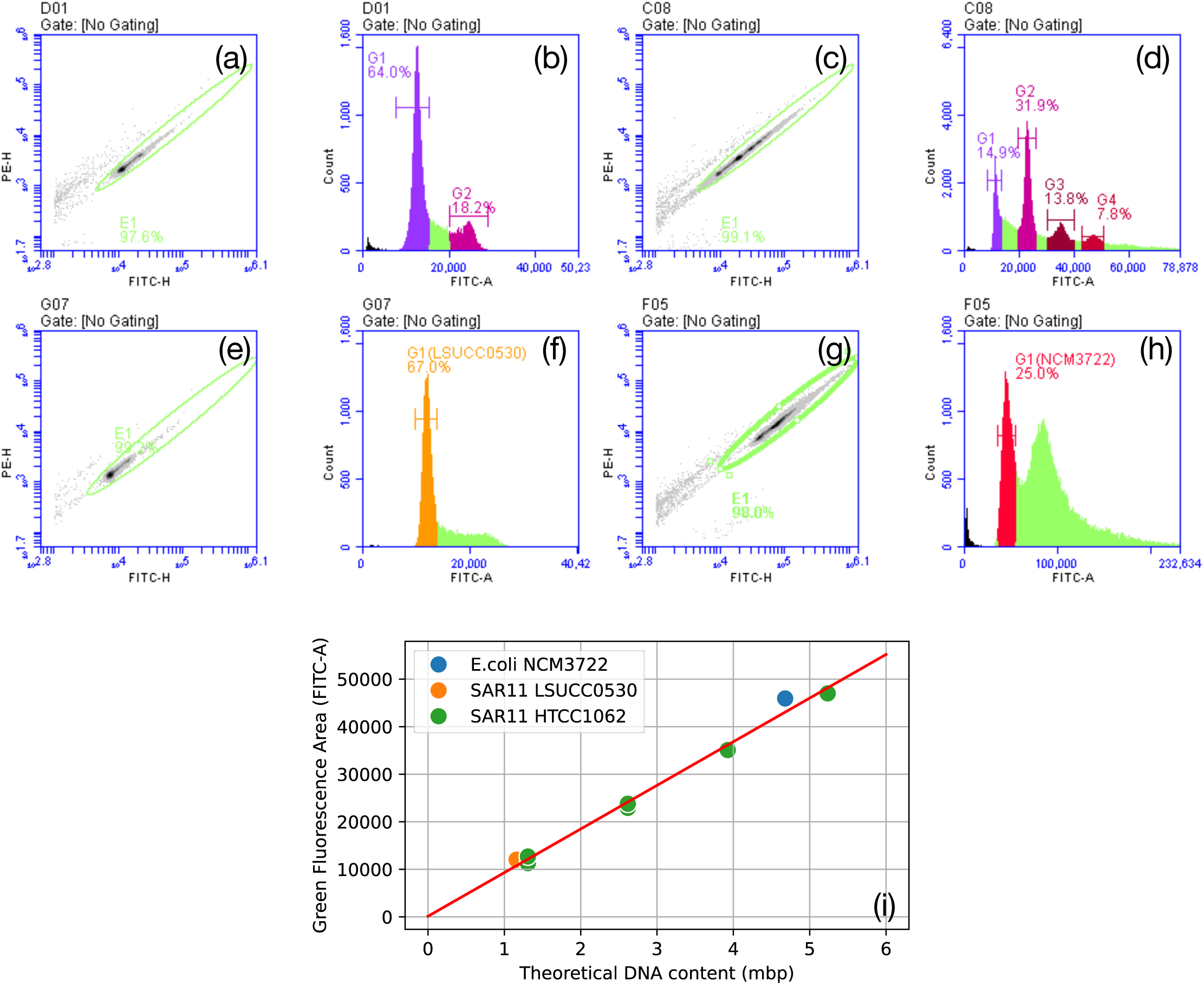
Flow cytometry gating strategy and ploidy calibration. **(a-h)** Flow cytometry scatter plots showing green versus yellow fluorescence (left panels) and corresponding DNA content histograms (right panels) for different bacterial strains and conditions: (a-b) SAR11 HTCC1062 samples with coordinated cell cycles, (c-d) SAR11 HTCC1062 samples with polyploidy, (e-f) SAR11 LSUCC0530, and (g-h) *E. coli* NCM3722. Green diagonal gates define cellular events based on the expected 3.5-6:1 ratio of green to yellow fluorescence for SYBR ® Green-stained DNA. In the histograms, peaks are gated and labeled according to genome equivalents (G1 = 1 chromosome, G2 = 2 chromosomes, etc.), with percentages indicating the proportion of events within each ploidy peak. **(i)** Linear regression between theoretical DNA content and green fluorescence area across different bacterial strains. The underlying data are in **Supplementary Table Tab. 24**.

**Extended Data Figure 10.**
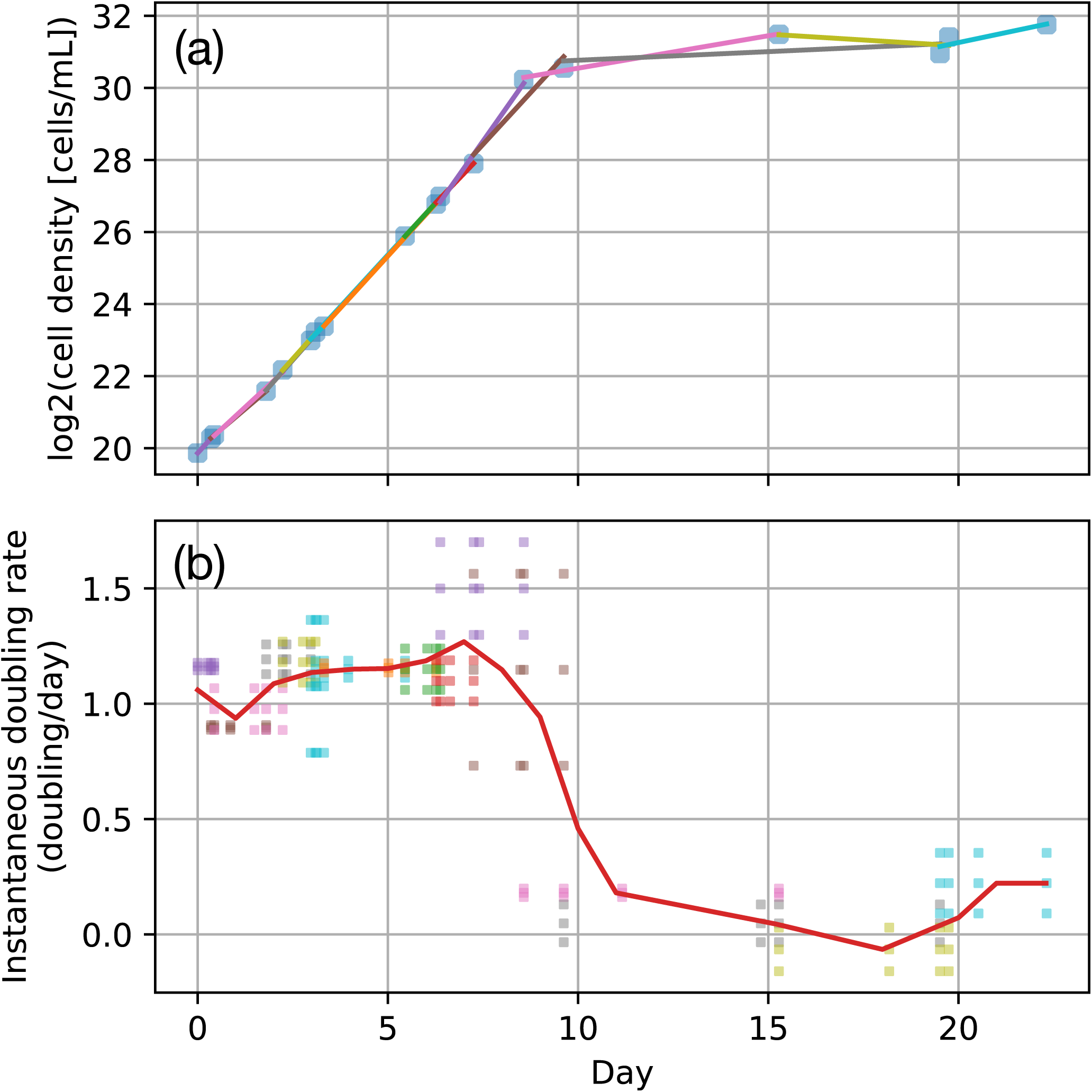
Example growth curve analysis. **(a)** Log 2 transformed cell density across time. Lines in different colors are the least squares linear regression fit within a sliding window of 3 time points across the full growth curve. **(b)** The estimation of instantaneous doubling rates for each sliding window are plotted as scattering dots with the corresponding color as in (a). The red curve is the non-parametric spatial average fit.

## References

1. Giovannoni, S. J., Thrash, J. C. & Temperton, B. Implications of streamlining theory for microbial ecology. ISME J. 8, 1553–1565 (2014).

2. McCutcheon, J. P. & Moran, N. A. Extreme genome reduction in symbiotic bacteria. Nat. Rev. Microbiol. 10, 13–26 (2011).

3. Morris, R. M. et al. SAR11 clade dominates ocean surface bacterioplankton communities. Nature 420, 806–810 (2002).

4. Schattenhofer, M. et al. Latitudinal distribution of prokaryotic picoplankton populations in the Atlantic Ocean. Environ. Microbiol. 11, 2078–2093 (2009).

5. Giovannoni, S. J. SAR11 Bacteria: The Most Abundant Plankton in the Oceans. Ann. Rev. Mar. Sci. 9, 231–255 (2017).

6. Giovannoni, S. J. et al. Genome streamlining in a cosmopolitan oceanic bacterium. Science 309, 1242–1245 (2005).

7. Grote, J. et al. Streamlining and core genome conservation among highly divergent members of the SAR11 clade. MBio 3, (2012).

8. Noell, S. E. & Giovannoni, S. J. SAR11 bacteria have a high affinity and multifunctional glycine betaine transporter. Environ. Microbiol. 21, 2559–2575 (2019).

9. Clifton, B. E., Alcolombri, U., Uechi, G.-I., Jackson, C. J. & Laurino, P. The ultra-high affinity transport proteins of ubiquitous marine bacteria. Nature 634, 721–728 (2024).

10. Grzymski, J. J. & Dussaq, A. M. The significance of nitrogen cost minimization in proteomes of marine microorganisms. ISME J. 6, 71–80 (2012).

11. Gibson, D. G. et al. Creation of a bacterial cell controlled by a chemically synthesized genome. Science 329, 52–56 (2010).

12. Lachance, J.-C., Rodrigue, S. & Palsson, B. O. Minimal cells, maximal knowledge. Elife 8, e45379 (2019).

13. Moger-Reischer, R. Z. et al. Evolution of a minimal cell. bioRxiv 2021.06.30.450565 (2021) doi:10.1101/2021.06.30.450565.

14. Cottrell, M. T. & Kirchman, D. L. Transcriptional Control in Marine Copiotrophic and Oligotrophic Bacteria with Streamlined Genomes. Appl. Environ. Microbiol. 82, 6010–6018 (2016).

15. Noell, S. E., Hellweger, F. L., Temperton, B. & Giovannoni, S. J. A reduction of transcriptional regulation in aquatic oligotrophic microorganisms enhances fitness in nutrient-poor environments. Microbiol. Mol. Biol. Rev. 87, e0012422 (2023).

16. Tripp, H. J. et al. SAR11 marine bacteria require exogenous reduced sulphur for growth. Nature 452, 741–744 (2008).

17. Tripp, H. J. et al. Unique glycine-activated riboswitch linked to glycine-serine auxotrophy in SAR11. Environ. Microbiol. 11, 230–238 (2009).

18. Carini, P., Steindler, L., Beszteri, S. & Giovannoni, S. J. Nutrient requirements for growth of the extreme oligotroph ‘Candidatus Pelagibacter ubique’ HTCC1062 on a defined medium. ISME J. 7, 592–602 (2013).

19. Carlson, C. A. et al. Seasonal dynamics of SAR11 populations in the euphotic and mesopelagic zones of the northwestern Sargasso Sea. ISME J. 3, 283–295 (2009).

20. Becker, J. W., Hogle, S. L., Rosendo, K. & Chisholm, S. W. Co-culture and biogeography of Prochlorococcus and SAR11. ISME J. 13, 1506–1519 (2019).

21. Dethlefsen, L. & Schmidt, T. M. Performance of the translational apparatus varies with the ecological strategies of bacteria. J. Bacteriol. 189, 3237–3245 (2007).

22. Roller, B. R. K., Stoddard, S. F. & Schmidt, T. M. Exploiting rRNA operon copy number to investigate bacterial reproductive strategies. Nat Microbiol 1, 16160 (2016).

23. Schwalbach, M. S., Tripp, H. J., Steindler, L., Smith, D. P. & Giovannoni, S. J. The presence of the glycolysis operon in SAR11 genomes is positively correlated with ocean productivity. Environ. Microbiol. 12, 490–500 (2010).

24. Carini, P., White, A. E., Campbell, E. O. & Giovannoni, S. J. Methane production by phosphate-starved SAR11 chemoheterotrophic marine bacteria. Nat. Commun. 5, 4346 (2014).

25. Lankiewicz, T. S., Cottrell, M. T. & Kirchman, D. L. Growth rates and rRNA content of four marine bacteria in pure cultures and in the Delaware estuary. ISME J. 10, 823–832 (2016).

26. Henson, M. W., Lanclos, V. C., Faircloth, B. C. & Thrash, J. C. Cultivation and genomics of the first freshwater SAR11 (LD12) isolate. ISME J. 12, 1846–1860 (2018).

27. Lanclos, V. C. et al. Ecophysiology and genomics of the brackish water adapted SAR11 subclade IIIa. ISME J. 17, 620–629 (2023).

28. Willis, L. & Huang, K. C. Sizing up the bacterial cell cycle. Nat. Rev. Microbiol. 15, 606–620 (2017).

29. Olsson, J. A., Nordström, K., Hjort, K. & Dasgupta, S. Eclipse-synchrony relationship in Escherichia coli strains with mutations affecting sequestration, initiation of replication and superhelicity of the bacterial chromosome. J. Mol. Biol. 334, 919–931 (2003).

30. Levin, P. A., Shim, J. J. & Grossman, A. D. Effect of minCD on FtsZ ring position and polar septation in Bacillus subtilis. J. Bacteriol. 180, 6048–6051 (1998).

31. Sundararajan, K. et al. The bacterial tubulin FtsZ requires its intrinsically disordered linker to direct robust cell wall construction. Nat. Commun. 6, 7281 (2015).

32. Dubarry, N., Willis, C. R., Ball, G., Lesterlin, C. & Armitage, J. P. In Vivo Imaging of the Segregation of the 2 Chromosomes and the Cell Division Proteins of Rhodobacter sphaeroides Reveals an Unexpected Role for MipZ. MBio 10, (2019).

33. Pelletier, J. F. et al. Genetic requirements for cell division in a genomically minimal cell. Cell 184, 2430–2440.e16 (2021).

34. Fujikawa, N. et al. Structural and biochemical analyses of hemimethylated DNA binding by the SeqA protein. Nucleic Acids Res. 32, 82–92 (2004).

35. Boye, E. & Løbner-Olesen, A. The role of dam methyltransferase in the control of DNA replication in E. coli. Cell 62, 981–989 (1990).

36. Blair, J. A. et al. Branched signal wiring of an essential bacterial cell-cycle phosphotransfer protein. Structure 21, 1590–1601 (2013).

37. Krupka, M., Sobrinos-Sanguino, M., Jiménez, M., Rivas, G. & Margolin, W. Escherichia coli ZipA organizes FtsZ polymers into dynamic ring-like protofilament structures. MBio 9, (2018).

38. Pichoff, S., Du, S. & Lutkenhaus, J. Roles of FtsEX in cell division. Res. Microbiol. 170, 374–380 (2019).

39. Corrales-Guerrero, L. et al. MipZ caps the plus-end of FtsZ polymers to promote their rapid disassembly. Proc. Natl. Acad. Sci. U. S. A. 119, e2208227119 (2022).

40. Letzkus, M., Trela, C. & Mera, P. E. Three factors ParA, TipN, and DnaA-mediated chromosome replication initiation are contributors of centromere segregation in Caulobacter crescentus. Mol. Biol. Cell 35, ar68 (2024).

41. Rappe, M. Pelagibacter ubiqueversans gen. nov. sp. nov. Preprint at 10.57973/SEQCODE.R:I44XXWS2 (2025).

42. Oren, A. A plea for linguistic accuracy - also for Candidatus taxa. Int. J. Syst. Evol. Microbiol. 67, 1085–1094 (2017).

43. Cooper, S. & Helmstetter, C. E. Chromosome replication and the division cycle of Escherichia coli B/r. J. Mol. Biol. 31, 519–540 (1968).

44. Jun, S., Si, F., Pugatch, R. & Scott, M. Fundamental principles in bacterial physiology—history, recent progress, and the future with focus on cell size control: a review. Rep. Prog. Phys. 81, 056601 (2018).

45. Skarstad, K., Steen, H. B. & Boye, E. Escherichia coli DNA distributions measured by flow cytometry and compared with theoretical computer simulations. J. Bacteriol. 163, 661–668 (1985).

46. Fu, H., Uchimiya, M., Gore, J. & Moran, M. A. Ecological drivers of bacterial community assembly in synthetic phycospheres. Proc. Natl. Acad. Sci. U. S. A. 117, 3656–3662 (2020).

47. Amin, S. A. et al. Interaction and signalling between a cosmopolitan phytoplankton and associated bacteria. Nature 522, 98–101 (2015).

48. Daniel, R. M. & Danson, M. J. Temperature and the catalytic activity of enzymes: a fresh understanding. FEBS Lett. 587, 2738–2743 (2013).

49. Løbner-Olesen, A., Skarstad, K., Hansen, F. G., von Meyenburg, K. & Boye, E. The DnaA protein determines the initiation mass of Escherichia coli K-12. Cell 57, 881–889 (1989).

50. Bremer, H. & Churchward, G. Deoxyribonucleic acid synthesis after inhibition of initiation of rounds of replication in Escherichia coli B/r. J. Bacteriol. 130, 692–697 (1977).

51. Dai, K. & Lutkenhaus, J. The proper ratio of FtsZ to FtsA is required for cell division to occur in Escherichia coli. J. Bacteriol. 174, 6145–6151 (1992).

52. Luo, H., Csűros, M., Hughes, A. L. & Moran, M. A. Evolution of divergent life history strategies in marine Alphaproteobacteria. MBio 4, (2013).

53. Smith, D. P. et al. Proteomic and transcriptomic analyses of ‘Candidatus Pelagibacter ubique’ describe the first PII-independent response to nitrogen limitation in a free-living Alphaproteobacterium. MBio 4, e00133–12 (2013).

54. Zheng, H. et al. General quantitative relations linking cell growth and the cell cycle in Escherichia coli. Nat Microbiol (2020) doi:10.1038/s41564-020-0717-x.

55. Lee, C. Characterizing Growth Promoters and Inhibitors of SAR11 Pelagibacter sp. HTCC7211. (2013).

56. Braakman, R. et al. Global niche partitioning of purine and pyrimidine cross-feeding among ocean microbes. Sci. Adv. 11, (2025).

57. Monod, J. The Growth of Bacterial Cultures. Annu. Rev. Microbiol. 3, 371–394 (1949).

58. Held, N. A. et al. Nutrient colimitation is a quantitative, dynamic property of microbial populations. Proc. Natl. Acad. Sci. U. S. A. 121, e2400304121 (2024).

59. Schaechter, M., MaalØe, O. & Kjeldgaard, N. O. Dependency on Medium and Temperature of Cell Size and Chemical Composition during Balanced Growth of Salmonella typhimurium. Microbiology 19, 592–606 (1958).

60. Si, F. et al. Mechanistic Origin of Cell-Size Control and Homeostasis in Bacteria. Curr. Biol. 29, 1760–1770.e7 (2019).

61. Guo, X. et al. Automated determination of ammonium at nanomolar levels in seawater by coupling lab-in-syringe with highly sensitive light-emitting-diode-induced fluorescence detection. Molecules 30, (2025).

62. Moran, M. A. et al. The Ocean’s labile DOC supply chain. Limnol. Oceanogr. 67, 1007–1021 (2022).

63. Seymour, J. R., Amin, S. A., Raina, J.-B. & Stocker, R. Zooming in on the phycosphere: the ecological interface for phytoplankton-bacteria relationships. Nat Microbiol 2, 17065 (2017).

64. Paerl, H. W. Why does N-limitation persist in the world’s marine waters? Mar. Chem. 206, 1–6 (2018).

65. Sarmento, H. & Gasol, J. M. Use of phytoplankton-derived dissolved organic carbon by different types of bacterioplankton: Use of phytoplankton-derived DOC by bacterioplankton. Environ. Microbiol. 14, 2348–2360 (2012).

66. Brüwer, J. D. et al. In situ cell division and mortality rates of SAR11, SAR86, Bacteroidetes, and Aurantivirga during phytoplankton blooms reveal differences in population controls. mSystems 8, e0128722 (2023).

67. Margolin, W. FtsZ and the division of prokaryotic cells and organelles. Nat. Rev. Mol. Cell Biol. 6, 862–871 (2005).

68. Barrows, J. M., Sundararajan, K., Bhargava, A. & Goley, E. D. FtsA regulates Z-ring morphology and cell wall metabolism in an FtsZ C-terminal linker-dependent manner in Caulobacter crescentus. J. Bacteriol. 202, (2020).

69. Wu, K. J. et al. Characterization of Conserved and Novel Septal Factors in Mycobacterium smegmatis. J. Bacteriol. 200, (2018).

70. Oh, H.-M. et al. Complete genome sequence of ‘Candidatus Puniceispirillum marinum’ IMCC1322, a representative of the SAR116 clade in the Alphaproteobacteria. J. Bacteriol. 192, 3240–3241 (2010).

71. Coelho, J. T. et al. Culture-supported ecophysiology of the SAR116 clade demonstrates metabolic and spatial niche partitioning. bioRxiv (2025) doi:10.1101/2025.01.06.631592.

72. Cho, J.-C. & Giovannoni, S. J. Parvularcula bermudensis gen. nov., sp. nov., a marine bacterium that forms a deep branch in the alpha-Proteobacteria. Int. J. Syst. Evol. Microbiol. 53, 1031–1036 (2003).

73. Dang, H., Li, T., Chen, M. & Huang, G. Cross-ocean distribution of Rhodobacterales bacteria as primary surface colonizers in temperate coastal marine waters. Appl. Environ. Microbiol. 74, 52–60 (2008).

74. Follows, M. J., Dutkiewicz, S., Grant, S. & Chisholm, S. W. Emergent biogeography of microbial communities in a model ocean. Science 315, 1843–1846 (2007).

75. Aumont, O., Ethé, C., Tagliabue, A., Bopp, L. & Gehlen, M. PISCES-v2: an ocean biogeochemical model for carbon and ecosystem studies. Geosci. Model Dev. 8, 2465–2513 (2015).

76. Stock, C. A. et al. Ocean biogeochemistry in GFDL’s Earth system model 4.1 and its response to increasing atmospheric CO_2_. J. Adv. Model. Earth Syst. 12, (2020).

77. Ross, A. C. et al. A high-resolution physical–biogeochemical model for marine resource applications in the northwest Atlantic (MOM6-COBALT-NWA12 v1.0). Geosci. Model Dev. 16, 6943–6985 (2023).

78. Martinez-Gutierrez, C. A., Uyeda, J. C. & Aylward, F. O. A timeline of bacterial and archaeal diversification in the ocean. Elife 12, (2023).

79. Hyun, J. C. & Palsson, B. O. Reconstruction of the last bacterial common ancestor from 183 pangenomes reveals a versatile ancient core genome. Genome Biol. 24, 183 (2023).

80. Staley, J. T. & Konopka, A. Measurement of in situ activities of nonphotosynthetic microorganisms in aquatic and terrestrial habitats. Annu. Rev. Microbiol. 39, 321–346 (1985).

81. Henson, M. W. et al. Expanding the Diversity of Bacterioplankton Isolates and Modeling Isolation Efficacy with Large-Scale Dilution-to-Extinction Cultivation. Appl. Environ. Microbiol. 86, (2020).

82. Paysan-Lafosse, T. et al. InterPro in 2022. Nucleic Acids Res. 51, D418–D427 (2023).

83. Jones, P. et al. InterProScan 5: genome-scale protein function classification. Bioinformatics 30, 1236–1240 (2014).

84. Thrash, C. Pangenomic analyses files. figshare 10.6084/M9.FIGSHARE.30087295.V1 (2025).

85. UniProt Consortium. UniProt: The universal protein knowledgebase in 2025. Nucleic Acids Res. 53, D609–D617 (2025).

86. Ahmad, S. et al. The UniProt website API: facilitating programmatic access to protein knowledge. Nucleic Acids Res. 53, W547–W553 (2025).

87. Rappé, M. S., Connon, S. A., Vergin, K. L. & Giovannoni, S. J. Cultivation of the ubiquitous SAR11 marine bacterioplankton clade. Nature 418, 630–633 (2002).

88. Henson, M. W. et al. Artificial Seawater Media Facilitate Cultivating Members of the Microbial Majority from the Gulf of Mexico. mSphere 1, (2016).

89. Sodium phosphate. Cold Spring Harb. Protoc. 2006, db.rec8303 (2006).

90. Cheng, C. & Thrash, J. C. sparse-growth-curve: a Computational Pipeline for Parsing Cellular Growth Curves with Low Temporal Resolution. Microbiol Resour Announc 10, (2021).

91. Lanclos, V. C. et al. New isolates refine the ecophysiology of the Roseobacter CHAB-I-5 lineage. ISME Commun. (2025) doi:10.1093/ismeco/ycaf068.

92. Cheng, C. Thrash-lab/SAR11_cell_cycle: V1.0.0. (Zenodo, 2025). doi:10.5281/ZENODO.17703344.

93. Stokke, C., Flåtten, I. & Skarstad, K. An easy-to-use simulation program demonstrates variations in bacterial cell cycle parameters depending on medium and temperature. PLoS One 7, e30981 (2012).

94. Michelsen, O., Teixeira de Mattos, M. J., Jensen, P. R. & Hansen, F. G. Precise determinations of C and D periods by flow cytometry in Escherichia coli K-12 and B/r. Microbiology 149, 1001–1010 (2003).

95. Thrash, C. Epimicroscopy images of SAR11. figshare 10.6084/M9.FIGSHARE.29396375.V1 (2025).

96. Messer, W. The bacterial replication initiator DnaA. DnaA and oriC, the bacterial mode to initiate DNA replication. FEMS Microbiol. Rev. 26, 355–374 (2002).

97. Katayama, T., Ozaki, S., Keyamura, K. & Fujimitsu, K. Regulation of the replication cycle: conserved and diverse regulatory systems for DnaA and oriC. Nat. Rev. Microbiol. 8, 163–170 (2010).

98. Boye, E. & Løbner-Olesen, A. Bacterial growth control studied by flow cytometry. Res. Microbiol. 142, 131–135 (1991).

99. Stokke, C., Waldminghaus, T. & Skarstad, K. Replication patterns and organization of replication forks in Vibrio cholerae. Microbiology 157, 695–708 (2011).

