## Supplementary Text and Figures for "Cell cycle dysregulation of globally important SAR11 bacteria resulting from environmental perturbation"

**Supplementary Notes** 1

Summary 2

1. Detailed evaluation of cell cycle regulation genes in SAR11 2

2. Bacterial cell cycles and the Cooper-Helmstetter Index 3

3. Reanalysis of previous SAR11 ploidy data 4

4. Automated processing of flow cytometry data files 5

5. DNA replication "run-off" experiments 6

6. Additional discussion points 8

Caveats regarding excessive nutrients 8

Temperature vs. elemental stochiometry 8

Implications for activity measurements 9

Nitrogen-sensing genes 9

Polyploidy as a possible mechanism for enhanced recombination 9

**Supplementary Figures Captions** 10

**Supplementary Tables Descriptions** 12

**Supplementary References** 13

### Supplementary Notes

#### Summary

This supplementary material provides foundational background on bacterial cell cycles and validation of our analytical approaches. **Section 1** is an expanded description of our findings of cell cycle regulation genes in SAR11. The corresponding section in the main text is an abbreviated version. **Section 2** introduces the Cooper-Helmstetter Index as a quantitative measure of cell cycle coordination, defining it as the ratio between maximum and minimum ploidy levels in bacterial populations. We demonstrate through reanalysis of published data from *E. coli* and *Vibrio cholerae* that coordinated cell cycles consistently maintain this ratio at approximately 2:1. **Section 3** reanalyzes legacy SAR11 data from Carini et al. 2013[1], revealing early evidence of cell cycle disruption under nutrient stress that supports our current findings. **Section 4** describes our automated flow cytometry data processing pipeline, which enables high-throughput analysis of ploidy distributions while maintaining consistency with manual analysis methods. **Section 5** presents DNA replication "run-off" experiments using antibiotic treatments to validate our measurements of cell cycle period durations and confirm the mechanisms underlying nitrogen-induced cell cycle disruption in SAR11. **Section 6** contains additional discussion points stemming from our results.

#### 1. Detailed evaluation of cell cycle regulation genes in SAR11

We searched for cell cycle regulation genes in SAR11 using a previous pangenome of SAR11 with 470 representatives[2], as well as searches in UniProt to compare the prevalence of these genes in a wider diversity of related bacteria in the Alphaproteobacteria and Pseudomonadota more broadly. As a positive reference, we examined the universal initiator protein DnaA, which binds *oriC* to trigger chromosome replication[3,4]. DnaA was present in nearly all SAR11 genomes, although absent in 45 of 470 (9.6%), likely reflecting incomplete genomes rather than true gene losses. We also used UniProt to provide context for the relative distribution of proteins at different levels of evolutionary distance from SAR11. DnaA was found in 50,897 different bacterial taxa and all 120 SAR11 genomes on UniProt **(Supplementary Table Tab 3)**.

Several regulators of replication initiation were absent across SAR11. We did not find evidence for homologs of SeqA, which binds the origin to prevent early re-initiation of chromosome replication[5], in any SAR11 genome **(Supplementary Table Tab 5)**. Dam (a methyltransferase that maintains the methylation pattern at the origin that SeqA recognizes[6]) was absent among our 470 SAR11 genomes, but was present in one MAG **(Supplementary Table Tab 6)** out of the 120 DnaA-containing SAR11 genomes on UniProt. The deletion or mutation of SeqA or Dam in *E. coli* causes dysregulated initiation and abnormal chromosome copy numbers, such as three to five chromosome equivalents[7,8]. SeqA was largely absent from Alphaproteobacteria: only 24 Alphaproteobacteria genomes had an annotated SeqA among 2,254 Pseudomonadota genomes on UniProt **(Supplementary Table Tab 5)**. However, the Dam protein had 1,767 annotations within the Alphaproteobacteria and 24,909 across the Pseudomonadota

**(Supplementary Table Tab 6)**, so its absence in SAR11 is unusual. We also examined ChpT, a phosphotransferase that links cell cycle signaling to replication initiation in *C. crescentus*[9]. Although ChpT was primarily within the Alphaproteobacteria (3,971 annotations in the Alphaproteobacteria of the 4,001 in the Pseudomonadota), we similarly did not find any evidence of this protein in in our SAR11 pangenome analysis and only found one example in the UniProt SAR11 genomes **(Supplementary Table Tab 7)**.

Genes essential for divisome assembly and positioning were also broadly missing in SAR11. ZipA, a membrane anchor that stabilizes FtsZ protofilaments for septum formation[10], had no homologs in SAR11 genomes. In *E. coli*, deletion of ZipA causes filamentation and chromosome missegregation[11]. ZipA was primarily found in Gammaproteobacteria (6,581 of 9,090 annotations were in Gammaproteobacteria - **Supplementary Table Tab 8**), but we also searched for a more universally distributed Z-ring protein, FtsE (2,779 annotations within Alphaproteobacteria out of 8,189 in the Pseudomonadota). FtsE is an integral Z-ring protein important for septum formation[12]. In *C. crescentus*, knocking out FtsE results in inhibition of newborn cell separation[13]. Similarly, we found no FtsE homologs in the SAR11 pangenome and only one in a freshwater SAR11 MAG on UniProt **(Supplementary Table Tab 9)**.

We also examined regulators involved in chromosome segregation. In *B. subtilis*, deletion of *minC* and *minD*, two universally distributed genes **(Supplementary Table Tabs 10, 11)**, leads to misplaced septa, abnormal chromosome segregation, and polyploidy[14]. Homologs for MinC, an inhibitor of FtsZ polymerization[15], and MinD, its ATPase partner that oscillates between cell poles[16], were almost entirely absent: we did not find MinC in any SAR11 genomes, and detected MinD in only a very minor subset of genomes (≤ 4 genomes per subclade) **(Supplementary Table Tabs 1, 3)**. Homologs for MipZ, an Alphaproteobacteria-biased ATPase (2,905/3,957 bacterial hits were in Alphaproteobacteria on UniProt- **Supplementary Table Tab 12**) that positions the Z-ring by forming gradients from chromosome origins in *C. crescentus*[17] and *C. sphaeroides*[18], were also extremely rare in SAR11 (three in subclade Ia, one in subclade Ib, one found on UniProt - **Supplementary Table Tab 1**). SepF, a membrane-associated bundling factor for FtsZ[19], and LemA, a poorly characterized divisome regulator[20], were absent across our SAR11 pangenome and were only identified in one freshwater SAR11 MAG on UniProt **(Supplementary Table Tabs 13, 14)**. Experiments with the synthetic minimal cell JCVI-syn3A identified both SepF and LemA as essential for maintaining uniform cell sizes, with knockouts in their genes producing polymorphic populations due to failures in cell division[20]. We corroborated that TipN, a polarity factor for *C. crescentus*[21], was largely restricted to the Caulobacterales **(Supplementary Table Tab 16)**, but only had nine hits in the SAR11 pangenome **(**although these were poor quality, see Methods, **Supplementary Table Tabs 1,3,15)**.

### 2. Bacterial cell cycles and the Cooper-Helmstetter Index

The coordination between chromosome replication and cell division was first characterized as the Cooper-Helmstetter model in the 1960s[22]. Though many later studies have updated the details of the model, one fundamental principle that has empirically worked well for decades to model the average population behavior of nutrient shifts in wildtype *E. coli*[23,24] and *Vibrio* *cholera*[25] is the 2:1 ratio between the maximum and minimum ploidy level **(Fig. S1)**. In bacterial cell cycles, each newborn cell gradually duplicates its cellular DNA, then evenly distributes it to the two children cells through cell division. Therefore, if we look at the ploidy level distribution of a growing population with coordinated cell cycles, we will see a 2:1 ratio between the maximum ploidy level (in the cells that are about to divide) and the minimum ploidy level (in the newborn cells). This 2:1 ratio does not necessarily mean 2 chromosomes vs. 1 chromosome, as cells with overlapping cell cycles **(Fig. S1)** would be duplicating DNA from 1X to 2X with X >> 1 chromosome equivalent[26], and the concept still applies to multi-chromosome taxa like *Vibrio*.

For the current study, we refer to the ratio between maximum and minimum ploidy levels as the "Cooper-Helmstetter Index" in recognition of Stephen Cooper and Charles Helmstetter's foundational contributions to understanding DNA replication dynamics[22]. In theory, a coordinated population would yield a Cooper-Helmstetter Index of 2. However, ploidy level measurements are often influenced by noise, likely due to variations in DNA staining and signal transmission noise in flow cytometry[23]. As a result, the ploidy level distribution data presented in this study, including our reanalysis of previously published data, represent theoretical signals convoluted with Gaussian noise[23]. To account for this noise in the real data, we applied a cutoff percentage (X%) and calculated the ratio between the (100-X)% and X% percentiles **(Fig. S2–S6)**.

The reference studies present the ploidy level distributions in histograms[1,23,25,27]. We downloaded or screenshotted the histograms and extracted the contour using the online app PlotDigitizer (plotdigitizer.com/app).

We calculated the Cooper-Helmstetter Index for *E. coli*[23] and *V. cholerae*[25] under varying nutrient conditions, finding that the Index consistently remained around 2 with a 10% cutoff **(Figs. S2, S3)**. Additionally, we examined the variation in ploidy level distribution across exponential and stationary phases for the same *E. coli* culture[27]. Despite fluctuations in the ploidy level distribution, the Cooper-Helmstetter Indices remained stable around 2 with the 10% cutoff **(Fig. S5)**. These results suggest that for these copiotrophic organisms, chromosome replication and cell division are well-coordinated across different nutrient conditions, with cells dividing as their DNA doubles **(Fig. S1)**. Previous reviews focused on many complex aspects of bacterial cell cycles, but always took this coordination for granted[26,28]. However, we argue that this coordination is non-trivial, even for model organisms, and is notably absent in SAR11 bacteria. Studies of mutants in model organisms have highlighted key marker genes that regulate this coordination, as well as the altered ploidy level distributions that occur when this coordination is disrupted (below).

### 3. Reanalysis of previous SAR11 ploidy data

We also extracted legacy data from prior SAR11 research[1]. These data were collected during stationary phase under varying pyruvate and glycine concentrations, enabling comparison with our data in **Extended Data Fig. 1**. The ploidy data from Carini et al. 2013[1] include histograms of DNA fluorescence signals obtained via flow cytometry and epifluorescence microscopy images **(Fig. S6)**. The detailed Python code for the analysis of the fluorescence microscopy images can be found on our GitHub repository[29]. In brief, we downloaded the microscopic images in jpg format from the reference[1]. Cells were segmented using the Otsu method, and their area was calculated by counting the pixels of each connected segment area. This image processing method was implemented through OpenCV (4.10.0). We then highlighted the cells with large volumes (larger than 20 pixels) in purple **(Fig. S6)**.

As noted in the methods section of our main text, we employed an improved DNA staining protocol that reduces variation in ploidy level measurements for SAR11, whereas the reference study used the original, less optimized protocol. Furthermore, the histograms in the reference study were plotted on a log scale, introducing more noise compared to the ploidy-level data presented in our study. Despite this increased noise, we observed that in the legacy data under low pyruvate (carbon) and high glycine (nitrogen) conditions, the Cooper-Helmstetter Index increased, with a significantly higher fraction of diploids (**Fig. S6a-d**). This observation aligns with our data in **Fig. 3** and supports the conceptual model we propose in **Fig. 5c**.

In the legacy epifluorescence microscopy images under fixed pyruvate concentrations and varying glycine concentrations (**Fig. S6f-j**) we observed a larger number of diploids under lower glycine concentrations (**Fig. S6f,g**) compared to higher glycine concentrations (**Fig. S6h,i**). This seems to contradict our findings, as we propose in our study that excessive nitrogen disrupts the SAR11 cell cycle leading to increased frequencies of diploids and polyploids. We believe this discrepancy can be explained in a few ways. First, the AMS1 medium used in the reference study contained 400 µM ammonium, which could disrupt the cell cycle as we showed in **Fig. 4**. Second, the microscopic images were collected in stationary phases, and in our work we also observed higher fractions of diploid cells during stationary phases in our positive control **(Fig. 4e, Extended Data Fig. 2a)**, though the underlying mechanisms remain unclear. Regardless, the reanalyses of legacy microscopic images still show some consistency with our results: compared to samples where [glycine] > [pyruvate] (**Fig. S6a-e**), we observed fewer filamentous cells or diploids in samples than when [pyruvate] > [glycine] (**Fig. S6f-j**). This is consistent with our findings that excessive glycine over pyruvate disrupts cell divisions.

### 4. Automated processing of flow cytometry data files

To handle the large volume of flow cytometry samples generated in this study, we developed an automated pipeline for processing FCS (Flow Cytometry Standard) files and extracting ploidy level distributions (**Fig. S7**). This approach maintained consistency with manual analysis performed on the BD Accuri software while enabling high-throughput processing.

We used the Python ‘fcsparser’ package to read and parse raw flow cytometry files (.fcs format). The automated pipeline begins by generating scatter density plots of fluorescence channels (**Fig. S7a,b**). Note that while our flow cytometry software channels are labeled FITC (533 ± 30 nm) and PE (585 ± 40 nm) based on standard fluorophore wavelengths, we did not use fluorescein isothiocyanate or phycoerythrin in our experiments— the FITC and PE labels simply correspond to the green and yellow fluorescence detection channels.

The gating process involves analyzing the ratios of yellow versus green fluorescence intensities alongside green fluorescence intensities versus areas (**Fig. S7c**). Events are identified as cells based on their green versus yellow fluorescence ratio (3.5–6:1, corresponding to SYBR Green-stained DNA), with cells highlighted in the density scatter plots (**Fig. S7d,e**).

Following cell identification, we extract ploidy level distributions through a multi-step process. First, we generate ridgeline plots showing the distribution of green fluorescence height versus area ratios (FITC-H/FITC-A) for each rounded fluorescence value (**Fig. S7f**). The peak of each distribution corresponds to populations with one chromosome equivalent, identified and marked with circles and red dashed lines.

These aligned distributions are then stacked together in chromosome equivalents (**Fig. S7g**), and their sum produces the final ploidy level distribution (**Fig. S7h**). The relative maximum abundance is normalized to 1, and we apply Gaussian kernel density estimation using the 'gaussian_kde' function from Scipy (v1.14.1) with a bandwidth of 0.05 to smooth the distribution.

The Cooper-Helmstetter Index is calculated from the normalized ploidy distributions (**Fig. S7i-k**). We generate cumulative distribution plots with colored right angles corresponding to different percentile ranges: 5%–95% (5% cutoff), 10%–90% (10% cutoff), 15%–85% (15% cutoff), and 20%–80% (20% cutoff) (**Fig. S7j**). The Cooper-Helmstetter Index is then calculated as the ratio between the upper and lower bounds of these percentile ranges (**Fig. S7k**).

For samples with coordinated cell cycles (Cooper-Helmstetter Index ≈ 2), we estimate B, C, and D periods using analytical methods based on previous studies[23,24]. This involves theoretical age distribution modeling where cell age of 1 represents cells about to divide and 0 represents newborn cells, following the equation $p(a) = 2(1-a)\ln(2)$[30] (**Fig. S7l**). We generate predicted distributions for subpopulations in each cell cycle period (B, C, D), with their sum representing the theoretical ploidy distribution (**Fig. S7m**). To verify our analytical approach, we perform numerical simulations, growing populations from 1 cell to $10^{11}$ cells with variation in generation times to smooth early growth fluctuations (**Fig. S7n-p**).

The automated pipeline showed excellent agreement with manual analysis, ensuring reliability for high-throughput processing. The complete Python implementation is available in our GitHub repository[29] as 'FCS_analysis.ipynb', providing a reproducible workflow for similar flow cytometry-based cell cycle studies.

### 5. DNA replication "run-off" experiments

Batch cultures actively growing in large volumes (> 500 mL) were aliquoted in triplicate into 10 mL falcon tubes when the cell density achieved > 5 x $10^6$ cells/mL. One of the 10 mL aliquots was left antibiotics-free as the positive control while the other two were given antibiotic treatments. Cephalexin, rifampin, and chloramphenicol were pre-made in 3,000 µg/mL concentrated stocks and syringe filter-sterilized. Concentrated stocks were then spiked in the corresponding falcon tubes. Experiments for cell wall inhibition only received 9 µg/mL cephalexin. Experiments with all antibiotic treatments included 9 µg/mL cephalexin, 6 µg/mL rifampin, and 6 µg/mL chloramphenicol.

We treated populations grown in different ammonium concentrations with two different antibiotic treatments designed to further tease apart the cell cycle processes most affected by high nitrogen **(Fig. S8)**. The first treatment was with cephalexin only, which inhibits cell wall synthesis[31], and thus cell division, but doesn't affect the other aspects of the cell cycle. The other treatment was a combination of cephalexin, chloramphenicol (inhibits translation), and rifampin (inhibits transcription)[32,33], which contrasts the first treatment by additionally preventing cells from producing new transcripts and proteins. In both treatments, cell growth ceased, confirming effective inhibition **(Fig. S8bc)**.

We closely tracked ploidy level distribution across a one-day period **(Fig. S8i–n)**, calculated the per-cell DNA content **(Fig. S8d–f)**, and compared the differences between samples with and without antibiotic treatments **(Fig. S8gh)**. We observed further evidence of cell division falling behind DNA replication with excessive nitrogen as we found in **Fig. 4k**. Without antibiotics, cells in high ammonium showed an increase in DNA per cell with time, whereas in no ammonium, DNA levels remained constant (**Fig. S8d**). This phenomenon is consistent with continued DNA replication without cell division.

In antibiotics-treated samples, we verified the cell cycle parameters measured in **Fig. 4k**. Under cell division inhibition (cephalexin-only treatment), the 1-chromosome population gradually transformed into higher ploidy levels and mostly disappeared within 20 hours **(Fig. S8j,m)**, consistent with our previously measured sum of the B and C periods at 20 hours (**Fig. 4k)**. Ploidy level distribution stabilized after 0.5 days under both antibiotic treatments, confirming our quantification of the C period at 10 – 12 hours (**Fig. 4k)**. This showed that even when aneuploidy occurred, DNA replication was mostly unvaried.

However, our observations were biased toward the cells with either one or two chromosome equivalents, since these dominated the total population. Despite the challenge of directly observing the polyploids, we did find evidence that their physiology was substantially different from the mono- and diploids. Diploids did not exponentially synthesize DNA, as only one chromosome would replicate. Triploids emerged in cephalexin-treated samples after 20 hours **(Fig. S8j)**, matching the 20-hour B and C period sum **(Fig. 4k)**. This suggests that 3-chromosome cells developed from 2-chromosome cells (as opposed to directly from single-chromosome cells), reflecting non-exponential DNA synthesis whereby only one chromosome was replicating, and consistent with the presence of cells with odd chromosome numbers in our nutrient enrichment experiments **(Fig. 4h)**. Additionally, polyploid cells exhibited a slowdown in DNA synthesis, particularly in cephalexin-only treated samples with high ammonia **(Fig. S8g)**, contrary to expectations that DNA synthesis would continue at the same rate given that cephalexin does not directly inhibit DNA replication. This indicates that polyploids had distinct physiology as the monoploids.

### 6. Additional discussion points

#### *Caveats regarding excessive nutrients*

Very high concentrations of ammonia or glycine (50–500 mM - much higher than the concentrations we tested on SAR11 - **Fig. 3a-f, Extended Data Fig. 2**) can inhibit growth of *E. coli* and *B. subtilis*, however, the mechanisms of inhibition in SAR11 are different. The inhibitory effect of ammonium in *E. coli, B. subtilis*, and *Corynebacterium glutamicum* was characterized as an outcome of increased ionic strength[34], which does not correspond to our observations in SAR11 given that HTCC1062 was already growing in much higher ionic strength media and our concentrations of ammonium were much lower. Similarly to what we observed for SAR11, excessive ammonia caused aneuploidy and filamentous cells in *C. crescentus*[35], and excessive glycine inhibited cell wall synthesis and cell division in *E. coli, B. subtilis, Selenomonas ruminantium, Streptococcus bovis,* and others[36]. However, our work also identified additional environmental perturbations, such as temperature or carbon shifts, that have analogous effects compared to excessive nitrogen.

The nutrient concentrations in our experiments were in the high micro- to low millimolar scale, because we needed enough cell density (above 1 million cells per milliliter) for good DNA staining and reliable ploidy level distribution measurements. Although the open ocean standing stocks of many of our nutrient compounds, such as pyruvate, ammonium chloride and amino acids are below the nanomolar scale[37,38] and often undetected, this can be due to their rapid turnover through biogeochemical processes[39], as concentrations in the local phycosphere around phytoplankton cells can be much greater[40]. Indeed, SAR11 requires extra nutrient amendment to grow when cultured with natural seawater[41], implying that the standing concentrations measured in the open ocean may not actively reflect the local concentrations experienced by SAR11 cells during growth. While the exact ambient concentrations are challenging to measure and remain poorly characterized, synthetic phycosphere studies have used 3 mM ammonium and 7.5 mM total organic carbon to mimic the microenvironment surrounding dinoflagellates or diatoms[42]. It's plausible that the concentrations inhibiting SAR11 growth occur in local environments near phytoplankton cells. But we note again that nutrient concentrations were not the only perturbation causing cell cycle dysregulation.

#### *Temperature vs. elemental stochiometry*

Our observations of aneuploidy occurring at temperatures above 20°C under a 1:1 pyruvate-to-glycine ratio (~5:1 C:N), but not in samples maintained below 20°C (**Fig. 3d, Extended Data Fig. 1e**) are notable given that the cellular C:N ratio of HTCC1062 is approximately 4–6:1[43], and the strain was originally isolated from ~12 °C seawater[41]. The emergence of coordinated cell cycles at lower temperatures under nutrient conditions resembling cellular stoichiometry suggests that such environments may more closely reflect SAR11's native ecological and physiological context.

##### *Implications for activity measurements*

SAR11 sensitivity to nutrient perturbations may also affect estimates of *in situ* bacterial activity. SAR11 cannot incorporate thymidine in common DNA synthesis activity assays[43], and our results show that adding 1 mM leucine to our positive control media (CCM2) inhibited SAR11 growth (**Extended Data Fig. 2d, Supplementary Table Tab 17**). Such assays are used with concentrations as low as 20 nM[44], suggesting that the inhibitory effects we observed might be unlikely. However, as we also found that relative C:N ratios and carbon substrate changes can significantly perturb the cell cycle. In that context, since the ambient organic carbon concentrations of natural sea water could be as low as nanomolar levels, 20 nM of leucine might be significant enough to underestimate SAR11's *in situ* activity, so we feel this warrants further investigation. Furthermore, recent measurements showing SAR11 single-cell respiration rates at very low levels relative to copiotrophic taxa may also have underestimated SAR11 respiration since many of the samples were taken during phytoplankton blooms[45], which we also expect to cause significant cell cycle disruption due to decreased C:N ratios and carbon turnover. Opposite these scenarios where *in situ* activity may be underestimated, our identification of compounds causing growth inhibition without aneuploidy (category 2 in **Fig. 3g**) reveals that SAR11 can undergo complete growth arrest under certain conditions, potentially representing a dormancy-like state[46]. For example, excessive alanine caused complete cessation of both population growth and DNA replication, while cells maintained normal ploidy levels throughout (4-8 mM, **Fig. 3f**). This contrasts with the chaotic cell cycle disruption caused by an excess of other nitrogen-containing compounds in excess, where DNA replication continued while cell division failed, leading to aneuploidy and eventual cell death. Our findings thus have broader implications for understanding the metabolic contributions of streamlined marine bacteria to ocean biogeochemistry, as standard activity measurements may not accurately capture their physiological state or ecological function.

##### *Nitrogen-sensing genes*

Filamentation of *C. crescentus* induced by spent media and ammonia includes upregulation of nitrogen metabolism genes and downregulation of cell cycle regulatory genes[35]. Notably, the presence of additional P-II genes in LSUCC0530 did not prevent this organism from experiencing cell cycle dysregulation vulnerabilities (**Fig. 6**). This indicates that cell cycle regulatory genes are more critical than nutrient sensing systems for maintaining division coordination under stress, and their absence represents deeper architectural constraints of extreme genome reduction. This may explain why P-II regulatory systems are absent in most SAR11 subclades other than subclade III **(Supplementary Table Tab 3)**: nutrient sensing provides little benefit when the fundamental cell cycle machinery cannot respond appropriately to environmental changes, and thus these genes are ripe for deletion with continued streamlining.

##### *Polyploidy as a possible mechanism for enhanced recombination*

Regardless of the evolutionary path, the absence of cell cycle regulating genes has compromised SAR11's ability to maintain homeostasis across environmental conditions, constraining steady-state growth to a narrow range of conditions unlike the broad regulatory flexibility of model copiotrophic organisms. However, this apparent evolutionary constraint may paradoxically provide population-level benefits through enhanced genetic diversification. The formation of polyploid cells under environmental stress could facilitate genetic exchange through two potential mechanisms. First, the presence of multiple chromosome copies within individual cells could enable intracellular recombination events between homologous chromosomes, potentially contributing to the exceptionally high rates of recombination observed in natural SAR11 populations[47,48]. When these polyploid cells subsequently undergo lysis, as indicated by our flow cytometry data showing decreased cell integrity during stationary phase (**Extended Data Fig. 7**), the resulting DNA fragments containing novel recombinant sequences could be released and taken up by naturally competent surviving cells, as observed in other bacterial systems[49,50]. Second, some polyploid cells may retain viability and undergo division to release progeny, similar to the phenomenon observed in filamentous *Caulobacter crescentus* cells[35]. Both pathways—whether through DNA release from lysing cells or progeny from dividing filamentous cells—could increase genetic diversity within SAR11 populations. This suggests that SAR11's cell cycle vulnerabilities may represent another evolutionary trade-off inherent to extreme genome streamlining: reduced regulatory robustness in exchange for enhanced opportunities for genetic diversification that could facilitate long-term adaptation in the oligotrophic marine environment.

### Supplementary Figures Captions

**Supplementary Figure 1|** Illustration of non-overlapping and overlapping chromosome replication cycles. In non-overlapping cycles, the generation time $\tau$=B+C+D. There is only one ongoing replication fork within each generation. In overlapping cycles, C+D > $\tau$. There are multiple replication forks within one generation -- as some periods of C0 vs. C1, D0 vs. C1, C1 vs. C2, and D1 vs. C2, are overlapping.

**Supplementary Figure 2|** Reanalysis of the ploidy level distributions from a previous *E. coli* B/r strain study[23]. Each row represents a cultivation condition (batch/chemostat, carbon source, doubling time $\tau$), with the histogram ploidy level distribution (left), cumulative distribution of the ploidy levels (middle), and the Cooper-Helmstetter Index calculated across different percentage cutoffs (right). For the left column plots, we used dual y-axes to reflect the rescaling of the original raw data in red to the relative frequencies in black. The cumulative distribution plots also highlight the region of X — 100-X percentiles under different X% cutoffs via right angles with different colors. The colors representing different X% cutoffs match the cumulative distribution plots and the Cooper-Helmstetter Index plots.

**Supplementary Figure 3|** Reanalysis of *V. cholera* 2740-80 (CTX$\varphi^-$)[51] ploidy level distribution data[25] in the same format as **Fig. S2**.

**Supplementary Figure 4|** Reanalysis of the ploidy level distribution of genetic mutants from *E. coli K-12*[8] in a similar format as **Fig. S2**. The genotypes are specified on the left side of the plots. Two nutrient conditions were used for cultivation: glucose only and glucose with casamino acid (cAA). For some of the conditions, doubling time was measured, which is labeled as $\tau_{Glucose}$ or $\tau_{cAA}$.

**Supplementary Figure 5|** Variation of the ploidy level distributions and the Cooper-Helmstetter Index (10% cutoff) of *E. coli K-12* AB1157 under glucose minimal media[27] over time. The ridge lines were adapted from Fig. 5 in the reference, where red with low OD values represents exponential phase while blue with OD 1.8 represents stationary phase.

**Supplementary Figure 6|** Reanalysis of the microscopic images of SYBR-green stained cells and the histograms of the ploidy level distribution from a previous SAR11 HTCC1062 study[1]. The original data were from Fig. S2 in the referenced paper, with the scale bars missing. The labels for the plots (A – J) match the original reference figure. For the microscopic images we highlighted the cells with large cell volumes in purple to contrast with green for the rest**.**

**Supplementary Figure 7|** Detailed data analysis process of flow cytometry data. **(a,b)** Scatter density plots of FITC-H vs. PE-H and FITC-A channels. **(c)** Histograms of yellow vs. green fluorescence intensity ratios (orange) and green fluorescence intensities vs. areas. **(d,e)** Density scatter plots with identified cellular events highlighted in orange and blue. **(f)** Ridgeline plots of green fluorescence height vs. area ratios with chromosome equivalent peaks marked. **(g)** Aligned distributions in chromosome equivalents. **(h)** Final normalized ploidy level distribution. **(i-k)** Cooper-Helmstetter Index calculation process showing normalized distribution (the normalization process was reflected by rescaling the relative abundance data in red to relative frequency in black), cumulative distribution with percentile ranges, and calculated indices. **(l,m)** Analytical estimation of B, C, and D periods showing theoretical age distribution and predicted subpopulation distributions. **(n-p)** Numerical simulation verification including simulated growth curve, cell age distribution comparison, and experimental vs. simulated ploidy distributions.

**Supplementary Figure 8| Antibiotic treatments point to the mechanisms of growth limitation by nitrogen.** All experiments- no ammonium (blue) vs. 4 mM ammonium (yellow). **(a-c)** 30-hour growth curves of the positive control (no antibiotics) and two antibiotic conditions. **(d-f)** Variation of chromosome equivalents per cell with time, according to the antibiotic condition in a-c. The sample mean ploidy levels are quantified by integrating the histograms in Fig. (i)--(n). In (d), the constant ploidy level for the 0 ammonia samples indicates that the cultures were under steady state growth while the increasing ploidy level for the 4mM ammonia samples shows the continuous accumulation of aneuploidy. **(i-n)** Ploidy level distributions of the samples in different combinations of ammonia concentrations and antibiotic treatments. **(g)** The difference in mean cellular ploidy level between cell wall synthesis inhibition (cephalexin-treated) samples and the positive control (no antibiotics). **(h)** Same calculation as in (g), but subtracting the positive control from the samples with the combined rifampin (transcription), chloramphenicol (translation), and cephalexin (cell wall synthesis) treatment. The lower plateau for the 4mM curve in both (h) and (g) indicates less activity for the aneuploid cells.

### Supplementary Table Captions

**1-2. Source Data of Fig. 1| Distributions of canonical cell cycle regulatory proteins across SAR11 subclades based on our pangenomic analysis (left) and within selected bacterial taxa on UniProt (right).** The DnaA protein serves as a positive reference that was expected to be present in all or at least most of the genomes. In the SAR11 pangenomic analyses section, the number of genomes with the protein out of the total number of genomes from each subclade is shown. White boxes indicate complete absence of the protein, while gray boxes show partial presence. Detailed presence or absence of each protein under each SAR11 genome is presented in **Supplementary Table Tab 3**. In the UniProt section, absolute counts are presented instead of fractions, and therefore DnaA as the positive control is expected to have the highest numbers. References for protein functions in model organisms are provided in the rightmost column. **(*)** The UniProt search yielded a higher number of MinD proteins than DnaA, which can result from organisms having multiple MinD InterProScan ID search results. **(**)** Up to September 5th, 2025, TipN did not have an Interprocan ID, nor was registered in any protein family databases on UniProt. Therefore, the number reported for TipN in the SAR11 pangenomic analysis was obtained from blastp searches **(Supplementary Table Tab 15)** and in UniProt search by identifying similar protein sequences (50% identity cutoff) **(Supplementary Table Tab 16)**.

**3. Marker genes SAR11 panG|** Presence or absence of each cell cycle marker gene (**Fig. 1**, **Supplementary Table Tab 1**) in each SAR11 genome using a previously generated pangenome analysis[2]. The order of the genomes is the same as in the phylogenomic tree of the reference study[2].

**4-14|** Detailed bacterial taxonomic distribution for each of the cell cycle marker genes in **Fig. 1 and Supplementary Table Tab 1**. Searches were completed on July 30th, 2025.

**15. TipN BLASTP|** Results of the BLASTP search of the *C. crescentus* TipN protein sequence against the total 551,103 amino acids sequences from 470 SAR11 genomes as well as the outgroup HIMB59 genome. 9 sequences were identified by the blastp search. As they were also identified in the InterProScan search, we also show the results of InterProScan annotation.

**16. TipN on UniProt|** Similar proteins to the *C. crescentus* TipN on UniProt. Searches were completed on September 1st, 2025.

**17. All media recipes|** Molar concentrations of the chemical components for all the media used in this study. The components that had varying concentrations in a given experiment are highlighted.

**18. CCM2 recipe|** Detailed medium recipe for our new optimized medium (CCM2) that facilitated HTCC1062 growth rates near 1 doubling/day and yields reaching $10^9$ cells/mL **(Extended Data Fig. 3d)**, including the molar and mass concentrations of all components, as well as the concentrated stock concentrations for some of the components.

**19-23|** Source data for **Fig. 2**, **Extended Data Fig. 3,6,8**.

**24. Fluorescence vs. Ploidy|** Data for the fluorescence signal and ploidy level estimation in **Extended Data Fig. 9**.

### Supplementary References


1. Carini, P., Steindler, L., Beszteri, S. & Giovannoni, S. J. Nutrient requirements for growth of the extreme oligotroph 'Candidatus Pelagibacter ubique' HTCC1062 on a defined medium. *ISME J.* **7**, 592–602 (2013).
2. Lanclos, V. C. *et al.* Ecophysiology and genomics of the brackish water adapted SAR11 subclade IIIa. *ISME J.* **17**, 620–629 (2023).
3. Messer, W. The bacterial replication initiator DnaA. DnaA and oriC, the bacterial mode to initiate DNA replication. *FEMS Microbiol. Rev.* **26**, 355–374 (2002).
4. Katayama, T., Ozaki, S., Keyamura, K. & Fujimitsu, K. Regulation of the replication cycle: conserved and diverse regulatory systems for DnaA and oriC. *Nat. Rev. Microbiol.* **8**, 163–170 (2010).
5. Fujikawa, N. *et al.* Structural and biochemical analyses of hemimethylated DNA binding by the SeqA protein. *Nucleic Acids Res.* **32**, 82–92 (2004).
6. Boye, E. & Løbner-Olesen, A. The role of dam methyltransferase in the control of DNA replication in E. coli. *Cell* **62**, 981–989 (1990).
7. Olsson, J., Dasgupta, S., Berg, O. G. & Nordström, K. Eclipse period without sequestration in Escherichia coli. *Mol. Microbiol.* **44**, 1429–1440 (2002).
8. Olsson, J. A., Nordström, K., Hjort, K. & Dasgupta, S. Eclipse-synchrony relationship in Escherichia coli strains with mutations affecting sequestration, initiation of replication and superhelicity of the bacterial chromosome. *J. Mol. Biol.* **334**, 919–931 (2003).
9. Blair, J. A. *et al.* Branched signal wiring of an essential bacterial cell-cycle phosphotransfer

protein. *Structure* **21**, 1590–1601 (2013).

10. Krupka, M., Sobrinos-Sanguino, M., Jiménez, M., Rivas, G. & Margolin, W. Escherichia coli ZipA organizes FtsZ polymers into dynamic ring-like protofilament structures. *MBio* **9**, (2018).

11. Hale, C. A. & de Boer, P. A. Direct binding of FtsZ to ZipA, an essential component of the septal ring structure that mediates cell division in E. coli. *Cell* **88**, 175–185 (1997).

12. Pichoff, S., Du, S. & Lutkenhaus, J. Roles of FtsEX in cell division. *Res. Microbiol.* **170**, 374–380 (2019).

13. Meier, E. L. *et al.* FtsEX-mediated regulation of the final stages of cell division reveals morphogenetic plasticity in Caulobacter crescentus. *PLoS Genet.* **13**, e1006999 (2017).

14. Levin, P. A., Shim, J. J. & Grossman, A. D. Effect of minCD on FtsZ ring position and polar septation in Bacillus subtilis. *J. Bacteriol.* **180**, 6048–6051 (1998).

15. Machado, L. E. S. F. *et al.* NMR study of the interaction between MinC and FtsZ and modeling of the FtsZ:MinC complex. *J. Biol. Chem.* **301**, 108169 (2025).

16. Nußbaum, P. *et al.* An oscillating MinD protein determines the cellular positioning of the motility machinery in Archaea. *Curr. Biol.* **30**, 4956–4972.e4 (2020).

17. Dubarry, N., Willis, C. R., Ball, G., Lesterlin, C. & Armitage, J. P. In Vivo Imaging of the Segregation of the 2 Chromosomes and the Cell Division Proteins of Rhodobacter sphaeroides Reveals an Unexpected Role for MipZ. *MBio* **10**, (2019).

18. Corrales-Guerrero, L. *et al.* MipZ caps the plus-end of FtsZ polymers to promote their rapid disassembly. *Proc. Natl. Acad. Sci. U. S. A.* **119**, e2208227119 (2022).

19. Sogues, A. *et al.* Essential dynamic interdependence of FtsZ and SepF for Z-ring and septum formation in Corynebacterium glutamicum. *Nat. Commun.* **11**, 1641 (2020).

20. Pelletier, J. F. *et al.* Genetic requirements for cell division in a genomically minimal cell. *Cell* **184**, 2430–2440.e16 (2021).

21. Letzkus, M., Trela, C. & Mera, P. E. Three factors ParA, TipN, and DnaA-mediated

chromosome replication initiation are contributors of centromere segregation in Caulobacter crescentus. *Mol. Biol. Cell* **35**, ar68 (2024).

22. Cooper, S. & Helmstetter, C. E. Chromosome replication and the division cycle of Escherichia coli B/r. *J. Mol. Biol.* **31**, 519–540 (1968).
23. Skarstad, K., Steen, H. B. & Boye, E. Escherichia coli DNA distributions measured by flow cytometry and compared with theoretical computer simulations. *J. Bacteriol.* **163**, 661–668 (1985).
24. Stokke, C., Flåtten, I. & Skarstad, K. An easy-to-use simulation program demonstrates variations in bacterial cell cycle parameters depending on medium and temperature. *PLoS One* **7**, e30981 (2012).
25. Stokke, C., Waldminghaus, T. & Skarstad, K. Replication patterns and organization of replication forks in Vibrio cholerae. *Microbiology* **157**, 695–708 (2011).
26. Willis, L. & Huang, K. C. Sizing up the bacterial cell cycle. *Nat. Rev. Microbiol.* **15**, 606–620 (2017).
27. Boye, E. & Løbner-Olesen, A. Bacterial growth control studied by flow cytometry. *Res. Microbiol.* **142**, 131–135 (1991).
28. Wang, J. D. & Levin, P. A. Metabolism, cell growth and the bacterial cell cycle. *Nat. Rev. Microbiol.* **7**, 822–827 (2009).
29. Cheng, C. *Thrash-lab/SAR11_cell_cycle: V1.0.0*. (Zenodo, 2025). doi:10.5281/ZENODO.17703344.
30. Powell, E. O. Growth rate and generation time of bacteria, with special reference to continuous culture. *J. Gen. Microbiol.* **15**, 492–511 (1956).
31. Løbner-Olesen, A., Skarstad, K., Hansen, F. G., von Meyenburg, K. & Boye, E. The DnaA protein determines the initiation mass of Escherichia coli K-12. *Cell* **57**, 881–889 (1989).
32. Bremer, H. & Churchward, G. Deoxyribonucleic acid synthesis after inhibition of initiation of rounds of replication in Escherichia coli B/r. *J. Bacteriol.* **130**, 692–697 (1977).

33. Zheng, H. *et al.* General quantitative relations linking cell growth and the cell cycle in Escherichia coli. *Nat Microbiol* (2020) doi:10.1038/s41564-020-0717-x.

34. Müller, T., Walter, B., Wirtz, A. & Burkovski, A. Ammonium toxicity in bacteria. *Curr. Microbiol.* **52**, 400–406 (2006).

35. Heinrich, K., Leslie, D. J., Morlock, M., Bertilsson, S. & Jonas, K. Molecular basis and ecological relevance of *Caulobacter* cell filamentation in freshwater habitats. *MBio* **10**, (2019).

36. Hishinuma, F., Izaki, K. & Takahashi, H. Effects of Glycine andd-amino acids on growth of various microorganisms. *Agric. Biol. Chem.* **33**, 1577–1586 (1969).

37. Guo, X. *et al.* Automated determination of ammonium at nanomolar levels in seawater by coupling lab-in-syringe with highly sensitive light-emitting-diode-induced fluorescence detection. *Molecules* **30**, (2025).

38. Wedyan, M. *et al.* The biogeochemistry of dissolved amino acids in the aquatic ecosystems. *Mar. Environ. Res.* **210**, 107264 (2025).

39. Moran, M. A. *et al.* The Ocean's labile DOC supply chain. *Limnol. Oceanogr.* **67**, 1007–1021 (2022).

40. Seymour, J. R., Amin, S. A., Raina, J.-B. & Stocker, R. Zooming in on the phycosphere: the ecological interface for phytoplankton-bacteria relationships. *Nat Microbiol* **2**, 17065 (2017).

41. Rappé, M. S., Connon, S. A., Vergin, K. L. & Giovannoni, S. J. Cultivation of the ubiquitous SAR11 marine bacterioplankton clade. *Nature* **418**, 630–633 (2002).

42. Fu, H., Uchimiya, M., Gore, J. & Moran, M. A. Ecological drivers of bacterial community assembly in synthetic phycospheres. *Proc. Natl. Acad. Sci. U. S. A.* **117**, 3656–3662 (2020).

43. White, A. E., Giovannoni, S. J., Zhao, Y., Vergin, K. & Carlson, C. A. Elemental content and stoichiometry of SAR11 chemoheterotrophic marine bacteria: SAR11 composition. *Limnol. Oceanogr.* **4**, 44–51 (2019).

44. Malmstrom, R. R., Kiene, R. P., Cottrell, M. T. & Kirchman, D. L. Contribution of SAR11 bacteria to dissolved dimethylsulfoniopropionate and amino acid uptake in the North Atlantic ocean. *Appl. Environ. Microbiol.* **70**, 4129–4135 (2004).
45. Munson-McGee, J. H. *et al.* Decoupling of respiration rates and abundance in marine prokaryoplankton. *Nature* **612**, 764–770 (2022).
46. McDonald, M. D. *et al.* What is microbial dormancy? *Trends Microbiol.* **32**, 142–150 (2024).
47. Vergin, K. L. *et al.* High intraspecific recombination rate in a native population of Candidatus pelagibacter ubique (SAR11). *Environ. Microbiol.* **9**, 2430–2440 (2007).
48. López-Pérez, M., Haro-Moreno, J. M., Coutinho, F. H., Martinez-Garcia, M. & Rodriguez-Valera, F. The Evolutionary Success of the Marine Bacterium SAR11 Analyzed through a Metagenomic Perspective. *mSystems* **5**, (2020).
49. Overballe-Petersen, S. *et al.* Bacterial natural transformation by highly fragmented and damaged DNA. *Proc. Natl. Acad. Sci. U. S. A.* **110**, 19860–19865 (2013).
50. Mell, J. C. & Redfield, R. J. Natural competence and the evolution of DNA uptake specificity. *J. Bacteriol.* **196**, 1471–1483 (2014).
51. Pearson, G. D., Woods, A., Chiang, S. L. & Mekalanos, J. J. CTX genetic element encodes a site-specific recombination system and an intestinal colonization factor. *Proc. Natl. Acad. Sci. U. S. A.* **90**, 3750–3754 (1993).

### Supplementary Figure 1

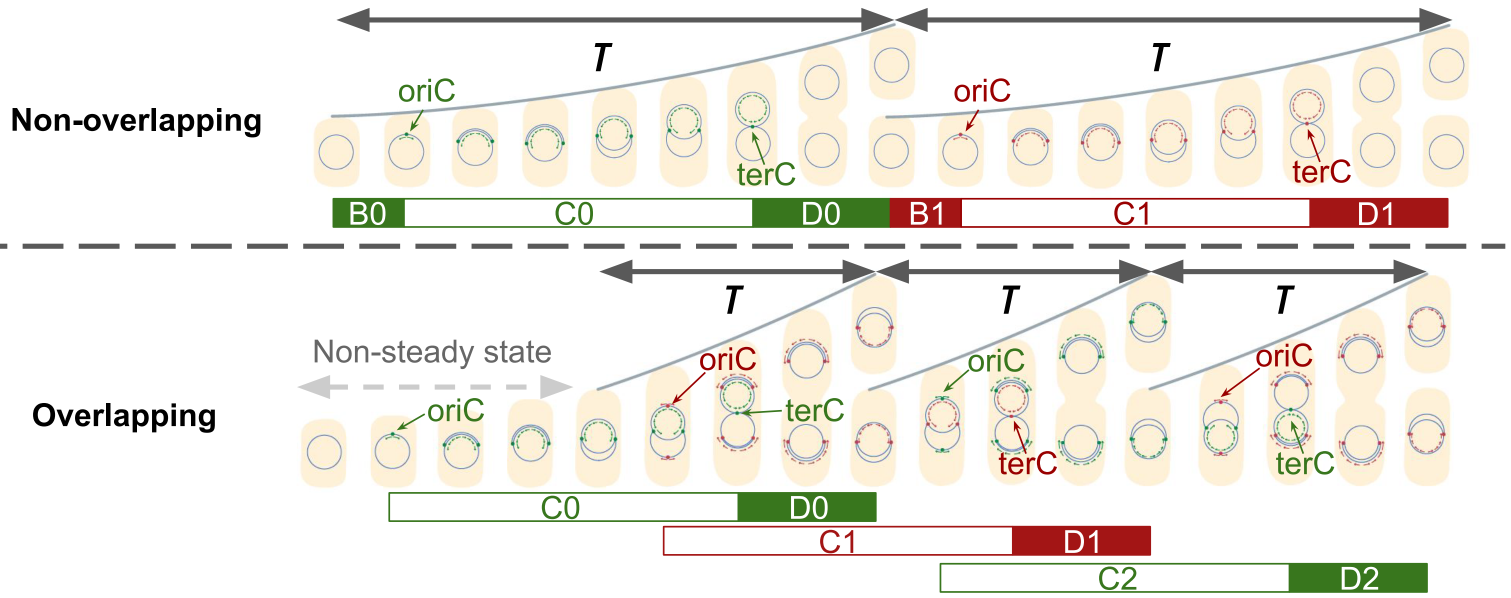

### Supplementary Figure 2

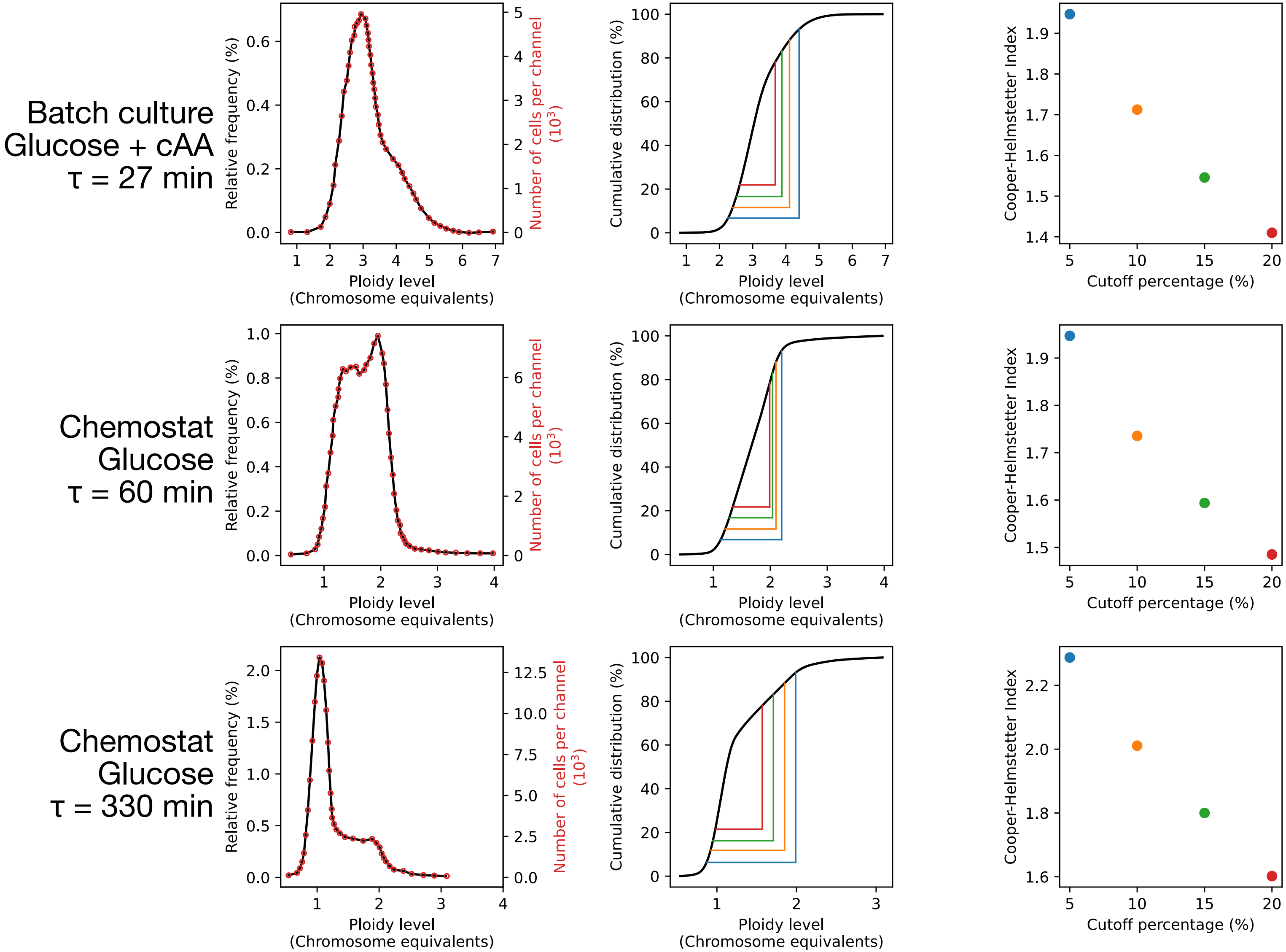

### Supplementary Figure 3

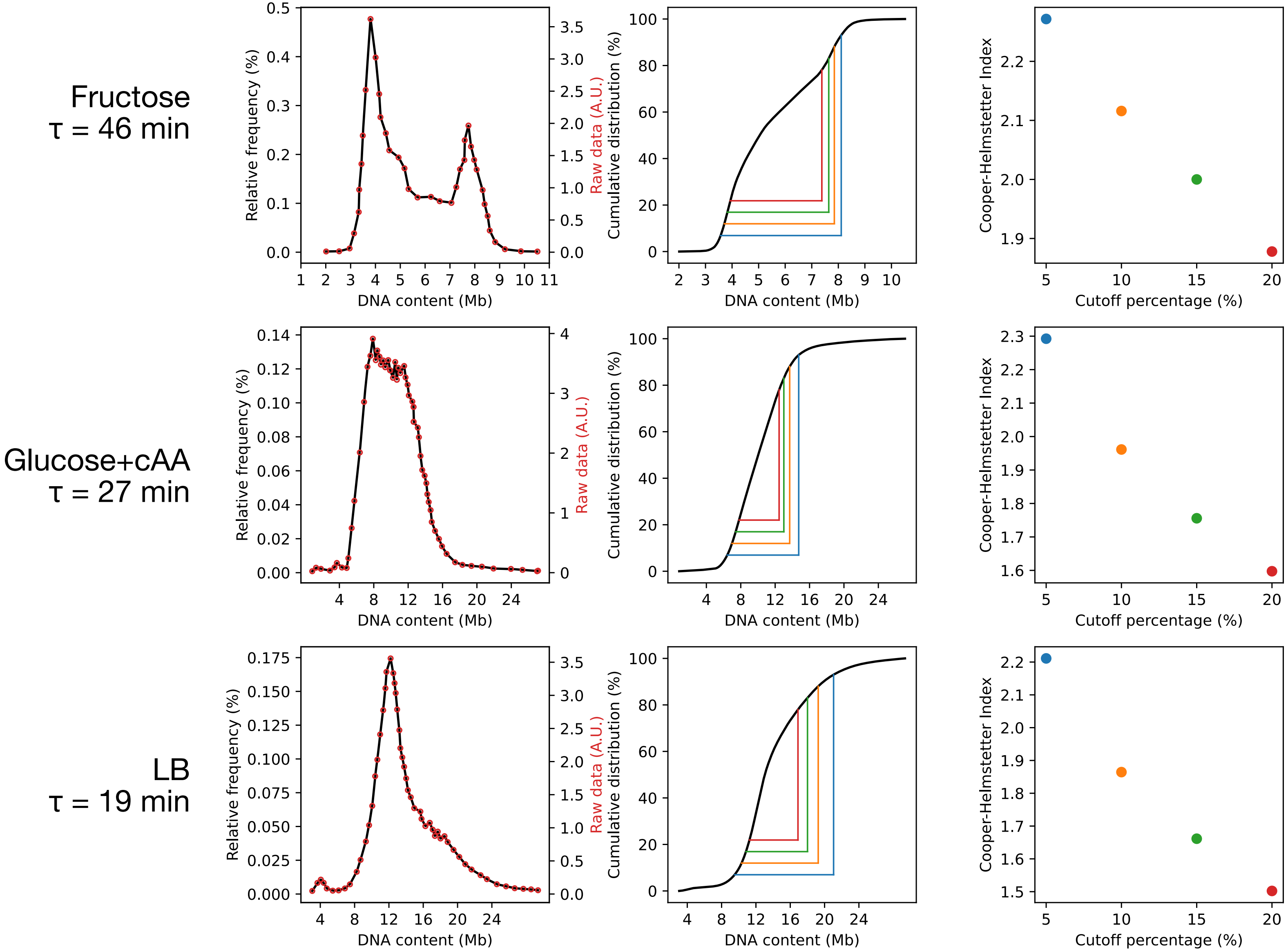

### Supplementary Figure 4

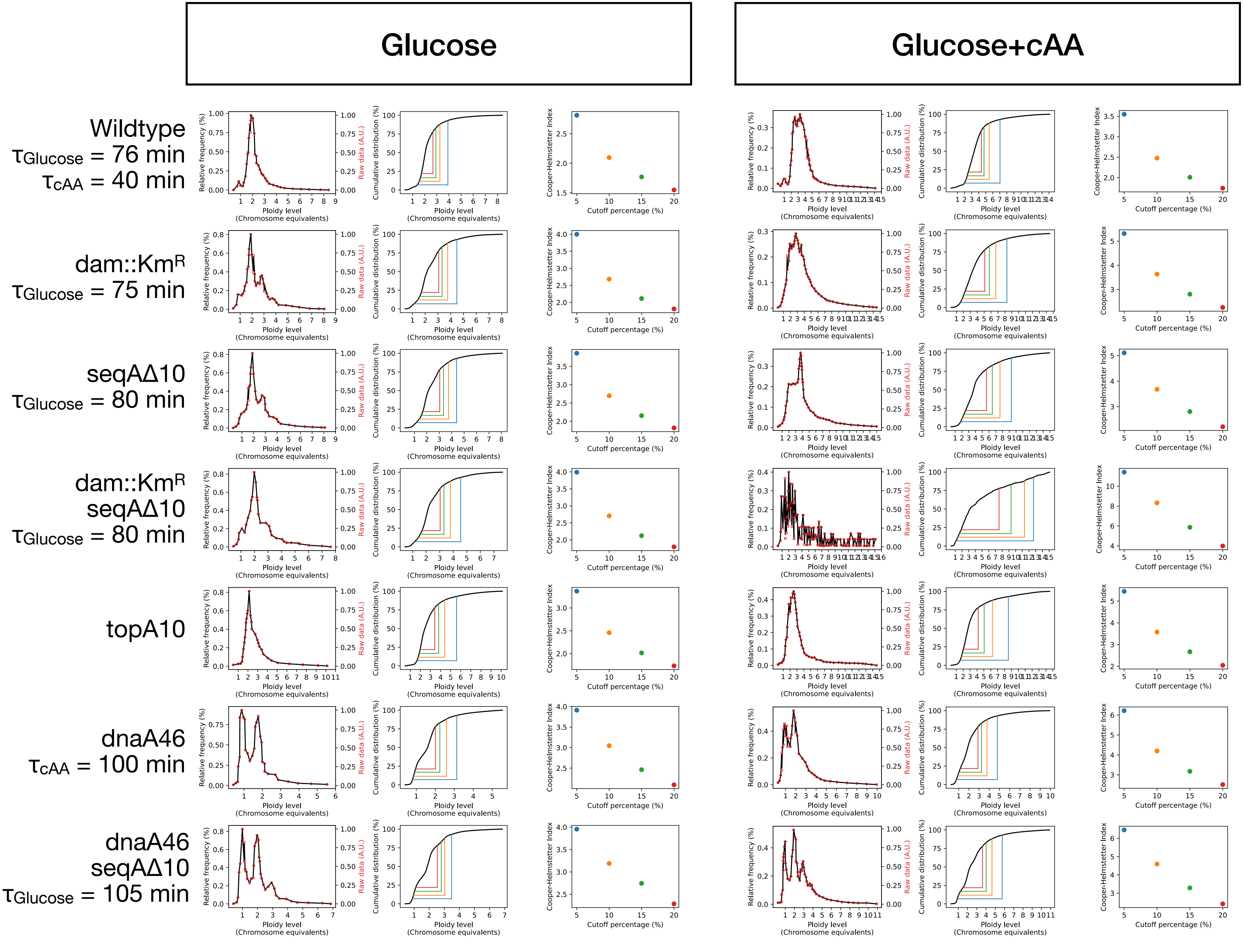

### Supplementary Figure 5

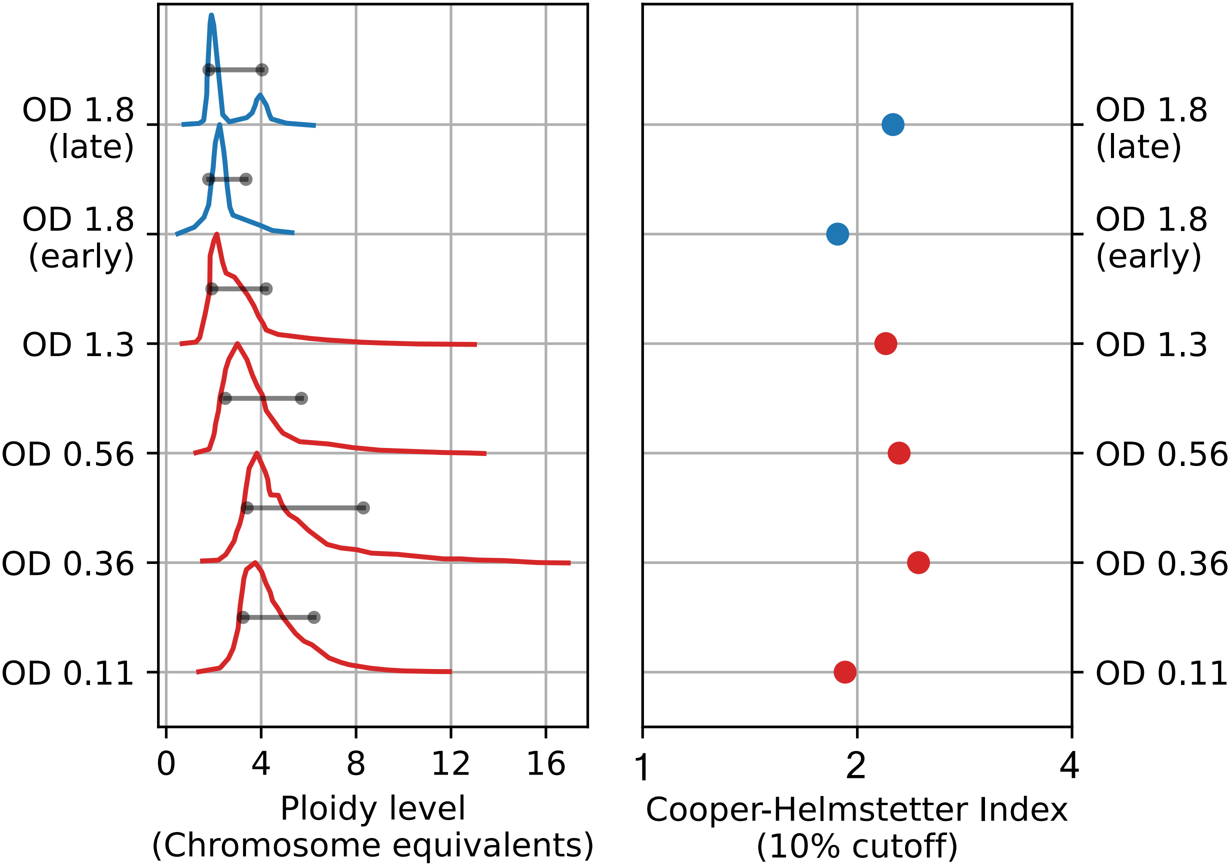

### Supplementary Figure 6

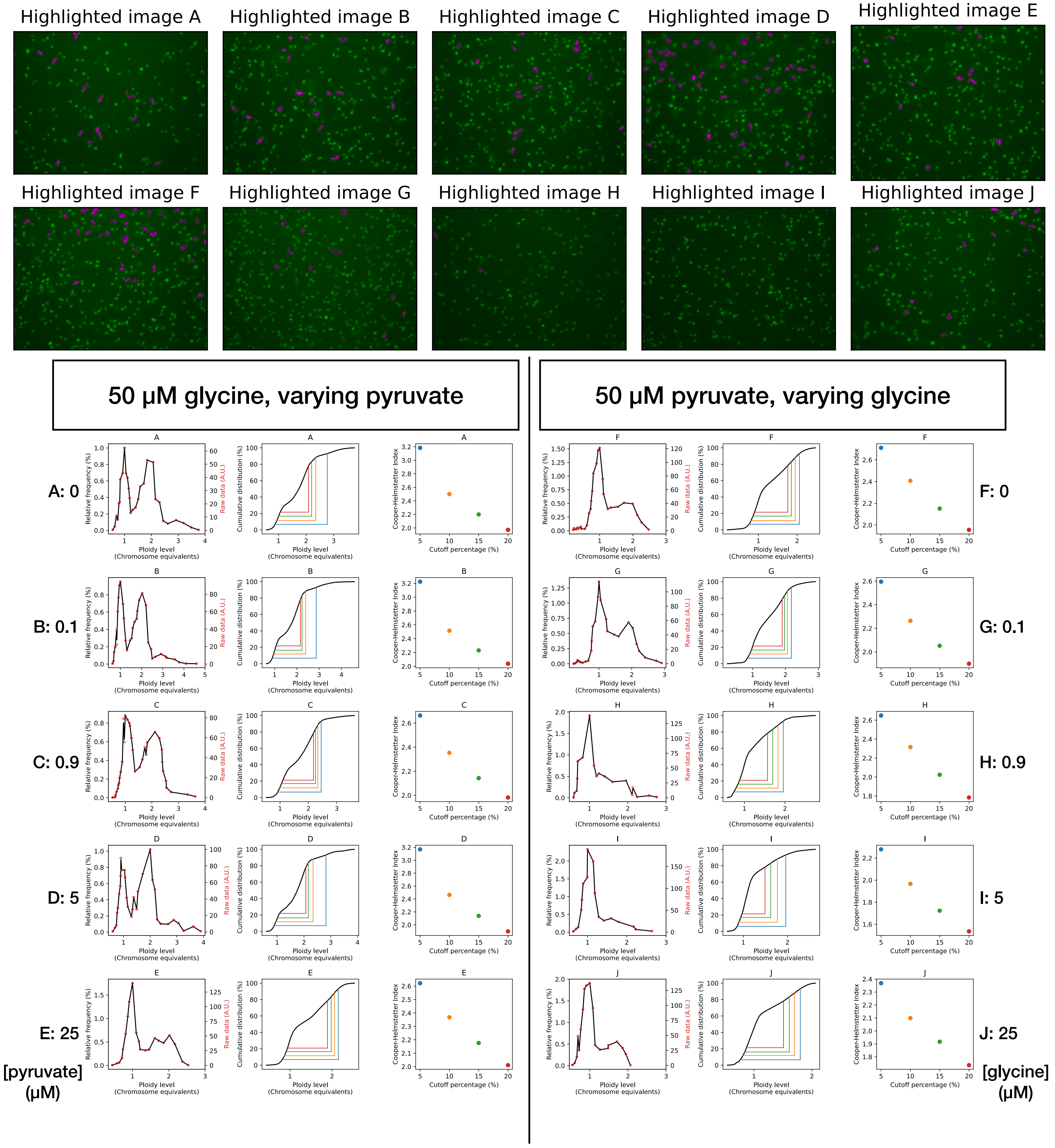

### Supplementary Figure 7

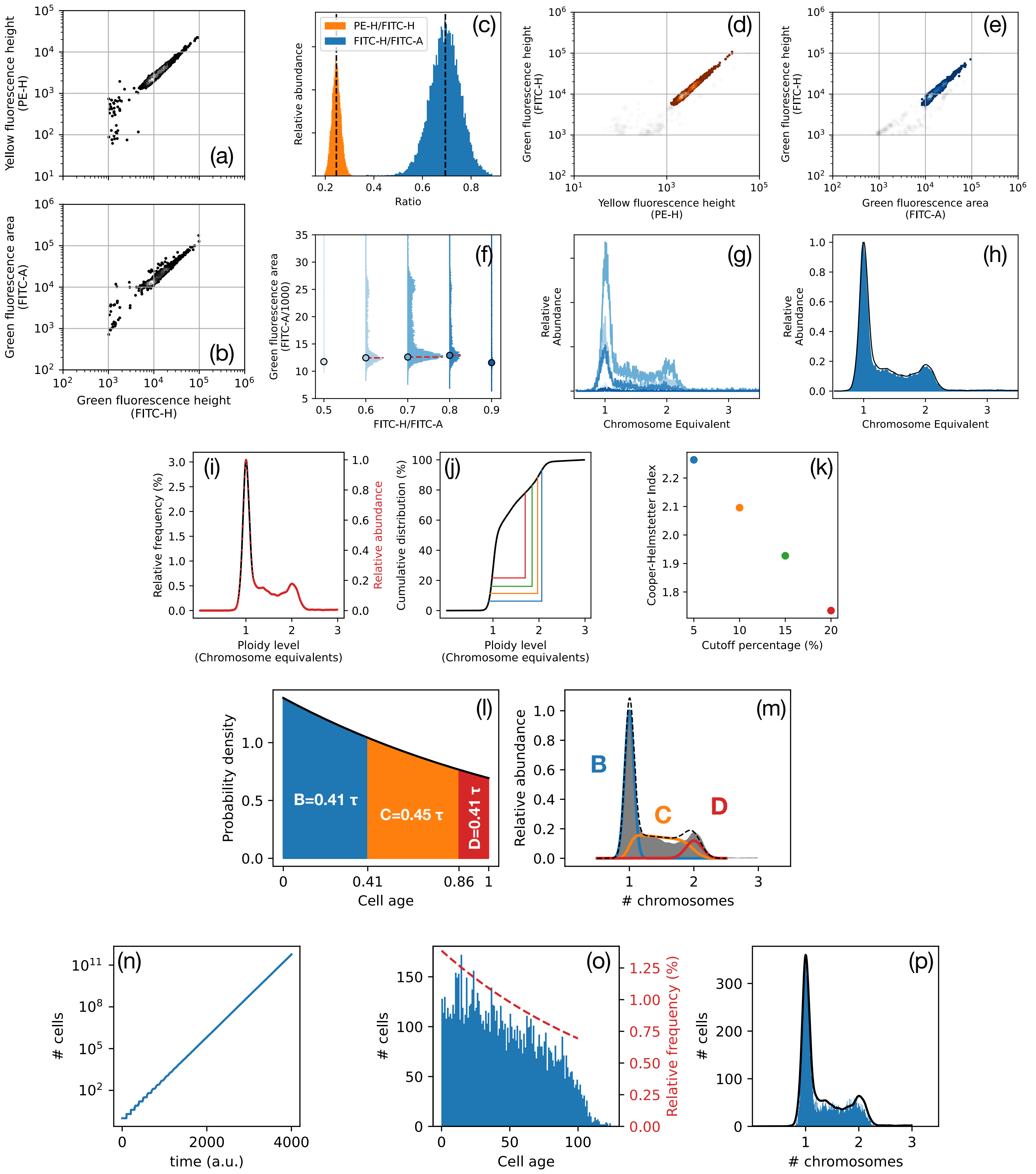

### Supplementary Figure 8

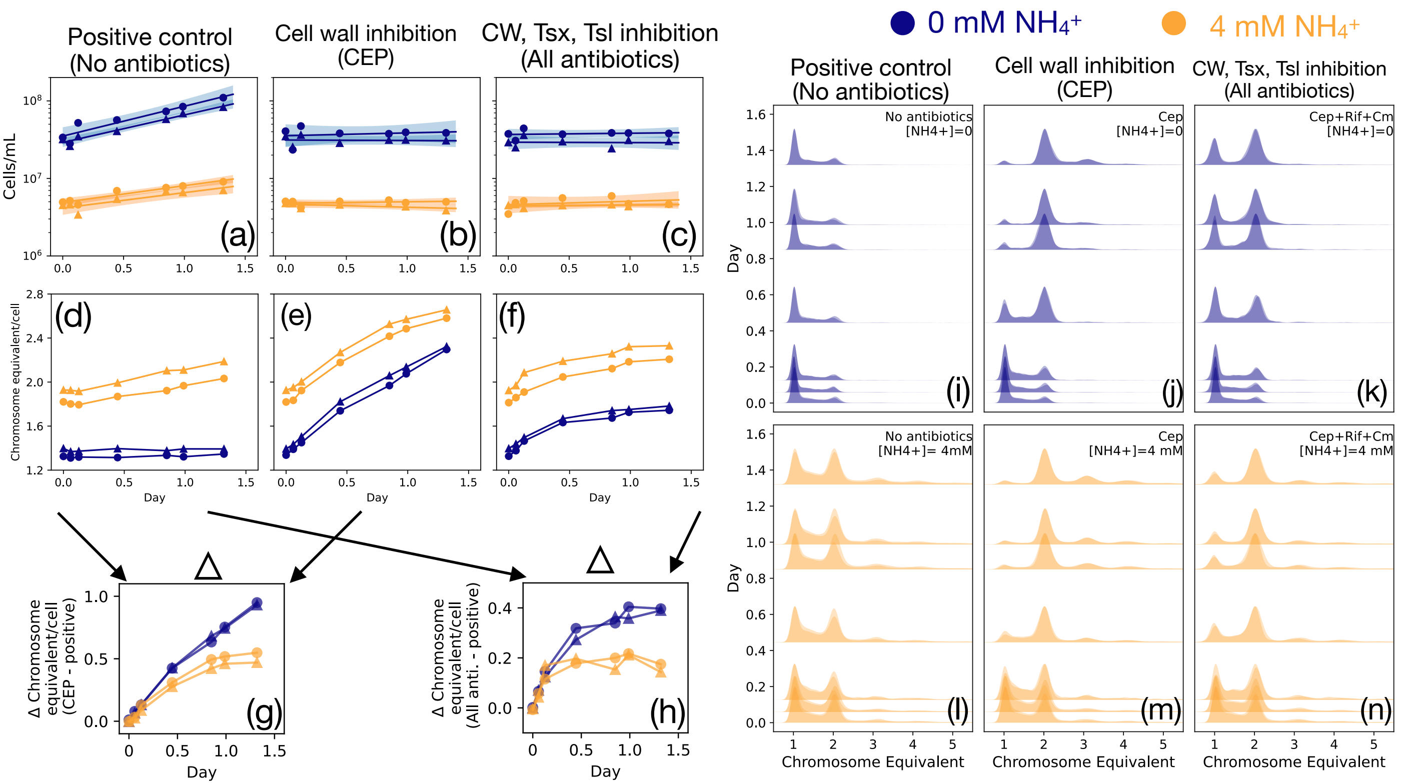
